# Fast adaptive super-resolution lattice light-sheet microscopy for rapid, long-term, near-isotropic subcellular imaging

**DOI:** 10.1101/2024.05.09.593386

**Authors:** Chang Qiao, Ziwei Li, Zongfa Wang, Yuhuan Lin, Chong Liu, Siwei Zhang, Yong Liu, Yun Feng, Xinyu Wang, Xue Dong, Jiabao Guo, Tao Jiang, Qinghua Wang, Qionghai Dai, Dong Li

## Abstract

Lattice light-sheet microscopy (LLSM) provides a crucial observation window into intra- and inter-cellular physiology of living specimens with high speed and low phototoxicity, however, at the diffraction-limited resolution or anisotropic super-resolution with structured illumination. Here we present the meta-learning-empowered reflective lattice light-sheet virtual structured illumination microscopy (Meta-rLLS-VSIM), which instantly upgrades LLSM to a near-isotropic super resolution of ∼120-nm laterally and ∼160-nm axially, more than twofold improvement in each dimension, without any modification of the optical system or sacrifice of other imaging metrics. Moreover, to alleviate the tremendous demands on training data and time necessitated by existing deep-learning (DL) methods, we devised an adaptive online training approach by synergizing the front-end imaging system and back-end meta-learning framework, which reduced the total time for data acquisition and model training down to tens of seconds. With this method, a new model can be well-trained with tenfold less data and three orders of magnitude less time than current standard supervised learning. We demonstrate the versatile functionalities of Meta-rLLS-VSIM by imaging a variety of bioprocesses with ultrahigh spatiotemporal resolution for long duration of hundreds of multi-color volumes, characterizing the dynamic regulation of contractile ring filaments during mitosis and the growth of pollen tubes, and delineating the nanoscale distributions, dispersion, and interaction pattern of multiple organelles in embryos and eukaryotic cells.

## Introduction

Elaborate bioprocesses occur in the three-dimensional (3D) space of living organisms in a complicated but organized way. Among all volumetric fluorescence imaging techniques, light-sheet microscopy (LSM) or called selective plane illumination microscopy (SPIM) stands out owing to its high spatiotemporal resolution and gentle 3D imaging capacity^1,2^. By using a second illumination objective placed perpendicular to the detection path, LSM confines the excitation within a micron-level thickness, which results in inherent optical sectioning and minimal out-of-focus excitation, massively reducing sample bleaching and phototoxicity compared to conventional epi-fluorescence imaging techniques, such as wide-field (WF) or confocal microscopy^3,4^. To extend the spatial resolution beyond the diffraction limit, several super-resolution (SR) techniques have been incorporated into LSM^5–8^, among which the lattice light-sheet structured illumination microscopy (LLS-SIM) achieves the optimal tradeoff between resolution and other equally important metrics for live cell imaging. Nonetheless, conventional LLS-SIM only permits a single orientation of structured illumination, so it suffers from anisotropic resolution and likely produces distortions along the orientations without resolution enhancement^8,9^. Therefore, current LSM is short of the full capability to accurately measure 3D subcellular morphology, biomolecular localization and signaling activity.

To overcome this issue, one recent study utilized oblique illumination coupled with a mechanical image rotation, which allows for three-orientation structured illumination (SI) and achieves laterally isotropic SR imaging^10^. The other latest work separated plane selection and SI pattern excitation by introducing reversibly photo-switchable fluorescent proteins, enabling higher excitation numerical aperture (NA) and spatial resolution than the oblique plane configuration^11^. Nevertheless, both solutions rely on repetitive excitation and acquisitions, i.e., thousands of raw images per volume, to reconstruct the final SR results, limiting the four-dimensional (4D) live imaging duration to no more than 50 timepoints^10^. Moreover, these methods only focus on addressing the anisotropy of lateral resolution, leaving the poor axial resolution as an outstanding problem.

In addition to advances in microscope hardware, computational approaches, especially deep learning-based methods, have brought a transformative impact on fluorescence microscopy. By learning the statistical inverse function of image transformation processes, deep neural networks (DNNs) have been applied to enhance both lateral^12–14^ and axial resolution^15–20^ of optical images. For instance, given the high-quality ground truth (GT) SR-SIM images as the training targets, DNNs are able to generate SR images directly from their diffraction-limited counterparts^13,14^. However, when applied to LLSM, such methods are subject to three major challenges: First, the isotropic GT-SIM data is unobtainable by conventional LLS-SIM, thus hindering from directly training an end-to-end network as previous works did; Second, improving the axial resolution from the diffraction limit (∼400 nm) to the lateral resolution of LLS-SIM (∼150 nm) simply by data-driven self-learning^12,17–20^ suffers from severe ill-posedness, leading to a significant risk of phantasmal generation^13,21^; Third, most DNN-based methods need to train a specific model for each biological specimen with a large amount of high-quality training data, consuming plenty of time spanning from several hours to a few days^22^. Consequently, these limitations in both optics and algorithms substantially impede a complete understanding of various animate bioprocesses, which requires high-resolution imaging across all four dimensions of space and time simultaneously.

In this study, we present the meta-learning-empowered reflective lattice light-sheet virtual structured illumination microscopy (Meta-rLLS-VSIM) as well as an assortative dual-stage near-isotropic SR reconstruction framework. We demonstrate the proposed method can extend the single-dimensional (1D) SR capability of LLS-SIM to all three dimensions, resulting in a near-isotropic super resolution of 120-nm in lateral and 160-nm in axial, without any modifications of the optical system or sacrifice of other imaging metrics compared to LLSM. In practical implementation, a DNN model for 1D SR inference is first trained using data acquired with the SI mode of the LLS-SIM system. Distinct from existing standard supervised learning, we devise a training scheme based on meta-learning^23^, which learns a comprehensive initial weight for fast adaptation to the scenarios of new biospecimens or signal-to-noise ratios (SNRs) by means of minimal training data. By synergizing the automatic data acquisition workflow and back-end meta-learning algorithms, we show that a new DNN model can be well-trained within tens of seconds, three orders of magnitude faster than standard training procedure. Next, we exploit the structural similarity and orientational randomness of biological specimens within the lateral space to infer anisotropic SR intermediates in multiple orientations (dubbed as virtual structured illumination), which are then combined using a generalized Wiener filter approach to obtain laterally isotropic SR volumes. Finally, instead of violently improving axial resolution by self-learning and isotropic prediction^17,20,24^, we adopt a physically rationalized strategy of reflective imaging^25^ that helps us create an additional virtual view of the sample complementary to the real one in sheet-scanning. Different from previous Bayesian-based joint deconvolution methods^25,26^, we design a self-supervised Richardson-Lucy dual-cycle fusion network (RL-DFN) to fully exploit the complementary resolution information from both views and recover the final near-isotropic SR data volume. As such, we only need one-way sheet-scan acquisition of raw data to reconstruct near-isotropic SR volumes, which thus fulfills an unmet need for rapid, long-term, near-isotropic SR volumetric observation of subcellular dynamics with high fidelity and quantifiability. We demonstrate the versatile usability of Meta-LLS-VSIM on a large variety of specimens, including the botanic pollen tubes, thick mouse embryos, developmental *C. elegans* embryos, and other eukaryotes during mitosis or interkinesis.

## Results

### Laterally isotropic SR reconstruction by virtual structured illumination

The homebuilt LLSM/LLS-SIM system was developed from the original design^8^, which operates at two modes in our experiments by assigning different phase patterns on the spatial light modulator: the structured illumination (SI) mode for training data acquisition, and the sheet-scan mode for fast light-sheet imaging (**Fig. 1a** and Methods). We noted that the biological specimens always arranged or grew in random orientation, so the training dataset of paired LLSM and LLS-SIM images as a whole contains complete SR information about the specimens, even each of which just carries SR information of a single dimension. Accordingly, the trained DNN model actually carries complete feature maps of producing anisotropic SR intermediates for any given single dimension. Therefore, anisotropic 1D SR images along any orientation can be produced by simply rotating the input image relative to the original input data, and then reapplying the trained DNN on the rotated images^27,28^. Similar to the reconstruction in standard SIM, multiple 1D SR stacks along equally spaced orientation angles were combined using a generalized Wiener filter approach to obtain laterally isotropic SR volumes. Therefore, we named this method as virtual structured illumination super-resolution (VSI-SR). Briefly, the VSI-SR scheme was implemented by three steps: (i) Acquire the training dataset using SI mode of the system and train a VSI-SR model (**Fig. 1b**); (ii) Apply the well-trained model onto the raw data that is rotated to three orientations equally spaced by 60°, hence generating three anisotropic SR components; and (iii) Combine the different components through joint deconvolution in the Fourier space (**Fig. 1c**). Of note, compared to the previous implementation^28^, the VSI-SR scheme possesses three advantages: First, we employed coherent structured illumination rather than incoherent photo-reassignment to obtain the GT-SIM training data, offering higher resolution improvement and acquisition efficiency (Methods); Second, we elaborately designed a multi-input/output generative adversarial network (GAN) architecture with a Fourier space-regularized loss function (Supplementary Note 1), yielding an optimal output with regard to both resolution enhancement and high fidelity of SR information (Extended Data Figs. 1-3 and Supplementary Fig. 1); Third, instead of calculating the max intensity projections (MIP) in frequency space^28^, we incorporated the prior knowledge of the deterministic optical transfer function (OTF) of each component into a generalized Wiener filter to combine the SR components at different orientations, resulting in higher contrast and fewer artifacts (Extended Data Fig. 4).

**Fig. 1.**
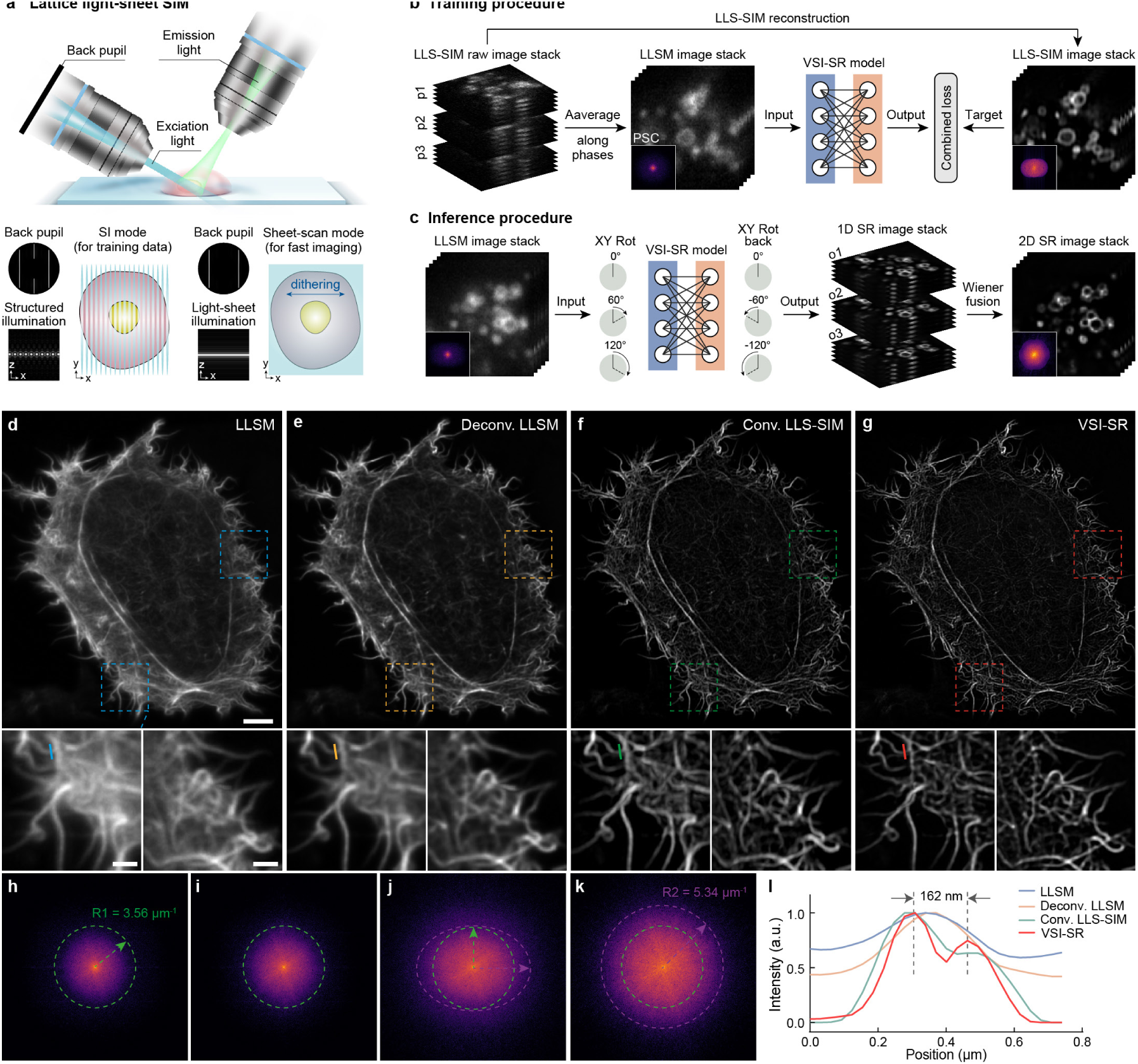
Laterally isotropic SR reconstruction by virtual structured illumination. **a**, Illustration of the LLS-SIM system (upper panel) and its two imaging modes: structured illumination mode (bottom left corner) and sheet-scan mode (bottom right corner), with sketches of the back pupil plane and illumination in x-z and x-y views shown for each mode. **b**, Schematic of the training processes of the 2D VSI-SR network. We trained the VSI-SR model with the diffraction-limited LLSM images as input and LLS-SIM images as targets to learn the mapping of resolution improvement along the illumination orientation. **c**, Schematic of the 2D isotropic reconstruction process with the trained VSR-SR model. **d-g**, Representative maximum intensity projections (MIPs) of F-actin image stacks obtained by LLSM (d) and processed by RL deconvolution (e), conventional LLS-SIM (f), and VSI-SR (g). **h**-**k**, Logarithmic power spectrum coverages (PSCs) of the MIP images shown in d-g. **l**, Comparison of the intensity profile plots of LLSM (blue), deconvolved LLSM (yellow), Conv. LLS-SIM (green), and VSI-SR (red) along the short lines labelled in d-g. Scale bar, 5 μm (d), 2 μm (zoom-in regions of d).

To characterize the performance of the VSI-SR method, we examined it on two different SIM systems, LLS-SIM (Fig. 1d-l and Supplementary Fig. 2) and total internal reflective fluorescence (TIRF) SIM (Supplementary Fig. 3), with a wide variety of subcellular structures including clathrin-coated pits (CCPs), microtubules (MTs), endoplasmic reticulum (ER), and F-actin filaments. We noted that although the OTF support could be extended along lateral *x*-axis by conventional LLS-SIM with respect to the original or deconvolved LLSM images, the diffraction-limited resolution along *y*-axis caused a bottleneck to clearly resolve the fine structures of threadlike F-actin, punctate CCPs, tubular MTs, and reticulated ER in close proximity (Fig. 1f and Supplementary Fig. 2). In contrast, VSI-SR reconstructions successfully circumvented this problem by replenishing the high-frequency information in all lateral dimensions (Fig. 1h-k), which permitted resolving the dense actin filaments crisscrossing each other (Fig. 1l). This high-fidelity resolution improvement by VSI-SR was further validated with TIRF-SIM system and cross-modality testing, where the isotropic TIRF-SIM images could serve as the GT references (Supplementary Figs. 3, 4, and Supplementary Note 2). These results illustrate the effectiveness and robustness of the proposed VSI-SR strategy for reconstructing laterally isotropic SR images from diffraction-limited lattice light-sheet raw images.

### Meta-learning-based fast model adaptation for diverse biological specimens

Due to the extreme diversity of subcellular biological structures and limited representation ability of DNNs, current deep learning-based SR methods usually need to train a dedicated model for each specific biological structure in order to ensure optimal inference performance. However, the training process for each model often necessitates a large amount of high-quality data acquired from more than 30 distinct regions of interest (ROIs), and takes very long time of several hours to a few days^13,14,28^, which considerably impedes the applicability and efficiency of using such methods in daily experiments. On the other hand, we noticed that meta-learning (or called learning-to-learn) has been attracting growing attention of computer vison community in recent years^29^. Instead of just learning task-specific knowledge, meta-learning aims to capture the commonality of different tasks and find a sensitive and transferable point in the parameter space where the trained meta-model can quickly adapt to a new task with a small number of gradient updates and minimal data. Inspired by the model-agnostic meta-learning (MAML) algorithm^23^, we devised a meta-learning framework for the VSI-SR model (Meta-VSI-SR), and equipped our LLSM system with the meta-learning-empowered fast adaptation capability (Meta-LLS-VSIM) by streamlining automatic data acquisition, pre-processing, and meta-finetuning procedures (**Fig. 2a-d** and Supplementary Note 3). Specifically, a meta-model (or meta-learner) was first trained with a large pre-acquired dataset of 10 distinct biological structures at two excitation intensity levels for each structure, which learned structure- and SNR-independent general knowledge of the SR task (**Fig. 2a**, Methods, and Extended Data Fig. 5). It is noteworthy that the pre-trained meta-model is not aimed to be directly used for image processing, but is capable of fast adapting to the unseen scenario of specific structures or SNRs. Once the meta-model is well-trained by experts, the adaptation procedure is very simple and friendly for end users: (i) firstly selecting three ROIs in the wide-field overview window of the control software; (ii) then the data acquisition and all computational procedures including LLS-SIM reconstruction, data augmentation, and meta-finetuning will be automatically executed (**Fig. 2b** and Methods). We demonstrated that the finetuning process could be accomplished in as fast as 30 seconds with 3 ROIs, i.e., 720-fold faster and 12-fold less data than standard training, respectively, while yielding over 6dB improvement from the starting point in peak signal-to-noise ratio (PSNR) and superb SR performance (**Fig. 2c**, Supplementary Video 1). In addition, distinct from most deep learning SR methods that work in 2D imaging scenarios, Meta-VSI-SR models aimed to optimize the volumetric SR reconstruction capability by adopting a custom-designed multi-slice input/output scheme and a Fourier space-regularized discriminative loss function, which permitted the Meta-VSI-SR network to effectively utilize the structural continuity along the axial axis, and efficiently discriminated the high frequency components in Fourier space (Extended Data Figs. 1-3 and Supplementary Note 1).

**Fig. 2.**
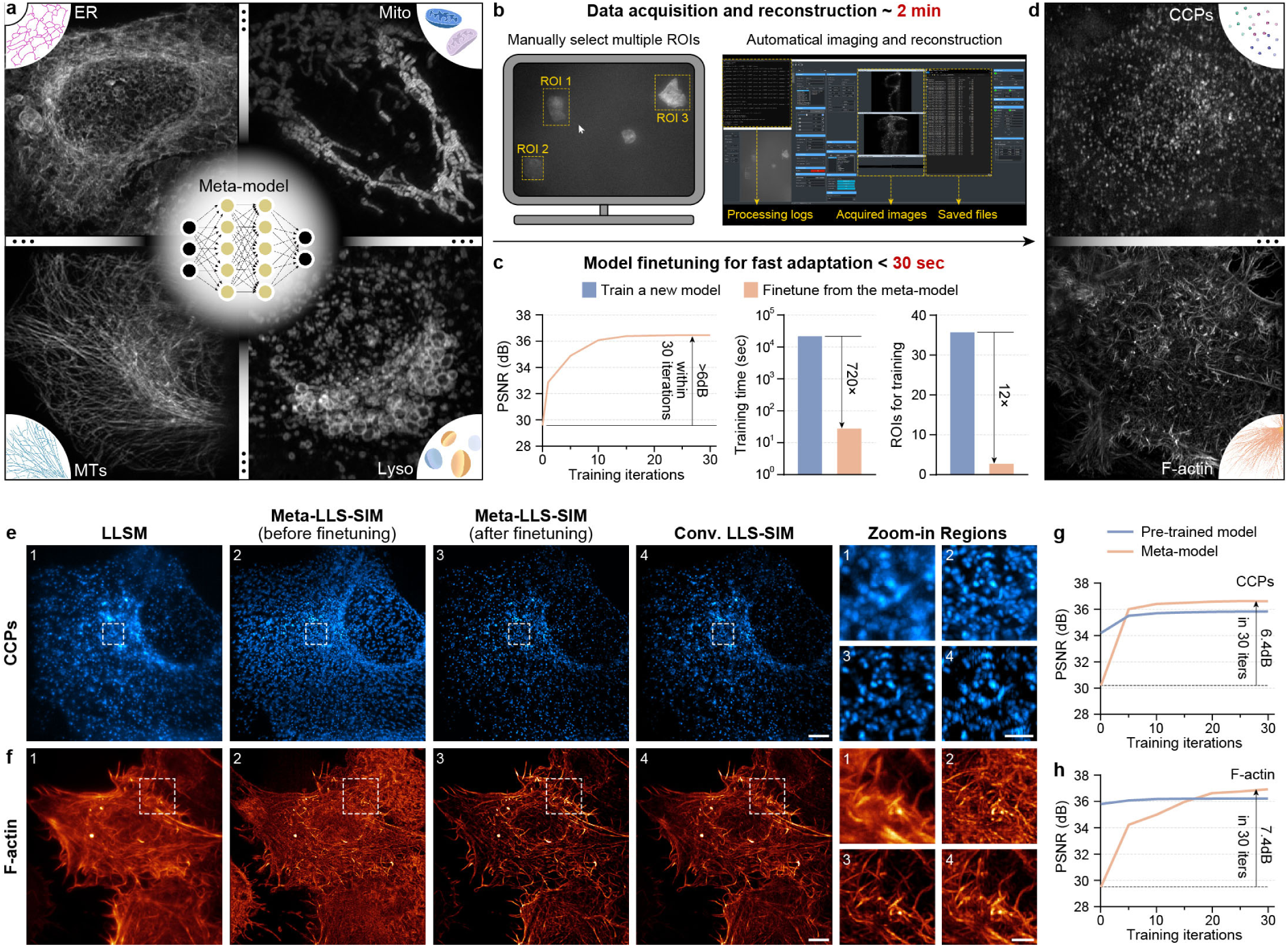
Meta-LLS-VSIM with laterally isotropic resolution and fast adaptation capability. **a-d**, The pipeline of Meta-LLS-VSIM imaging. A well-trained meta-model (a) can fast adapt to new biological structures (d) by a few simple steps: manually selecting three ROIs in the overview window of the operation interface (left part of b), waiting for automatic imaging, data pre-processing (right part of b), and meta-finetuning (c). The overall time for ROI selection, data acquisition, reconstruction and model finetuning is often less than 2.5 minutes (Supplementary Video 1). **e**, **f**, Representative max intensity projections (MIPs) of CCPs (e) and F-actin (f) imaged by LLSM (first column), Meta-LLS-VSIM before (second column) and after (third column) finetuning, and conventional LLS-SIM (fourth column). The right panel shows the magnified images of the boxed regions in the left images. Scale bar, 5 μm (e, f), 1.5 μm (zoom-in regions of e), 2 μm (zoom-in regions of f). **g**, **h**, Reconstruction PSNR progressions of meta-models and pre-trained models during finetuning procedures for CCPs (g) and F-actin (h) images.

Next, we performed an evaluation of Meta-VSI-SR methods by processing two new structures of CCPs and F-actin filaments which were not included in the meta-training dataset (**Fig. 2d**). We found that although the original meta-model without finetuning generated massive background artifacts, after meta-finetuning, the SR performance was instantly boosted by a large margin for both structures (**Fig. 2e, f**). In particular, Meta-VSI-SR eliminated the reconstruction artifacts in anisotropic LLS-SIM images, thereby clearly resolving finer details of either CCP distribution or actin filament crisscrossing. Moreover, we compared the finetuning progression of meta-models and pre-trained models, i.e., the model trained with the standard gradient descent algorithm, both of which were trained using the same dataset. Compared to the pre-trained model, the meta-model evolved dramatically faster and converged to a higher PSNR with only 30 minibatch iterations during finetuning (**Fig. 2g, h**), indicating that simply employing transfer learning or finetuning from a pre-trained network does not yield superior SR performance or generalization capability similar to our Meta-VSI-SR scheme.

### 4D SR live-cell imaging with extended duration and enhanced optical sectioning capacity

Volumetric SR imaging usually requires much higher excitation light intensity and more acquisition time per volume than diffraction-limited 3D imaging^8,9^, thus its live imaging duration has been limited to ∼50 timepoints even for single color^10,28^. Resorting to the fast adaptation ability and superior SR performance of Meta-VSI-SR scheme, the Meta-LLS-VSIM is able to generate laterally isotropic multi-color SR images for a wide variety of specimens by operating at the sheet-scan mode and relatively low SNR condition. Hence, Meta-LLS-VSIM enables multi-color imaging of light-sensitive bioprocesses at unparalleled spatial and temporal resolution for a prolonged observation window. For instance, the F-actin contractile ring (CR) plays a vital role in generating the constricting force to segregate the two dividing cells during mitosis^30^, but whether it facilitates the partitioning of other organelles has rarely been studied. Here, we employed Meta-LLS-VSIM to image HeLa cells stably expressing Lifeact-mEmerald, KDEL-mCherry, and Lamp1-Halo for 357 timepoints (>190,000 slices in total) at 8 seconds per three-color whole cell volume, which recorded the entire process from CR formation, contraction to disassembly during mitosis (**Fig. 3a** and Supplementary Video 2). The enhanced resolution and volumetric imaging enabled accurate quantification of the contraction dynamics (**Fig. 3b**). We identified that the CR was contracted at a stable velocity in each individual cell, although varied from cell to cell (**Fig. 3c**, gray; Supplementary Fig. 5a). Moreover, the multi-color imaging allowed us to examine the coordination dynamics between CR contraction and the partitioning of membranous organelles. We observed that both the continuous ER and distributed lysosomes (Lyso) autonomously moved away from the cross-sectional area of the CR in advance of ring closing (**Fig. 3c**, magenta and yellow, and **Fig. 3d**), during which there are little interactions between the CR and ER or Lyso (Supplementary Fig. 5b-i). These observations together implied that the CR was not directly involved in ER fission or Lyso partitioning in cytokinesis.

**Fig. 3.**
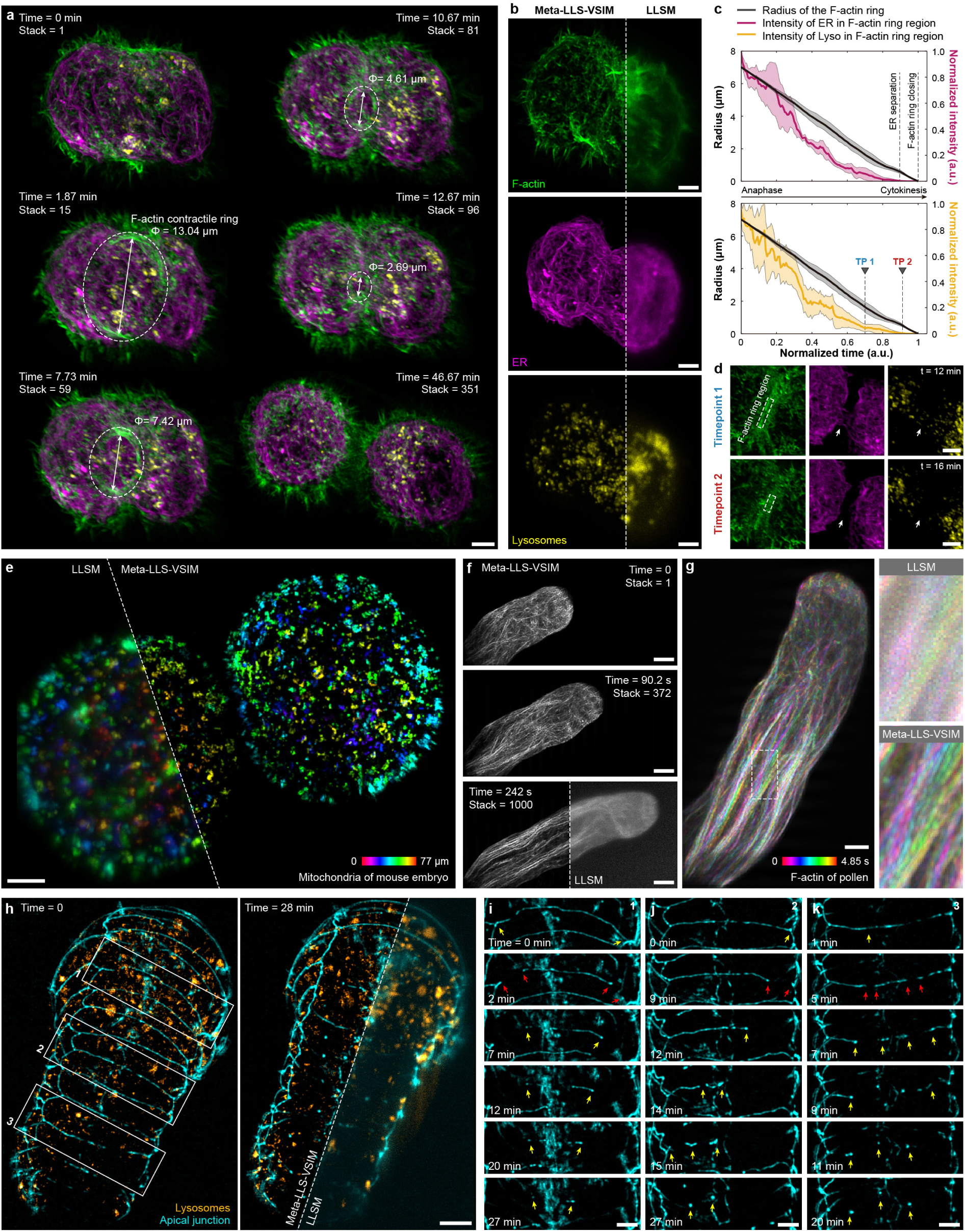
4D SR live imaging by Meta-LLS-VSIM. **a**, Three-color Meta-LLS-VSIM images of F-actin (green), ER (magenta), and Lyso (yellow) at different timepoints of mitosis, showing the spatiotemporal dynamics of the three organelles along with the constriction of the CR (Supplementary Video 2). **b**, Meta-LLS-VSIM and LLSM images of a representative timepoint, showing that Meta-LLS-VSIM resolves finer details of all three structures than conventional LLSM. **c**, Plots of the contractile ring radius (gray), total intensity of ER (magenta) and Lyso (yellow) in the cross-sectional area of the CR over the normalized time course. **d**, SR images of F-actin, ER, and Lyso at two representative timepoints marked in the lower panel of c, showing the spatial distribution of CR, ER, and Lyso from a view parallel to the CR plane. **e**, Representative Meta-LLS-VSIM image of a developing mouse embryo expressing TOMM20-mEmerald, which is 0color-coded for distance from the substrate (Supplementary Video 3). **f**, Time-lapse Meta-LLS-VSIM images of a growing pollen tube (Supplementary Video 4). **g**, Temporally color-coded images of the pollen tube, the boxed regions imaged by Meta-LLS-VSIM and conventional LLSM are magnified on the right for comparison. **h**, Two-color Meta-LLS-VSIM images of a *C. elegans* embryo labelled with apical junction and Lyso before (left) and after (right) seam cell fusion. LLSM image is shown in the rightmost corner for comparison. **i-k**, Three representative cases with different initiation fusion sites on apical junction between adjacent seam cells: fusions beginning from both ends (i), from a single end (j), and from several intermediates (k). These magnified images correspond to regions labelled by white boxes in h. Scale bar, 3 μm (a, b, d, g, and i-k), 8 μm (e), 6 μm (f), 5 μm (h).

The performance of conventional LLSM when imaging thick or scattering samples is limited because of the sample-induced scattering and light-sheet broadening at the distance far from the beam waist. Synchronizing the line illumination mode with rolling-shutter detection of an electronic scientific complementary metal-oxide-semiconductor (sCMOS) to achieve a partial confocal effect is able to enhance the optical sectioning capability^31^, which, however, is not compatible with the super-resolution LLS-SIM mode^8^. To extend the application scope of volumetric SR imaging to deep and scattering tissues, we incorporated the Meta-VSI-SR scheme with synchronized rolling-shutter confocal slit-scan. Within this strategy, the Meta-VSI-SR model learned SR capacity of LLS-SIM from shallow parts of the specimen, i.e., with little scattering-induced aberration, and then was applied to the whole sample captured with the rolling-shutter confocal slit-scan mode (Methods), thereby enabling laterally isotropic 4D SR imaging for thick and scattering samples. To illustrate the potential of this method, we first employed it to record the dynamics of mitochondria in a mouse embryo labelled with TOMM20-mEmerald across a thick area of 95×95×96 μm^3^ for 100 timepoints (**Fig. 3e**). The high-quality volumetric SR images enabled us to observe and track the active dynamics of mitochondria (Supplementary Video 3), and we noticed that unlike mature somatic cells, the mitochondria in early mouse embryo mostly present rounded or punctate morphology, which is consistent with the observation by electron microscopy^32^.

Next, we examined the optical sectioning capability of Meta-LLS-VSIM by imaging a growing pollen tube labelled with Lifeact-GFP (Methods) at a high speed of 4.125 Hz for 1,000 consecutive volumes (**Fig. 3f** and Supplementary Video 4). Despite being captured via the slit-scan mode, the raw images were heavily contaminated by the strong scattering of plant cytoplasm, and the fine details of F-actin could not be distinguished due to the poor spatial resolution. In contrast, the isotropic SR reconstruction of Meta-LLS-VSIM substantially improved both contrast and resolution, revealing the spatiotemporal dynamics of cytoskeleton during the growing of pollen tubes (**Fig. 3g**).

The plasma membrane fusion in the development of *C. elegans* has been long attracting interest of cell biologists, but was mostly studied with confocal or two-photon imaging deployed within a thin layer at an interval of several minutes because of the vulnerability of the developing embryo^33^. In virtue of the low phototoxicity and remarkable optical sectioning capacity of slit-scan Meta-LLS-VSIM, we could capture a two-color video that clearly records the whole plasma membrane fusion process during *C. elegans* embryo development with high spatiotemporal resolution (**Fig. 3h** and Supplementary Video 5), which is difficult for existing SR techniques. Interestingly, we observed three typical cases with different initiation fusion sites on the apical junction between adjacent seam cells, i.e., fusions beginning from both ends (**Fig. 3i**), from a single end (**Fig. 3j**), and from several intermediates (**Fig. 3k**), which suggested a multitudinous mechanism of seam cell fusion regulation.

### Near-isotropic SR reconstruction by Meta-rLLS-VSIM

Although the VSI-SR scheme can effectively extend the 1D SR capability to other lateral orientations, it cannot be applied to enhance the axial resolution. In recent years, several DL-based methods were developed to directly improve axial resolution by deep self-learning^12,17,19,20^ or cycle-consistent generative adversarial network^18^ in a reference-free manner. However, these kinds of techniques are subject to two major defects. First, the precondition for using them is that the sample structure itself has an isotropically morphological distribution in the 3D space, so that the SR knowledge learned from the x-y plane can be generalized into x-z or y-z planes. Unfortunately, this prerequisite is not always true for biological data, e.g., the majority of tubular structures extend laterally but not axially in adherent cells. Second, the axial resolution is usually three times worse than lateral, but improving both optical resolution and sampling rate by three times directly from diffraction-limited acquisitions suffers from severe ill-posedness, which substantially degrades output fidelity^13^ (Extended Data Fig. 6).

To rationally enhance axial resolution of Meta-LLS-VSIM, we modified our live imaging configuration by replacing the transparent coverslip with a reflective one, which allowed us to simultaneously detect two symmetrical views of the sample^25^ (**Fig. 4a** and Methods). Instead of deploying the naive multi-view deconvolution^25,26,34^ for view fusion, we devised a self-supervised dual-view fusion algorithm, dubbed Richardson-Lucy dual-cycle fusion network (RL-DFN), that incorporated the multi-view Richardson-Lucy (RL) iteration and deterministic point spread function (PSF) priors into the network architecture and loss design (**Fig. 4b**, Methods, Extended Data Fig. 7 and Supplementary Figs. 6). We reasoned that the inclusion of an additional view essentially rationalizes the axial resolution enhancement, and the elaborately designed prior-guided learning scheme is able to effectively improve the dual-fusion procedure compared with either self-learning or conventional RL deconvolution (Supplementary Note 4). To validate the performance of RL-DFN, we examined it on synthetic image stacks of tubular structures and spherical shells (**Fig. 4c**, Extended Data Fig. 6, and Supplementary Note 5). We found that although current self-learning-based methods could generate perceptually good isotropic images under such ideal conditions where the simulated sample itself is isotropically distributed, RL-DFN more precisely reconstructed the fine details of both structures with isotropic resolution and higher PSNR (**Fig. 4d** and Extended Data Fig. 6).

**Fig. 4.**
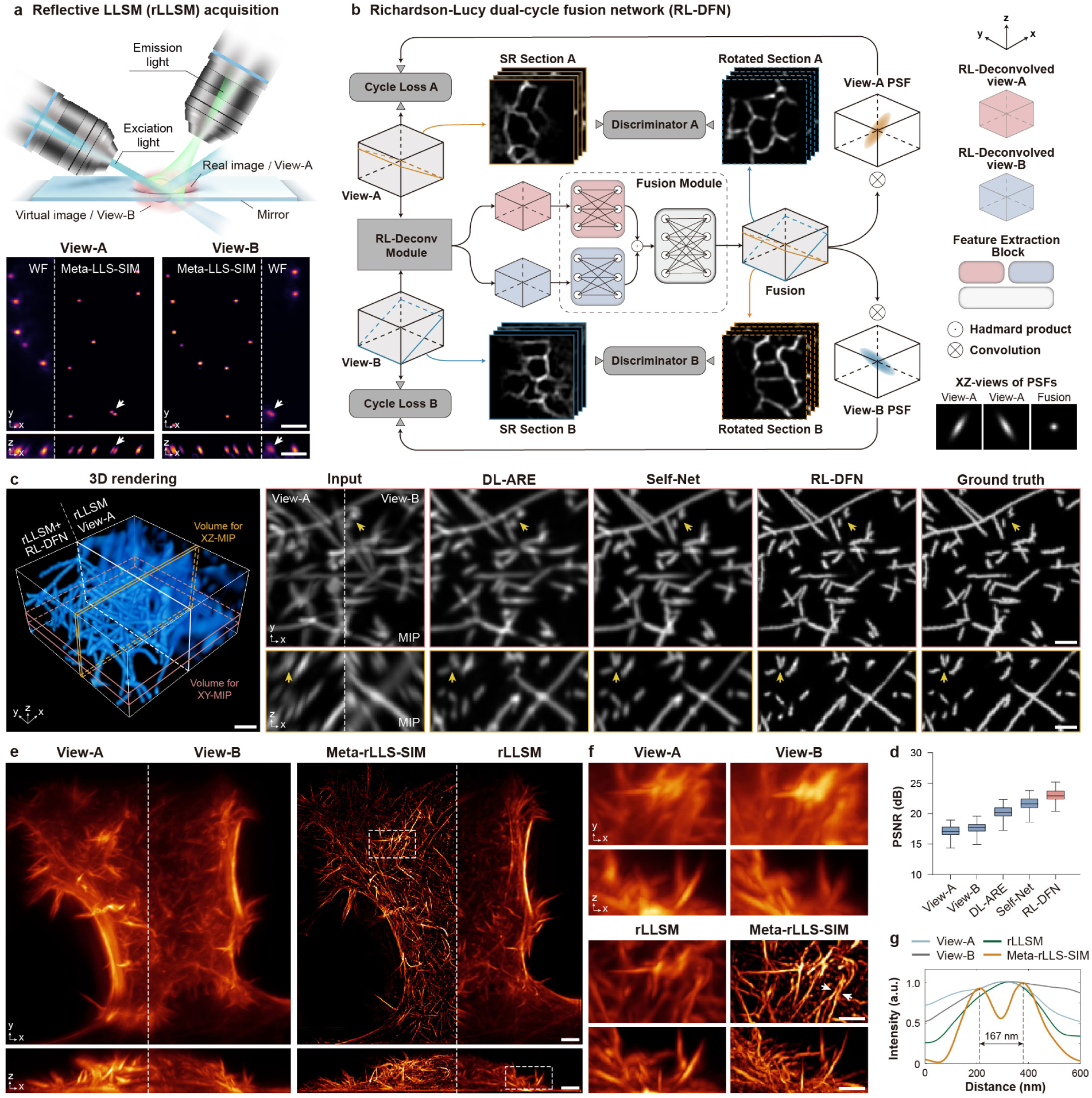
Near-isotropic SR reconstruction by Meta-rLLS-VSIM. **a**, Illustration of the acquisition process via reflective lattice light-sheet microscopy (rLLSM). The lower panel shows representative view-A and view-B images acquired by rLLSM system (labelled by WF) or processed with the laterally isotropic SR reconstruction algorithm, i.e., the Meta-VSI-SR method (labelled with Meta-LLS-VSIM). **b**, Schematic network architecture and data forward propagation of RL-DFN. The x-z views of PSFs of view-A, view-B, and network output are shown in the bottom right corner. **c**, 3D renderings (left) of synthetic wide-field microtubule data and its isotropic SR reconstruction by RL-DFN, and corresponding orthogonal views generated via deep-learning for axial resolution enhancement (DL-ARE)^19^, Self-Net^20^, and RL-DFN. Partial synthetic view-A and view-B, as well as the ground-truth images are provided for reference. **d**, Statistical PSNR comparisons of DL-ARE, Self-Net, and RL-DFN on the synthetic microtubule images. The PSNR values for original view-A and view-B are plotted for reference. **e**, Representative x-y and x-z MIPs of view-A, view-B, joint deconvolved images (labelled with rLLSM), and Meta-rLLS-VSIM reconstructions from view-A and view-B. **f**, Magnified images of the boxed regions in e. **g**, Plots of the intensity profiles of the view-A, view-B, joint deconvolved image, and Meta-rLLS-VSIM image along the lines indicated by the two arrowheads in f. Scale bar, 2 μm (a), 1 μm (3D rendering view of c), 1.5 μm (planar MIP view of c), 3 μm (e), 1.5 μm (f).

Subsequently, in order to equip our Meta-LLS-VSIM system with reflection-based axial resolution enhancement (Meta-rLLS-VSIM), we constructed an integrated 3D isotropic reconstruction framework that consists of epifluorescence contamination removal, image stack deskew, laterally isotropic reconstruction and rotation, dual-view separation and registration, and the final dual-view fusion with RL-DFN (Extended Data Fig. 8 and Supplementary Note 6). We demonstrate that after the entire processing workflow, the spatial resolution reaches 119 nm (n=12 beads) laterally and 157 nm axially (n=9 beads), improving volumetric resolution by 15.4-fold compared with conventional LLSM (Extended Data Fig. 9). Next, we compared Meta-rLLS-VSIM with other isotropic reconstruction techniques using experimentally acquired data of F-actin, microtubules, and ER, and found that due to the huge morphology difference between lateral and axial sections, self-learning-based methods cannot recover axial resolution satisfactorily and generate massive artifacts (Extended Data Fig. 6). In contrast, Meta-rLLS-VSIM dramatically improves resolution in all dimensions from the original dual-view acquisitions (**Fig. 4e**), achieving the highest spatial resolution and best SR quality compared with either deconvolution-based reflective LLSM (**Fig. 4f, g**) or DL-based computational reconstruction (Extended Data Fig. 6).

### Rapid, long-term, near-isotropic SR 4D subcellular imaging

In Meta-rLLS-VSIM, the reflective configuration introduces the other complementary view of the sample within single captures, not sacrificing any imaging metrics relative to Meta-LLS-VSIM or LLSM. To demonstrate the superior capability of Meta-rLLS-VSIM in fast, long-term, near-isotropic SR 4D imaging applications, we imaged Cos-7 cells transferred with GFP-SKL, ER-mCherry, and Lyso-Halo (labelling peroxisomes, ER, and lysosomes, respectively), collecting 400 three-color whole cell volumes at 12-second intervals (**Fig. 5a**, Supplementary Video 6). Despite working at almost the same speed and photon budget with standard LLSM, Meta-rLLS-VSIM dramatically improved the resolution both laterally and axially compared to the diffraction limit, clearly resolving reticular ER, punctate Perox, and ring-like Lyso (**Fig. 5b**). We quantified the spatial resolution of Meta-rLLS-VSIM to be 122 nm in lateral (**Fig. 5c**) and 162nm in axial (**Fig. 5d**), respectively, by measuring the full width at half maximum (FWHM) of intensity profiles of ER or Perox sections indicated by arrowheads in Fig. 5b (n=10). The power spectrum coverages shown in Fig. 5e also indicated that Meta-rLLS-VSIM expectedly reached a near-isotropic resolution in live-cell imaging experiments. Furthermore, we reduced the acquisition interval to 8 seconds between adjacent timepoints and increased the total number of three-color volumes to over 720, realizing 1.6-hour long visualization of another Cos-7 cell without any noticeable photobleaching and phototoxicity (Supplementary Video 7, Extended Data Fig. 10), which corresponds to prolonging the imaging duration by more than 10-fold compared to the latest isotropic 3D-SIM working at even the single-color mode^19^. The ultrahigh volumetric resolution and extended time course of Meta-rLLS-VSIM allowed us to investigate interactions between multiple organelles and discover interesting cases, e.g., tubular ER generation by hitchhiking on a moving Lyso (**Fig. 5f**), much more easily and explicitly than ever before (**Fig. 5g**).

**Fig. 5.**
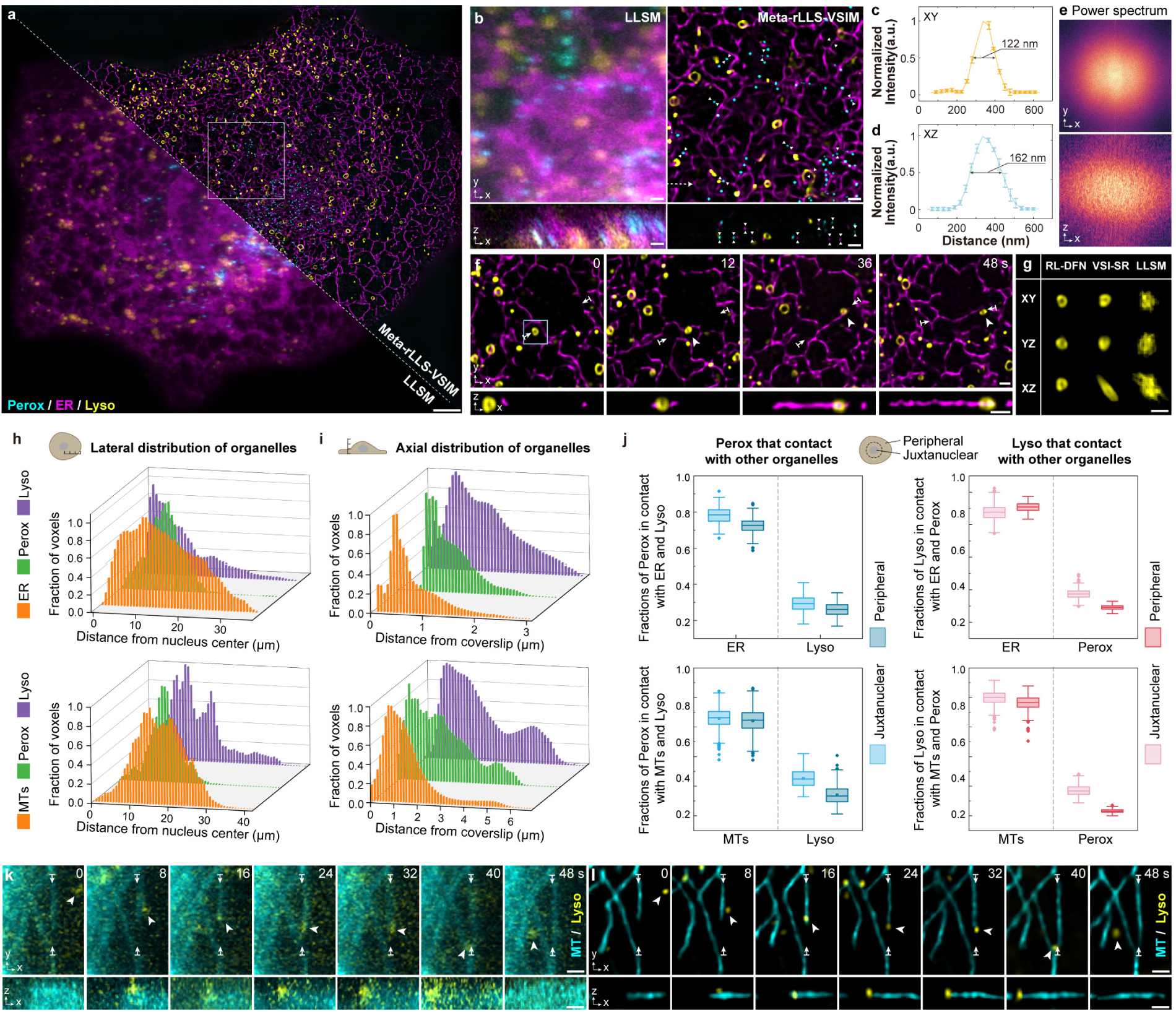
Rapid, long-term, near-isotropic SR 4D subcellular imaging by Meta-rLLS-VSIM. **a**, Representative LLSM (bottom left corner) and Meta-rLLS-VSIM (top right corner) image of a live Cos-7 cell expressing GFP-SKL, ER-mCherry, and Lyso-Halo from a three-color video of 400 timepoints (Supplementary Video 6). **b**, Magnified x-y and x-z sections of the region labelled with the white box in a imaged by LLSM (left) and Meta-rLLS-VSIM (right). **c**,**d**, Lateral (c) and axial resolution (d) evaluation by profiling intensity along the lines indicated by white arrowheads in b (n=10 for both lateral and axial resolution). **e**, Logarithmic power spectrum coverages of the Perox images displayed in b. **f**, Time-lapses Meta-rLLS-VSIM images of ER and Lyso showing a special case of ER generation hitchhiking on a moving Lyso. **g**, Comparison of images of a Lyso indicated by the blue box in f, imaged by LLSM and reconstructed by RL-DFN and VSI-SR. **h**,**i**, Distributions of organelles in the lateral (h) and axial (i) dimensions of two Cos-7 cells. **j**, Box whisker plots showing the fraction of Perox (left column) and Lyso (right column) contacting each of the other labelled compartments in the juxtanuclear or peripheral regions of the cell (n=721 and 812 frames for the cells labelled ER and MTs, respectively). **k**,**l**, Time-lapse LLSM (k) and Meta-rLLS-VSIM images showing a microtubule elongated by hitchhiking on a moving Lyso. Gamma value, 0.7 for Meta-rLLS-VSIM images of ER, MT, and Perox in a, b, f, k, and l. Scale bar, 3 μm (a), 1 μm (b, f, g, k, and l).

Different organelles each exhibit a characteristic distribution and dispersion pattern within the three-dimensional space that is affected by cytoskeleton such as microtubules^35^. To quantify the spatiotemporal coordination and interactions between multiple organelles and cytoskeleton, we employed Meta-rLLS-VSIM to image another Cos-7 cell labelled by 3×mEmerald-Ensconsin, mCherry-SKL, and Lamp1-Halo for 812 timepoints at 8 seconds per three-color whole cell volume (Supplementary Video 8), and characterized the organelle distribution patterns in both lateral and axial dimensions (**Fig. 5h, i**, Supplementary Fig. 7, Methods). We observed that in the lateral dimension, ER and MT both had a wide distribution, whereas Perox displayed a narrow distribution around the juxtanuclear zone (**Fig. 5h**). In the axial dimension, ER, MTs and Perox were generally localized throughout the cell and kept relatively stable within the hour-long observation window, while the distribution of Lyso varied along with time markedly (**Fig. 5i**, Supplementary Fig. 7), implying their high dynamics and complex functionalities in maintaining cell homeostasis^36^. Next, we quantified the fraction of globular organelles (Perox and Lyso) that made contacts with ER and MTs by tracking individual Lyso and Perox and mapping their contacts with other organelles over time (Methods, **Fig. 5j**). We observed that ER and MTs showed high contact rates with other two globular organelles in both peripheral and juxtanuclear regions, whereas the Perox-Lyso contacts preponderantly happened in the juxtanuclear region with relatively low contact rates (**Fig. 5j**). Additionally, during the investigation of the intracellular contacts, we noticed an interesting hitchhiking event between MTs and Lyso (**Fig. 5k, l**), which was previously reported to happen on ER tubules that adhered to moving mitochondria or Lyso in mammalian cells^37^. These observations and results illustrate the superior SR capability of Meta-rLLS-VSIM in developing hypotheses about cellular organization and dynamics.

## Discussion

In this work, we first presented a virtual structured illumination SR reconstruction strategy, that extends the 1D SR capability of conventional LLS-SIM to all lateral dimensions without any hardware modification. By collaboratively optimizing the DNN architecture, loss function, and joint deconvolution algorithms, we demonstrate the VSI-SR scheme achieves superior SR reconstruction for LLSM images with laterally isotropic resolution of 180 nm (Extended Data Figs. 1-3, 9). Moreover, to overcome the longstanding problem of high costs for image SR network training in practical usage, we devised Meta-LLS-VSIM by synergizing the advantages of both optical front-end and meta-learning back-end methodologies, that could rapidly adapt to any type of biological specimens or data SNR within tens of seconds, reducing the training data and time cost by 12-fold and 720-fold, respectively. Of note, although demonstrated on SR processing for LLSM, the proposed meta-learning scheme can be well applied to other DL-based image restoration tasks for various imaging modalities, such as denoising^12,15,38^, phase retrieval^39^, virtual staining^40,41^, etc.

Furthermore, to enhance the axial resolution while not sacrificing other imaging metrics, we incorporated reflective acquisition strategy^25^ into our Meta-LLS-VSIM system to simultaneously capture two complementary views of the specimen with one-time scanning. Then we devised a Richardson-Lucy dual-view fusion network, dubbed RL-DFN, that combined an unmatched back projector-based RL iteration^34^ for fast deblurring and the conditional generative adversarial network^42^ (cGAN) for feature fusion in a self-supervised manner. Compared with existing methods that directly improve axial resolution by self-learning, RL-DFN is more physically rational and achieves the best performance in axial resolution enhancement for both simulated and experimental data. We also constituted an integrated reconstruction framework, which covers the whole processing pipeline from the original dual-view acquisitions to near-isotropic SR reconstructions, improving the spatial resolution to 120 nm in lateral and 160 nm in axial.

Taken all these advances together, we demonstrate that Meta-LLS-VSIM and Meta-rLLS-VSIM fulfill the unmet requirement for 4D SR imaging at ultrahigh spatiotemporal resolution with long duration of hundreds of multi-color volumes for a large variety of subcellular bioprocesses, revealing fast dynamics and long-term interactions of multiple organelles. In particular, by integrating the proposed SR reconstruction scheme with synchronized rolling-shutter confocal slit detection, the isotropic SR capability can be further extended to image thick or scattering samples such as pollen tubes, *C. elegans* and mouse embryo. These characteristics underlie the versatile utility and superior performance of Meta-LLS-VSIM and Meta-rLLS-VSIM methodology, offering great opportunities to better investigate and understand diverse biological phenomena.

## Acknowledgements

The authors thank Dr. Hao Wang for the donor pollen tubes and help in recording the growth process of them. This work was supported by grants from National Natural Science Foundation of China (32125024, 92254306, 32271513, 62088102, and 62231018); the Ministry of Science and Technology (2021YFA1300303 and 2020AA0105500); the Chinese Academy of Sciences (ZDBS-LY-SM004 and XDA16021401); Collaborative Research Fund of the Chinese Institute for Brain Research, Beijing (2021-NKX-XM-03); China Postdoctoral Science Foundation (2022M721842, 2023T160365); the New Cornerstone Science Foundation; the Shuimu Tsinghua Scholar Program (2022SM035).

## Author contributions

D.L., C.Q. and Z.L. conceived the idea. D.L. and Q.D. supervised the research. D.L. and C.Q. designed the experiments. Yong Liu, S.Z. and X.D. built and improved the microscope systems at the Institute of Biophysics under the supervision of D.L. C.L., J.G., W.F., and T.J. prepared samples and performed experiments. C.Q., Z.L., Z.W., Yuhuan Lin, and Q.W. analyzed the data with conceptual advice from D.L. and C.Q. C.Q., Z.L., Z.W., Yuhuan Lin, and Y.F. composed the figures and videos under the supervision of D.L. C.Q., Z.L., and D.L. wrote the manuscript with input from all authors. All authors discussed the results and commented on the manuscript.

## Competing interests

D.L., C.Q., S.Z., and Z.L. have a pending patent on the presented frameworks. Remaining authors declare no competing interests.

## Methods

### LLS-SIM system

The home-built LLSM/LLS-SIM system was developed from the original design^8^. Three lasers of 488 nm, 560 nm, and 640 nm (MPB Communications) were controlled by an AOTF, and then illuminated the LLS pattern displayed on the SLM. In the SI mode, the LLS patterns of 3-phase were sequentially displayed onto the SLM and synchronized with the programmed “ON” time of AOTF. The diffracted light was filtered by an annular mask equivalent to 0.5 outer NA and 0.375 inner NA for the excitation objective (Special Optics). Subsequently, the filtered excitation light was scanned by the *z*-galvo-objective in a step size of 0.2 μm or by sample piezo in a step size of 0.39 μm to acquire the volumetric raw LLS-SIM images. In the ordinary sheet-scan mode, a fixed LLS pattern was filtered by another annular mask equivalent to 0.35 outer NA and 0.14 inner NA to elongate the light-sheet, then quickly dithered by *x*-galvo (Cambridge Technology, 6210H), and then scanned by using of sample piezo in a step size of 0.39 μm, rather than using *z*-galvo-objective scan mode, because the sample holder has lighter-weight than objective, which allows the sample piezo to run in high-speed with small hysteresis. Moreover, instead of driving the sample piezo with the ramp wave, we used the triangle wave to minimize the flyback time when reversing the scanning direction of the piezo stage. For the rolling-shutter confocal slit-scan mode, a Gaussian beam instead of LLS patterns was quickly scanned along the *x*-axis to create the light-sheet, which is synchronized with the camera’s rolling shutter to form a virtual confocal slit effect. We used an active 15-pixel column in the sCMOS camera (Hamamatsu, Orca Fusion) in our experiments for optimal tradeoff between the SNR and contrast.

Live cell specimens were placed in a custom designed microscope incubator (OKO lab, H301-LLSM-SS316) to maintain the physiology condition of 37℃ and 5% CO2 during imaging. Fluorescence emission was collected by the detection objective (Nikon, CFI Apo LWD 25XW, 1.1NA) and captured by the sCMOS camera. The reflective coverslips used in the Meta-rLLS-VSIM experiments were customized by sputtering a 150-nm-thick aluminum film over the round glass coverslip (*φ*=12 mm × 0.17-mm-thickness) and then protected with a 700-nm-thick layer of SiO_2_. The imaging conditions of live-cell experiments are detailed in Supplementary Table 1.

### Network architecture of the VSI-SR model

The VSI-SR neural network model is constructed based on the conditional generative adversarial network (cGAN)^43^, consisting of two models: one is the generator *G*, i.e., the model used for inference (in the main manuscript the ‘VSI-SR model’ referred to *G* except as otherwise noted), which transfers low-resolution (LR) images into SR images, and the other is the discriminator *D* which determines whether an image comes from training dataset or *G*. Specifically, the VSI-SR model *G* receives LR fluorescence images with the size of *H* x *W* x *C*_in_, and outputs SR images upscaled by 1.5-fold with the size of 1.5*H*x 1.5*W* x *C*_out_. We adopted 7 and 3 as *C*_in_ and *C*_out_, respectively, in our experiments, which empirically achieve the best SR performance (Extended Data Fig. 1, and Supplementary Note 1). The generator *G* begins with a 2D convolutional layer for channel augmentation, and then the output feature maps are fed into four sequential residual groups (RGs), each of which contains four channel attention blocks as is defined in previous paper^44^. After RGs, the extracted features are upscaled, and fed into two Conv-LeakyReLU modules to produce the final SR images. The discriminator *D* takes the outputs from *G* or GT SR images as the input, and provides the probability of the input being the GT. *D* begins with 11 Conv-LeakyReLU modules for deep feature extraction. Then their outputs are fed sequentially into a global average pooling layer, two fully connected layers with a LeakyReLU activation, and a sigmoid activation function to output the estimated probability. The overall network architecture of *G* and *D* are depicted in Supplementary Fig. 1.

During the training phase, we elaborately designed a combined objective function for *G* (Extended Data Fig. 3), which consists of four terms: mean square error (MSE) loss, structural similarity (SSIM) loss, fast Fourier transform (FFT) loss, and the discriminative loss:

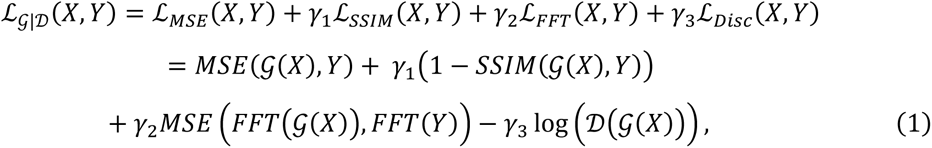

where *X* and *Y* denote the LR input images and the SR targets, respectively; *γ*_1_, *γ*_2_, and *γ*_3_ are weighting scalars to balance the contributions from these four terms, and we set *γ*_1_ = 0.1, *γ*_2_ = 1, *γ*_3_ = 0.1 empirically in our experiments. Within the combined loss function, the first two terms, i.e., ℒ_MSE_ and ℒ_SSIM_, penalize the difference between predictions and GT images in the spatial domain, while the third term ℒ_FFT_ minimizes the errors in frequential space, which we found to be helpful to learn more high frequency information^45^ (Extended Data Fig. 3). The last term ℒ_Disc_ serves as regularization based on the knowledge from the discriminator *D*, which also benefits for generating finer details of biological structures.

On the other hand, the objective function of *D* is a binary cross-entropy function described as:

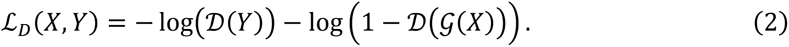

*G* and *D* are trained alternately in the inner loop of meta-learning (see details in the next section), during which they compete with each other, and finally reach an equilibrium state.

### Meta-model training and finetuning

The datasets of different biological specimens were acquired via our home-built LLS-SIM system. We generally acquired raw LLS-SIM images from about 30-50 distinct ROIs for each type of samples. For each ROI, five different levels of light intensity ranging from low (about 100 average photon counts) to high (more than 1000 average photon counts) fluorescence level were acquired, and the images of the highest fluorescence level (i.e. the GT raw images) were reconstructed into high-quality GT LLS-SIM images via the conventional LLS-SIM reconstruction algorithm. In order to generate the training dataset for meta-learning, we established 20 diverse task datasets from 10 distinct biological specimens with low (about 200 average photon counts) and high (more than 1000 average photon counts) fluorescence levels. Each task tackles a specific image SNR, i.e., low or high SNR, of a specified biological structure. It needs to be emphasized that instead of generating more task datasets by introducing various fluorescence levels, we only considered low or high SNR tasks since we found that too many tasks that were similar to each other would degrade the meta-model into an ordinary one without fast adaptation capability. The ten biological specimens used in meta-training in this paper were the outermost granular component, chromosomes, innermost fibrillar center, fibrillarin, Lyso, MTs, F-actin in pollen tubes, inner mitochondrial membrane, ER in adherent Cos-7 cells and ER in mitotic Hela cells during metaphase. The detailed information of the datasets used for training the meta-model is shown in Supplementary Table 2.

The pre-processing procedure of the raw data contains following steps: (i) applying deskew to all LLSM images (averaged from the raw LLS-SIM images) and their corresponding GT LLS-SIM images; (ii) removing the camera background, i.e., ∼100 sCMOS counts, for LLSM images and applying a 2D Gaussian filter with the standard deviation of 0.6 pixels for GT LLS-SIM images to slightly suppress the noise-induced reconstruction artifacts; (iii) normalizing all images to [0,1]. Then, the whole dataset was augmented into ∼ 54, 000 image patch pairs of LLSM patches (64 x 64 x 7 voxels) and their corresponding GT LLS-SIM patches (96 x 96 x 3 voxels), that is, 2,700 pairs for each task. As illustrated above, we adopted a multi-slice 2D training scheme which considered volumetric LLSM data as multi-channel 2D images, and inferred the central three x-y slices from the corresponding seven consecutive input x-y slices. This scheme outperformed others since it fully took advantage of the spatial continuity of z-axis of both LLSM inputs and LLS-SIM targets (Extended Data Figs. 1 and 2).

The training process of meta-learning is different from conventional gradient descent since in meta-training, the ultimate goal is to train a model such that fast learning and great improvement occur on a new task within a small number of gradient updates using minimal training data. Inspired by the MAML algorithm^23^, we devised a meta-learning strategy for our cGAN-based VSI-SR model, which generally consisted of outer-loop and inner-loop. In the outer-loop, three tasks were randomly sampled from task sets and fed into the inner-loop to calculate task-specified loss at every iteration. Then, all task-specified losses were used to update the meta-generator *G* and meta-discriminator *D* via the Adam optimizer. The outer learning rates of *G* and *D* were set as 1 x 10^-4^ and 2 x 10^-5^, respectively, in our experiments. In the inner loop, 16 LLSM and LLS-SIM image pairs were randomly sampled from every selected task, in which 8 pairs were used as the supported set and 8 pairs as the query set. The basic *D* learner and *G* learner were alternately updated three times with the supported set via the SGD optimizer, then every task-specified loss can be evaluated with the updated learners and the query set. The inner learning rates of *G* and *D* were 1 x 10^-2^ and 2 x 10^-3^, respectively. In our experiments, deep-learning models were trained and implemented on a computer work station equipped with an Intel(R) Xeon(R) Platinum 8358 CPU at 2.60 GHz and three NVIDIA A800 graphic processing cards with Python version 3.7 and Pytorch version 2.1.0. The meta-training process typically lasted for ∼24 hours with ∼100,000 mini-batch iterations when three A800 cards were used simultaneously for distributed data parallel training (DDP) of Pytorch. It is noteworthy that the meta-training is a one-time procedure, and all VSI-SR models used in live imaging experiments of this work were finetuned from the same Meta-VSR-SR model. The overall meta-training procedure is depicted in Extended Data Fig. 5.

At the finetuning stage, three ROIs of a target biological specimen were augmented into 600-1000 LLSM/LLS-SIM patch pairs as the supported set, and we finetuned the meta-model *G* with supported set in 30 gradient steps with a batch size of 20 via the SGD optimizer, of which the learning rate was 2-10 times higher than the inner learning rate of *G*. After finetuning, the meta-model *G* quickly adapted to the target specimen, yielding a specified VSI-SR model for the target biological structure. The finetuning process was typically less than 10 seconds with an NVIDIA A800 or RTX3090 graphic processing card on our computer work station.

### Laterally isotropic SR reconstruction via VSI-SR

Given the fluorescence distribution *S* of a sample, the image captured with LLSM can be approximated as *I*_LR_ ⦻ *S*⊗ *P*_LR_, where *P*_LR_ represents the system PSF. If illuminating the sample with sinusoidal stripe patterns in orientation *φ*_0_ (assume *φ*_0_⦻0 for simplicity afterward), we get the anisotropically frequency extended SR image by the conventional SIM reconstruction algorithm expressed as *I*_SR-0_ ⦻ *S*⊗ *P*_SR-0_. Here, *P*_SR-0_ is the anisotropic SR PSF as a narrowing version of *P*_LR_ in orientation *φ*_0_ ⦻ 0. As described above, the VSI-SR or Meta-VSI-SR model was trained to map from the LLSM image *I*_LR_ to anisotropic SR image *I*_SR-0_ and we applied the trained VSI-SR model to process the image rotated by *φ*-angle from the original *I*_LR_ (denoted as *I*_LR-_*φ* ⦻ *RφI*_LR_ ⦻ *RφS* ⊗ *P*_LR_, where *R_φ_* is the rotation operator by angle *φ*) and got the anisotropically super-resolved prediction *f*_*θ*_{*I*_LR-_*φ*} ⦻ *f*_*θ*_{*RφS* ⊗ *P*_LR_}, where *f*_*θ*_ denotes the forward propagation of the VSI-SR model with trained parameters *θ*. Then by rotating the output SR image backward, we got the final estimation of the anisotropic SR image in orientation *φ*:

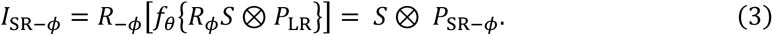

Here, rotating the anisotropic PSF *P*_SR-0_ in the illumination orientation by angle *φ* approximates the anisotropically narrowed PSF *P*_SR-*φ*_ in orientation *φ*. Generally, three orientations of illumination equally spaced between 0 to π are used to isotropically fill the OTF extension. Thus, we simply set *φ* to be 0, π/3, and 2π/3.

To compute the final isotropic VSI-SR image, we combined the above anisotropic predictions in Fourier space through a generalized Wiener filter approach. Each anisotropic SR image *I*_SR-*φ*_ transformed to the Fourier space 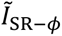 is a summation of the zero-order frequency component, denoted as 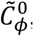, and the symmetrically distributed first-order components 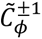. We simplify the zero component to be 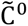 since it is unaffected by the illumination. The image assembled from *m* orientations in Fourier space is expressed as

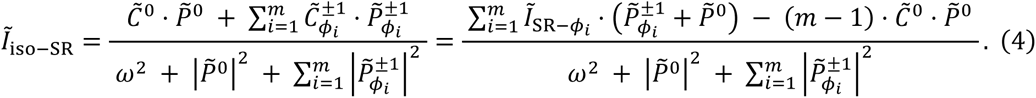

The wiener filtering compensates for the frequency attenuation introduced by the system OTF, with the parameter *ω* relaxing the compensation in regions where the OTF value is low and set according to the imaging SNR. The 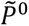 and 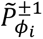 represent the OTF for zero-order and first-order frequency components, respectively, and the zero component 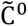 is computed as the Fourier transform of the LLSM image *I*_LR_.

### Network architecture of RL-DFN

The RL-DFN is constructed based on the cGAN framework with dual discriminative cycles. Specifically, it consists of three individual models: a 3D generator *G*_RL-DFN_ that fuses features from dual-view inputs and reconstructs isotropic SR image volumes, and two 2D discriminators *D*_A_ and *D*_B_ that distinguish whether a sectioned image comes from the input data or from the SR volume generated by *G*. Of note, here ‘3D’ and ‘2D’ refer to the feature dimensions propagated in the neural networks, i.e., the 3D model utilizes 3D convolutional layers and the 2D model uses 2D ones.

The network architecture of *G*_RL-DFN_ is illustrated in Fig. 4b and detailed in Supplementary Fig. 6a, which begins with a Richardson-Lucy deconvolution module (RDM) that explicitly executes one classical RL update step for two anisotropic inputs *I*_A_(*x*) and *I*_B_(*x*). The calculation in RDM can be formulated as

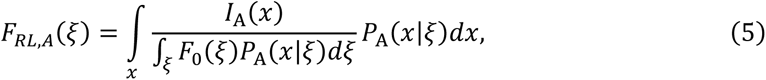

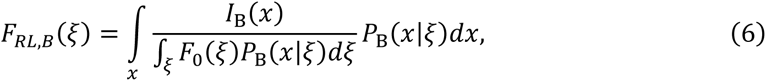

where *F*_0_(*ζ*) denotes the average of *I*_A_(*x*) and *I*_B_(*x*), *P*_A_(*x*|*ζ*) and *P*_B_(*x*|*ζ*) are PSFs of view A and B, respectively, both as functions of their respective pixel locations *x* and *ξ*. Then the RL-deconvolved volumes *F*_RL,A_ and *F*_RL,B_ are fed into the fusion module to implicitly generate an isotropic SR volume. In the fusion module, two identical feature extraction blocks, FEB_A_ and FEB_B_, are separately applied to extract features from *F*_RL,A_ and *F*_RL,B_, with each FEB consisting of four convolutional layers (two of them followed by batch normalization and LeakyReLU activation) for feature extraction and a sigmoid activation to normalize the output to [0, 1]. Afterwards, the output features are merged by Hadamard product to obtain the fusion feature in embedding space. Finally, another U-shaped feature extraction block FEB_U_(detailed in Supplementary Fig. 6a) is applied to generate the final isotropic SR volume. The overall operation in the fusion module can be represented as

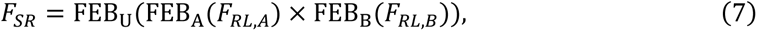

where x denotes the Hadamard product.

The discriminator *D*_A_ and *D*_B_ share the same architecture and objective function, hence they are collectively denoted as *D*_AB_ hereafter. *D*_AB_ consists of six convolutional layers, in which the central four convolutional layers are followed by batch normalization and LeakyReLU activation with a leaky factor of α=0.1. Then the output of the last convolutional layer is fed into the sigmoid activation function to obtain the predicted probability.

The objective function of *G*_RL-DFN_ and *D*_AB_, denoted as 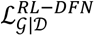 and 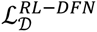, respectively, are defined separately in a self-supervised manner. Specifically, 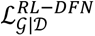 is defined as the sum of three terms: cycle loss *L_cycle_*, discriminative loss *L_Disc_*, and total variational (TV) regularization *R*_TV_, which can be formulated as

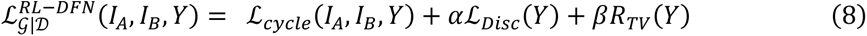

where *I_A_*, *I_B_* are the original dual-view inputs, *Y* is the output SR volume by *G_RL-DFN_*, and *α*, *β* are scalar weighting factors to balance the corresponding terms, which are set empirically to *α* = 0.02, γ=2 x 10^-6^ for best performance in our experiments.

In the objective function described in Eq. (8), the cycle loss ℒ_cycle_ penalizes the difference between the anisotropic input of each view and the isotropic output of *G*_RL-DFN_ degraded by corresponding PSF *P*_A_ and *P*_B_. As such, the ℒ_cycle_ contains two components as such

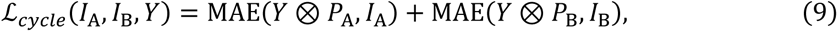

where MAE denotes the mean absolute error. ℒ_cycle_ is devised based on the physical model and utilizes the prior knowledge of PSFs, which accelerates the training process and guarantees the output fidelity. On the other hand, the discriminative loss ℒ_Disc_ is associated with the predicted probability by *D*_A_ and *D*_B_ for image sections sampled from the *G*_RL-DFN_ output. The sectioning operation is performed using a rotation sectioning module, which samples two 2D slices *S*_A_ and *S*_B_ along the orientations perpendicular to the optical axis of view-A and view-B with a random deviation ranging in [-π/6, π/6] (Extended Data Fig. 7a). We empirically found that the introduction of the randomness of the sampling angle improved the robustness and overall performance of RL-DFN. The ℒ_Disc_ can be formulated as

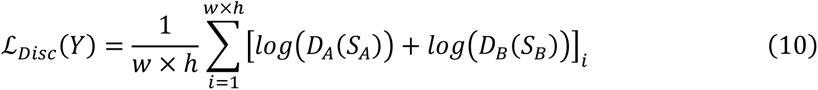

The TV regularization *R*_TV_ is applied as a spatial continuity prior of biological specimens, which is calculated by

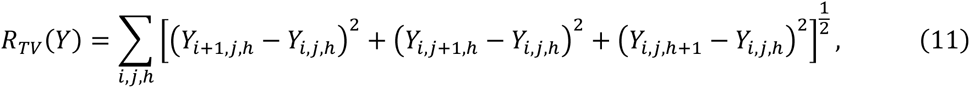

where *i*, *j*, *h* denote the 3D coordinates of *Y*. The objective function of the discriminator *D*_AB_ is defined as the binary cross-entropy. Taking *y* as the output of *D*_AB_ and *y*_GT_ as the ground truth matrix (all 0 or 1 matrix in practice), then 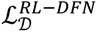 can be described as

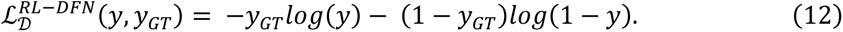

During the training process, *G*_RL-DFN_ and *D*_AB_ is playing an adversarial game. The objective of *G*_RL-DFN_ is to deceive *D*_AB_ into taking the sections sampled from its output as the real isotropic SR section sampled along the SR orientation of the anisotropic inputs. In contrary, the objective of *D*_AB_ is to determine if the input image is sectioned from *G*_RL-DFN_’s output or the input anisotropic volumes.

### Training and application of RL-DFN

To generate the training dataset of RL-DFN, we applied random cropping from the feature-only regions, and augmented the original data into thousands of volume pairs of view-A and view-B (256×256×64 pixels for each). During the training process, we randomly initialized the networks and trained the generative and discriminative models with the Adam optimizer and a typical starting learning rate of 1 x 10^-4^. The generator and discriminator were updated once and twice in each iteration with a batch size of 3 and 2, respectively. The rotation sectioning module sampled a new set of 2D slices (250×60 pixels for each) before each training iteration along randomly sampled orientations as described above. The overall training process of RL-DFN typically lasted 15,000 minibatch iterations, taking ∼8 hours. After the self-supervised training, a well-trained RL-DFN, more specifically the generator of RL-DFN, can be utilized in the dual-view fusion task for the same type of data within or out of the training dataset. It typically took ∼1 second to merge two anisotropic image volumes of 512×512×150 pixels.

### Organelle distribution and contact quantification

For analysis of organelle spatial distribution shown in Fig. 5h and i, the pixel-based quantification is performed. Each channel of the images was low-pass filtered using a 3D Gaussian kernel with sigma of 1 pixel. The smoothed images were then binarized with a threshold optimized by the Otsu algorithm^46^. The histograms of positions of pixels above the threshold in centroid coordinates are calculated. The nucleus center in lateral plane was identified visually and the cell boundary was identified by using ER as the marker structure. The pixel-based quantification was applied to each timestamp of the time-lapse images, and an averaged result on both the full recording time and for each 10-minute duration was analyzed (Supplementary Fig. 7).

For analysis of the organelle interactions shown in Fig. 5j, the multi-channel images were first segmented using Imaris software with the *surfaces* tool. Background subtraction and manual thresholding were performed for each channel, and split touching objects were used for the globular organelles (i.e., peroxisomes and lysosomes). Next, distance transformation was performed on each of the segmented surface channels to calculate the distance between organelles. Segmented objects with a minimum distance of less than one voxel of the isotropic SR volume between surfaces were considered to be interacting. For analysis of organelle interactions related to spatial positions, the circular region in the cell with a thickness greater than a predefined threshold (∼2.5 μm) was manually masked to be juxtanuclear. The contact fraction for each granular organelle was calculated by dividing the count of objects in contact with other organelles by its total count.

### Cell culture, lentivirus packaging, stable cell line and transfection

HeLa, Cos7 and HEK293T cells were cultured in DMEM (Gibco, cat. no. 11965092) supplemented with 10% FBS (Gibco, cat. no. 10099141C) and 1% penicillin–streptomycin (Thermo Fisher, 15140122) under 37 ℃ and 5% CO2 until 60-80% confluency was reached. For live cell imaging experiments, the coverslips were pre-coated with 50 mg/ml of collagen for 1 hour, and cells were then seeded onto the coverslips for 16 hours before transient transfection or imaging.

For lentivirus packaging, 1 μg lentiviral transfer vector DNA, together with 0.5 μg psPAX2 packaging vector and 0.5 μg pMD2.G envelope vector DNA were co-transfected to 90% confluence HEK293T cells in a 6 cm petri dish using Lipofectamine 3000 (Invitrogen) following the manufacturer’s protocol. After 2 days, supernatant containing viral particles was harvested and filtered with a 0.22-μm filter (Millipore). Supernatant was immediately used for transduction or stored at −80 °C in aliquots.

For construction of stable cell line, HeLa cells were cultured to 10–20% confluency in six-well plates, and 100–300 μl of filtered viral supernatant was added to the cells. Specifically, lentiviruses carrying corresponding fluorescent protein-tagged organelle markers, including mEmerald-NPM1, mCherry-H2B, Halo-RPA49, FBL-mEmerald, Mito-dsRed, mEmerald-Lifeact, Lyso-Halo, and KDEL-mCherry, were used. Media containing virus were replaced with fresh growth medium 24 hours after infection. 48 hours later, the cells were enriched by flow cytometer (FACSAria III, BD Biosciences) and then plated one cell per well into 96-well plates. Monoclonal cells were used for our experiments.

For live cell imaging of Cos7 cells, samples were transfected with corresponding plasmids using Lipofectamine 3000 (Invitrogen, cat. no. L3000150) according to the manufacturer’s protocol. The plasmids used in transient transfection KDEL-mCherry, Lyso-Halo, GFP-SKL, mCherry-SKL and 3×mEmerald-Ensconsin. Where indicated, the cells transfected with Halo Tag plasmids were labelled with 10 nM JF642 ligand for 30 min before imaging. The inner mitochondrial membrane was marked and stained without transfection by using PK Mito Red (Genvivo, cat. No. PKMR-1).

### *Nicotiana tabacum* growth, transgenic plant generation and pollen *in vitro* germination

Tobacco (*Nicotiana tabacum*) plants were grown in a greenhouse at 22 °C under a light cycle of 12-h light and 12-h darkness and the transgenic tobacco plants expressing *UBQ10::Lifeact-GFP* were prepared as described before^47,48^. Fresh and mature pollen grains were collected from these individual plants and used for particle bombardment and *in vitro* pollen tube germination. Tobacco pollen grains were suspended in tobacco-specific pollen germination medium containing 10% sucrose, 0.01% boric acid, 1 mM CaCl_2_, 1 mM Ca(NO_3_)_2._4H_2_O, 1 mM MgSO_4._7H_2_O, pH 6.5 at 27.5°C in an 85 rpm orbital rotary shaker for 2 h. Germinated pollen tubes were gently and carefully pipetted onto the bottom cover slide for imaging.

### Mouse embryo preparation

Mice used in this study were C57BL/6J background. All animal experiments were approved by the Animal Care and Use Committees (IACUC) of the Institute of Biophysics, Chinese Academy of Sciences, Beijing, China. Pre-implantation embryos were isolated from 5-6-week-old females, superovulated by intraperitoneal injection of 5 international units (IU) of pregnant mares’ serum gonadotropin (PMSG; LEE BIOSOLUTIONS) and 5 IU human chorionic gonadotropin (hCG; Millipore) 48 h later, and mated with male mice. Zygotes were recovered at E0.5 in M2 medium (Millipore).

T7 promoter-kozak sequence-Tomm20-mEmerald was constructed into RN3P plasmid vector. Target sequence Tomm20-mEmerald was in vitro-transcripted with T7 in vitro transcription kit (Thermo Fisher, AM1344). Zygotes were microinjected with 330 ng/μL Tomm20-mEmerald with microinjection device (Eppendorf) and then cultured in KSOM medium (Millipore) in CO_2_ incubator (Thermo Fisher) at 37℃ with 5% CO_2_. When imaging using LLSM, the chamber filled with KSOM was pre-warmed with the live cell device of LLSM. Embryos were seeded on 12 mm coverslips suited for LLSM into the KSOM of chamber.

Experiments involving mouse tissue were performed in accordance with protocols approved by the Institutional Animal Care and Use Committee of CAS, Center for Excellence in Molecular Cell Science, Institute of Biochemistry and Cell Biology, Chinese Academy of Sciences.

### C. elegans embryo preparation

C. elegans strains were cultured at 20 °C on nematode growth medium (NGM) plates seeded with OP50 following standard protocols^49^. TV52712*[wyEx51119[dlg-1p::GFP::PLCdPH]*; *jcIs1[ajm-1::GFP+UNC-29(+)+rol-6(su1006)]*; *qxIs257 [ced-1p::nuc-1::mCherry + unc-76(+)]]* was used in this study. The plasmid *dlg-1p::GFP::PLCdPH* was constructed following the Clontech In-Fusion PCR Cloning System^50^ and microinjected to *jcIs1;qxIs257*. Extrachromosomal array *wyEx51119* marked epidermal cell membrane. *jcIs1* marked the apical junctional domain of C. elegans^50^*. qxIs257* marked lysosomes in epidermal cells^51^.

About 50 L4 stage transgenic worms were put onto NGM plates with freshly OP50 48 to 60 hours before experiments. Transgenic eggs were collected under the dissecting fluorescent microscope (Olympus MVX10), and mounted on 3% agarose pads. Lima bean to 2-fold stage embryos were then imaged using our LLSM system.

### Data availability

The LLS-SIM data for training the SR meta-model will be uploaded onto the publicly available Zenodo repository after the paper is accepted by a peer-review journal. Other data that support the findings of this study included in Figs. 1-5, Extended Data Figs. 1-10, Supplementary Figs. 1-7, Supplementary Notes 1-6, Supplementary Tables 1, 2, and Supplementary Videos 1-8 and all the source data presented in this paper are available upon reasonable requests.

### Code availability

The python codes of meta-learning, isotropic SR reconstruction, RL-DFN, several representative pre-trained models, as well as some example images for testing will be uploaded on GitHub after the paper is accepted by a peer-review journal.

## Extended Data Figures

**Extended Data Fig. 1.**
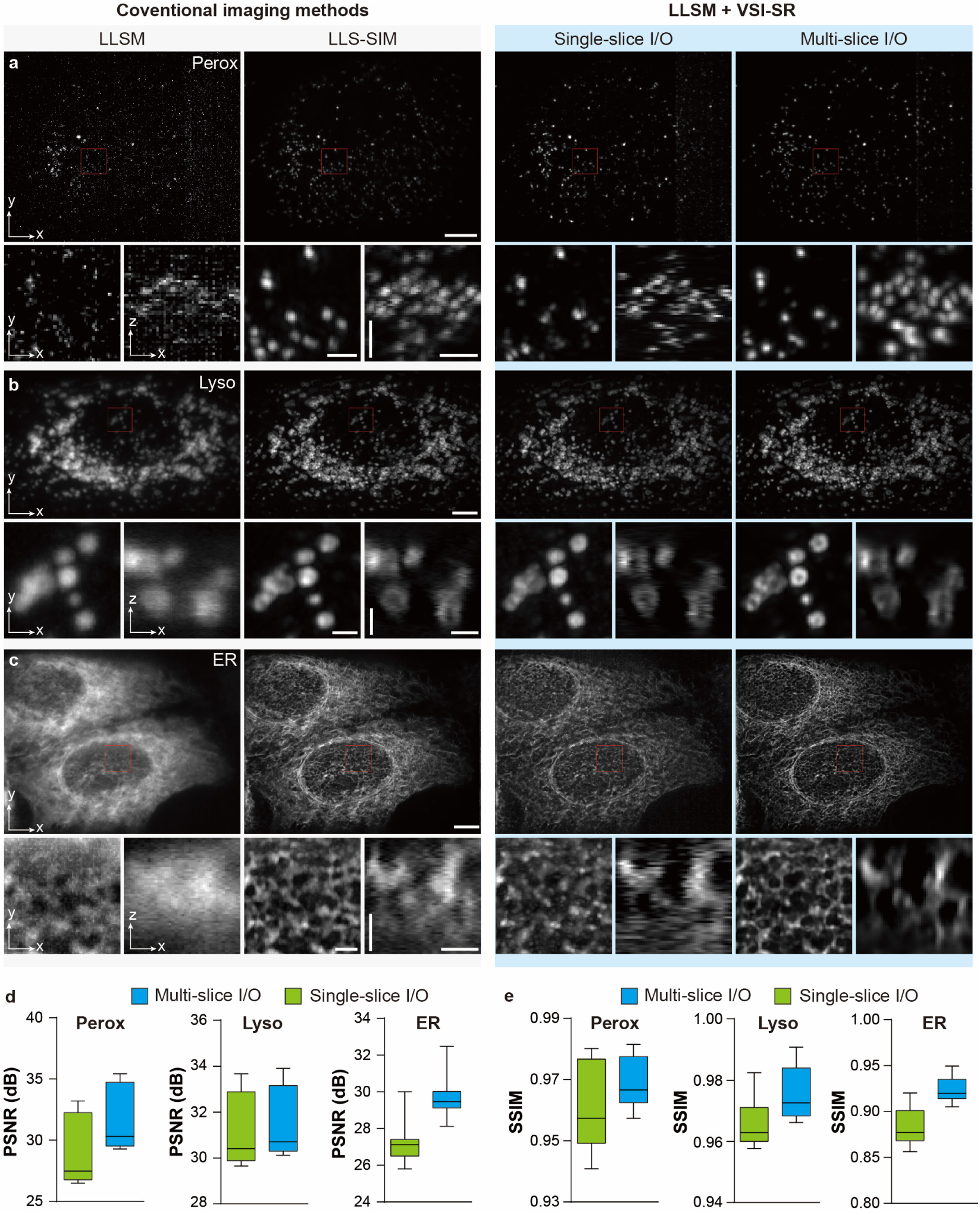
Comparison of VSI-SR models with inputs/outputs (IO) of single slices or multiple slices. **a-c,** Representative images (MIP) acquired and generated by LLSM, conventional LLS-SIM, and VSI-SR models with single-slice or multi-slices IO, which were trained by datasets of peroxisomes (a), lysosomes (b) and ER (c), respectively. Scale bar, 5 μm (full-FOV images), 1 μm (horizontal bars in magnified regions), and 3 μm (vertical bars in magnified regions). **d,e**, Box-and-whisker plots of the PSNR (d) and SSIM (e) for images reconstructed by VSI-SR models using dataset of peroxisome, ER and lysosomes (n=20 for each sample). Central line, medians; limits, 75% and 25%; whiskers, maximum and minimum.

**Extended Data Fig. 2.**
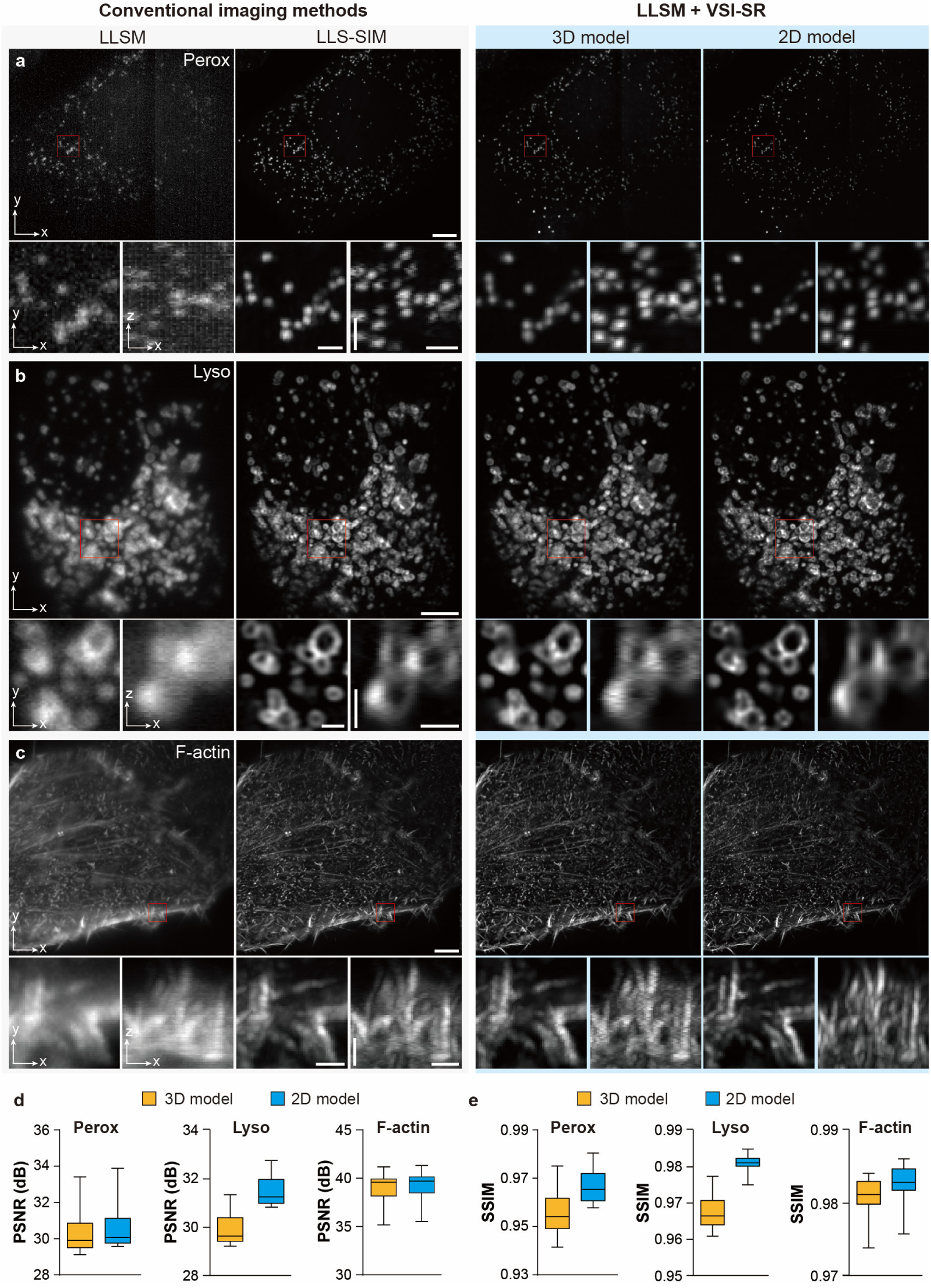
Comparison of VSI-SR models constituted with 2D and 3D convolutional architecture. **a-c,** Representative images (MIP) acquired and generated by LLSM, conventional LLS-SIM, and VSI-SR models constructed with 2D and 3D convolutional architectures, which were trained by dataset of peroxisomes (a), lysosomes (b), and F-actin (c), respectively. Scale bar, 5 μm (full-FOV images), 1 μm (horizontal bars in magnified regions), and 3 μm (vertical bars in magnified regions). **d**,**e**, Box-and-whisker plots of the PSNR (d) and SSIM (e) for images reconstructed by VSI-SR models (n=20 for each specimen). Central line, medians; limits, 75% and 25%; whiskers, maximum and minimum.

**Extended Data Fig. 3.**
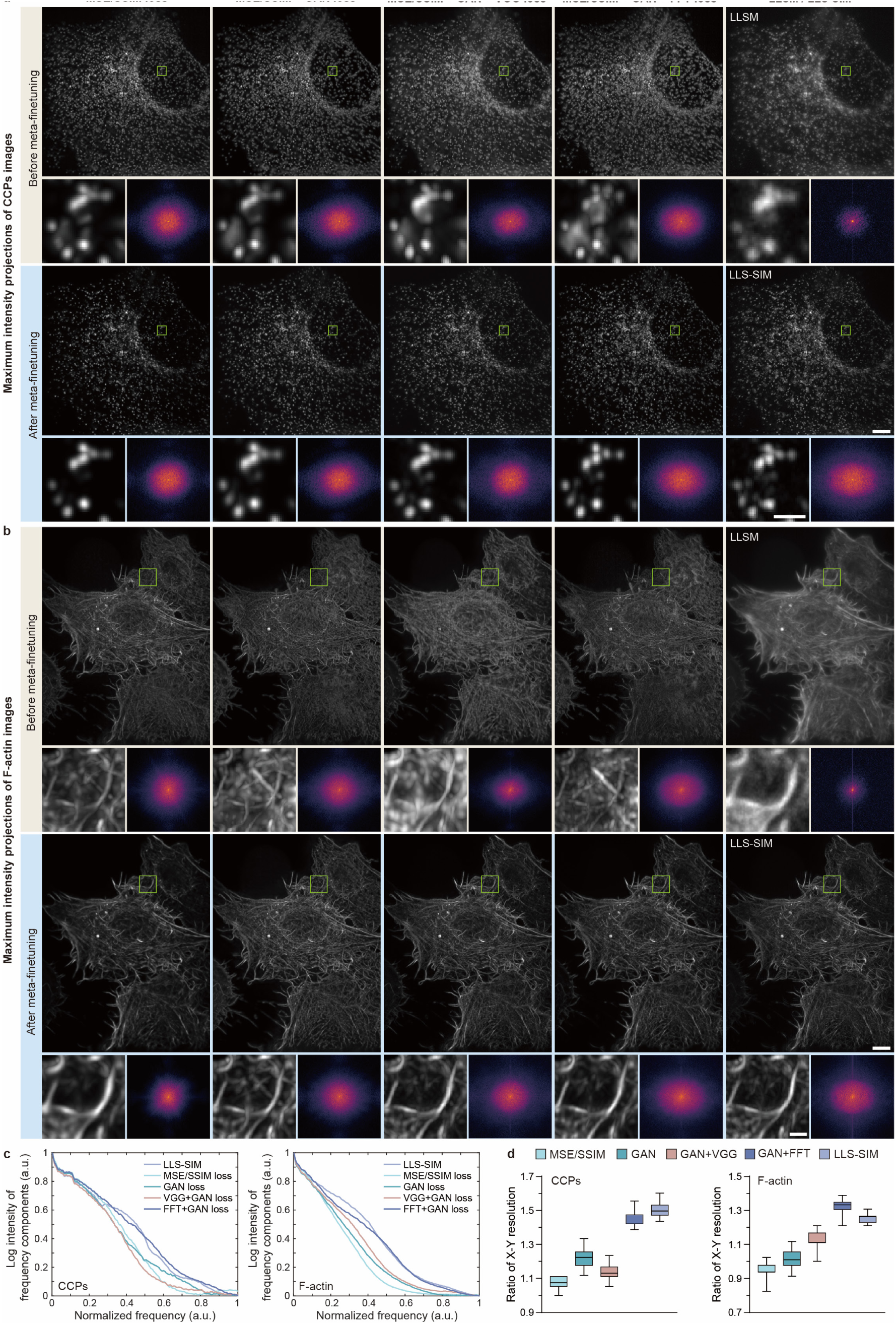
**Comparison of VSI-SR models trained with different loss functions. a**,**b**, Representative images (MIP) acquired and generated by LLSM, conventional LLS-SIM, and Meta-VSI-SR models trained with MSE+SSIM loss (first column), MSE+SSIM+GAN loss (second column), MSE+SSIM+VGG+GAN loss (third column), and MSE+SSIM+FFT+GAN loss (fourth column), which were finetuned by datasets of CCPs (a) and F-actin (b), respectively. Scale bar, 5 μm, 1 μm (magnified regions). **c**, Normalized log frequency intensity profiles of CCPs (left) and F-actin (right) images reconstructed by methods compared in a and b. Each curve was averaged by log frequency intensity profiles of 50 corresponding images. **d**, Box-and-whisker plots of the X-Y resolution ratio for SR images of CCPs and F-actin (n=50 for each type of samples) reconstructed by the methods compared in a and b. Central line, medians; limits, 75% and 25%; whiskers, maximum and minimum. Gamma value, 0.6 for all CCPs images and 0.7 for all F-actin images.

**Extended Data Fig. 4.**
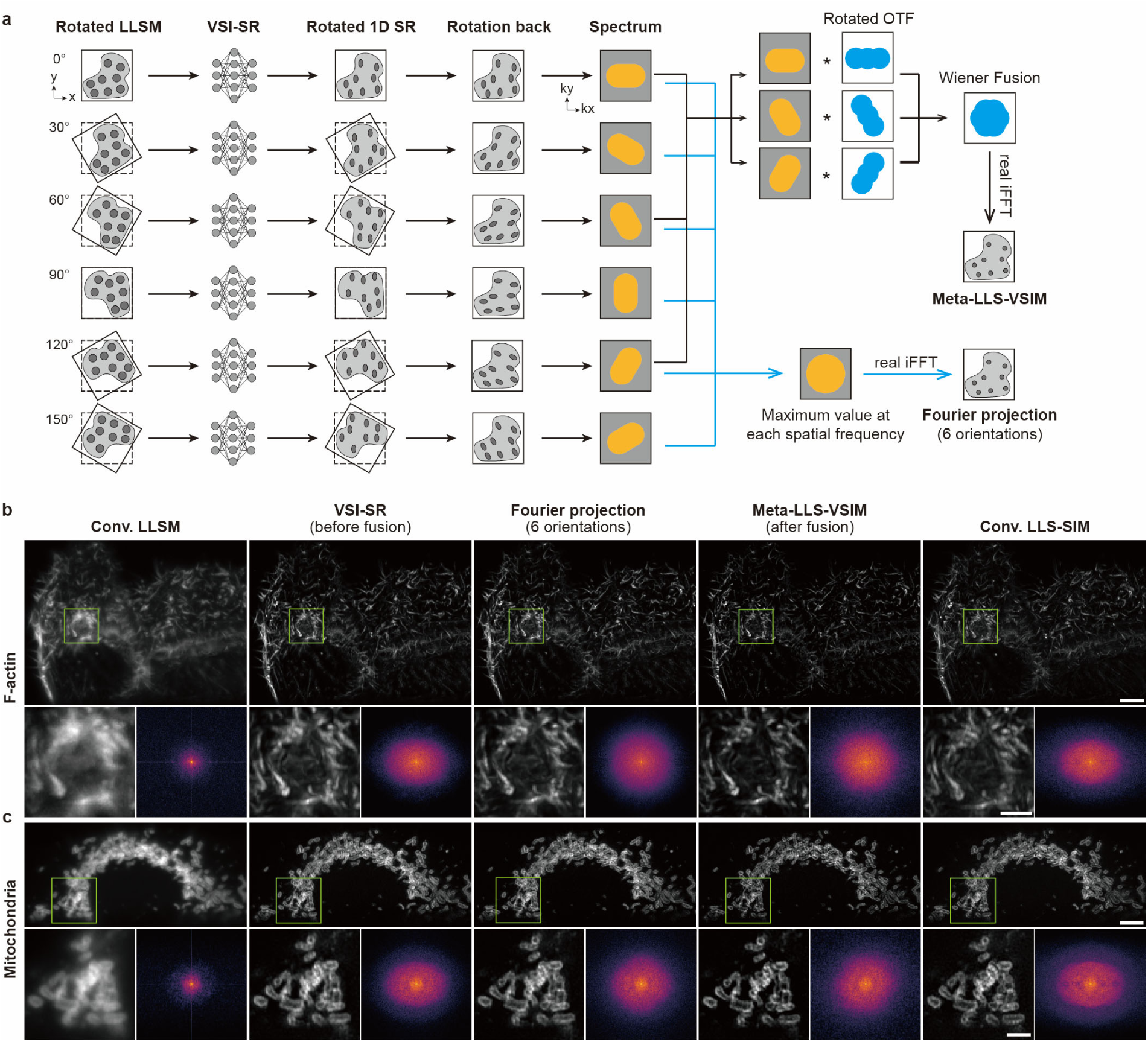
Comparison of different joint deconvolution strategies. **a,** Schematic of two different joint deconvolution strategies: the proposed generalized Wiener filter approach (upper panel) and the previous Fourier projection approach^19,28^ (lower panel). **b,c**, Representative LLSM images, anisotropic SR images directly inferred by VSI-SR models or imaged by conventional LLS-SIM, and laterally isotropic SR images reconstructed via the Fourier projection method and the generalized Wiener approach (Meta-LLS-VSIM) for F-actin and outer mitochondrial membrane. Scale bar, 5 μm (b, c), 2 μm (magnified regions of b, c).

**Extended Data Fig. 5.**
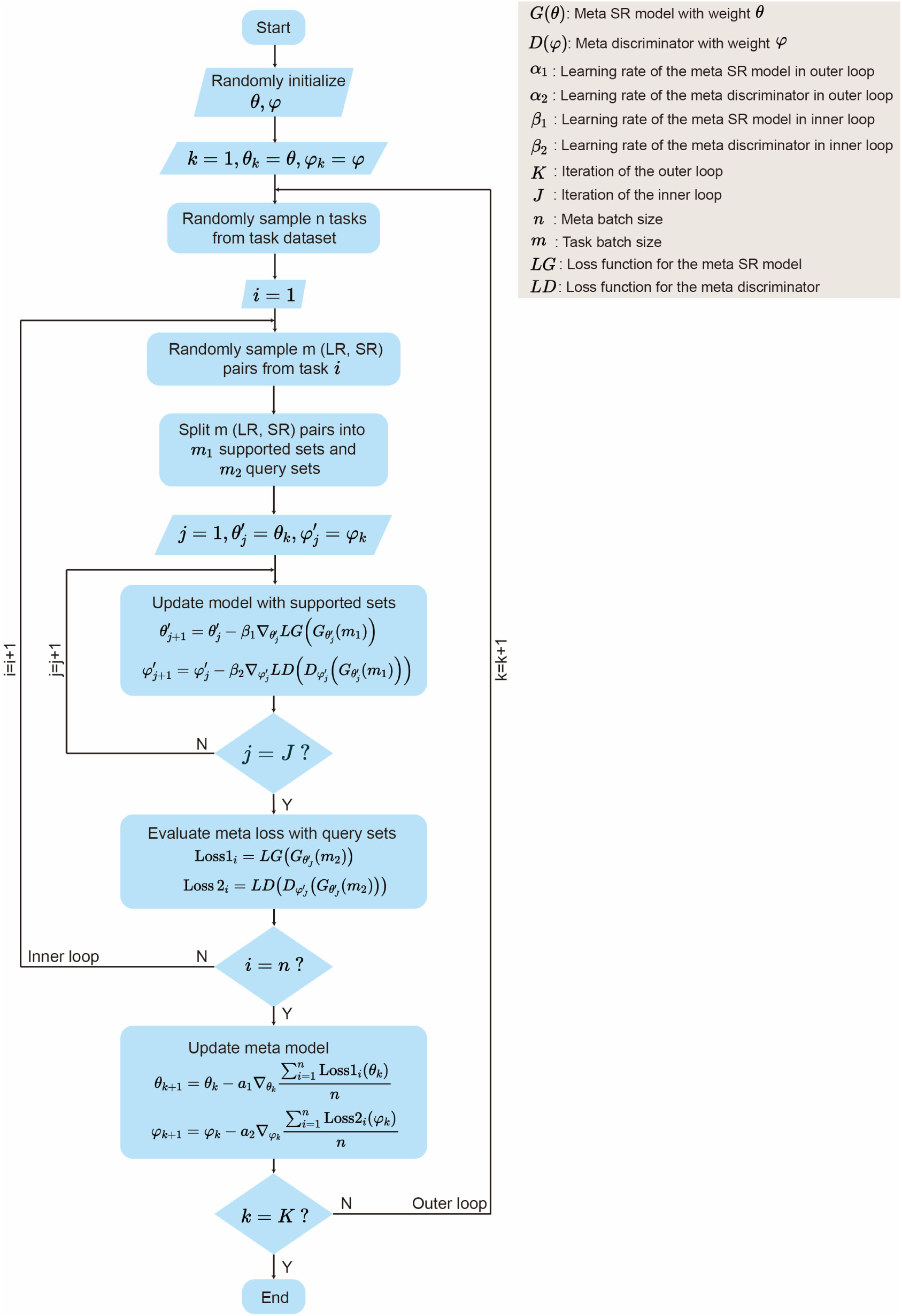
Meta-training workflow of the Meta-VSI-SR model.

**Extended Data Fig. 6.**
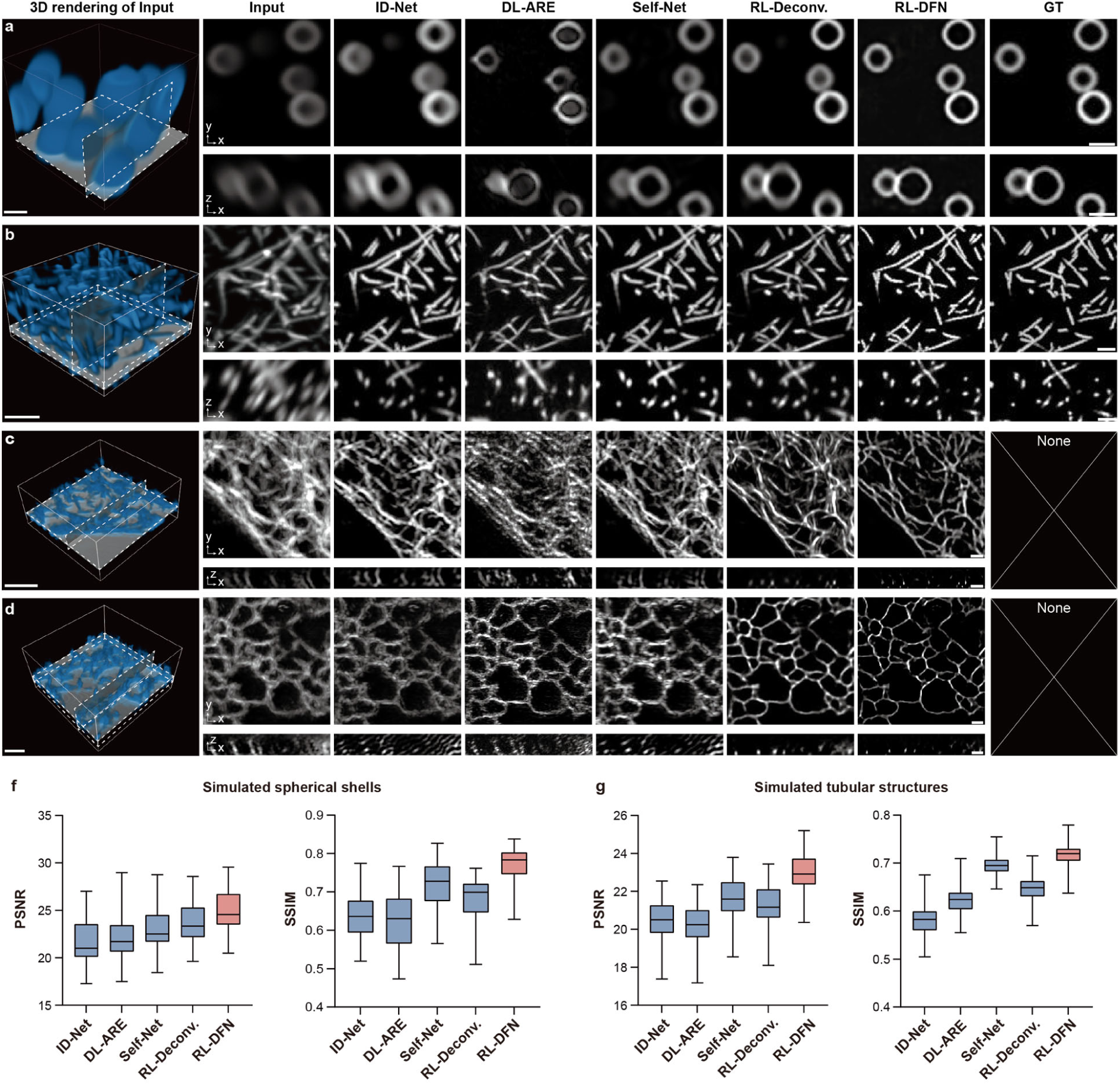
Comparison of RL-DFN with existing axial resolution enhancement methods. **a-d**, Representative x-y slices and x-z slices of simulated spherical shells (a), simulated tubular structures (b), experimentally acquired MTs (c), and experimentally acquired ER (d), reconstructed by multiple axial resolution enhancement methods, including ID-Net^18^, DL-ARE^19^, Self-Net^20^, multi-view RL deconvolution^26^, and the proposed RL-DFN. Anisotropic inputs, 3D rendering of the inputs, and the GT images are shown for reference. **f**,**g**, Statistical comparisons of ID-Net, DL-ARE, Self-Net, multi-view RL deconvolution, and RL-DFN in terms of PSNR and SSIM using simulated data of spherical shells (f) and tubular structures (g). Central line, medians; limits, 75% and 25%; whiskers, maximum and minimum. Scale bar, 2 μm (3D rendering images in a-d), and 1 μm (slice-images in a-d).

**Extended Data Fig. 7.**
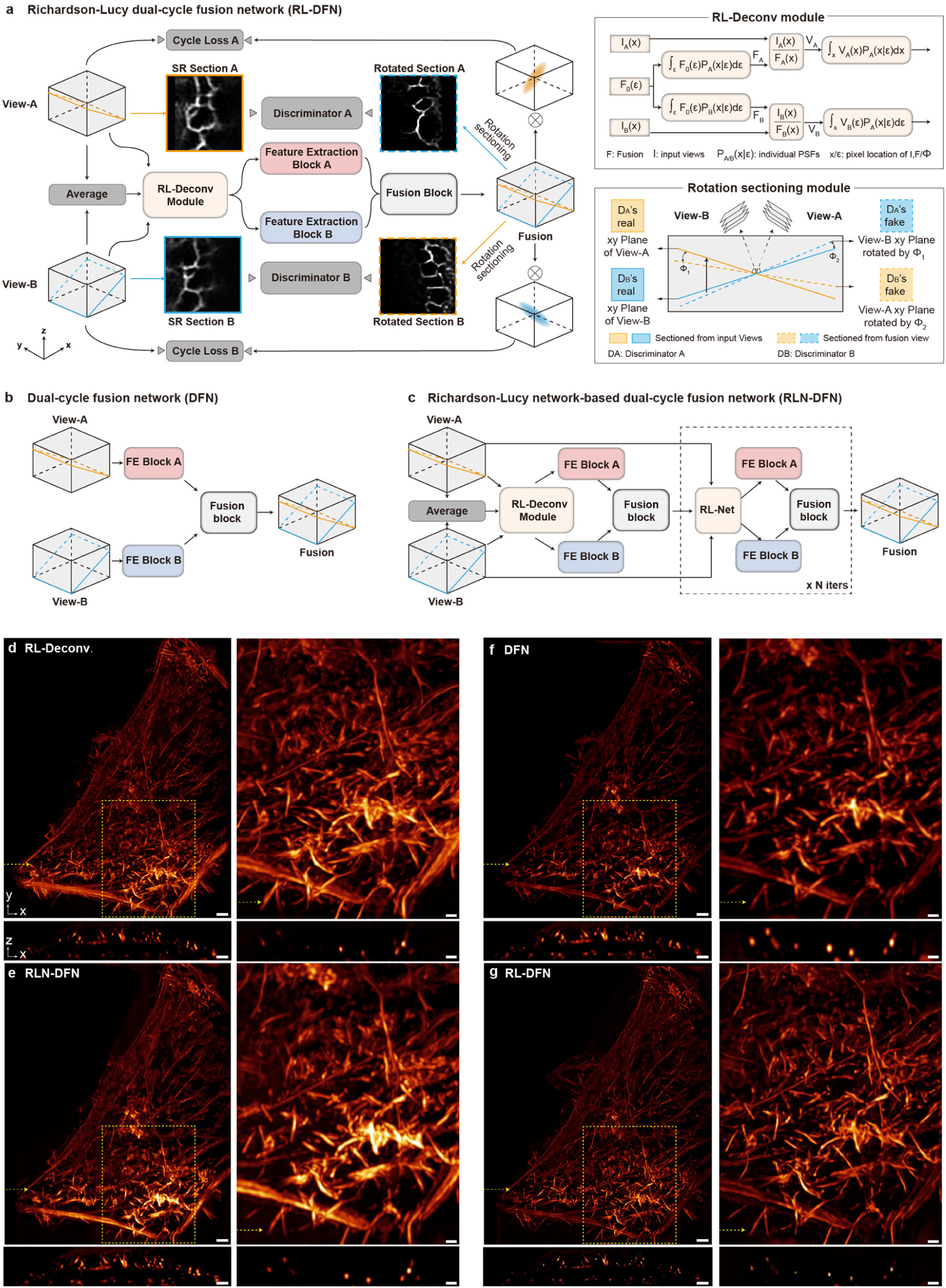
Comparison of RL-DFN with its variant architectures. **a-c**, Schematic of three compared network architectures, i.e., RL-DFN (a), DFN (b), and RLN-DFN (c). Details of RL deconvolution module and rotation sectioning module are schematically illustrated. **d**-**g**, Reconstruction results of F-actin images using multi-view RL deconvolution (d), RLN-DFN (e), DFN (f), and RL-DFN (g). x-y MIPs and two different x-z slices along the dashed arrows are shown. Scale bars, 3 μm (full FOV images in d-g) and 1 μm (magnified regions in d-g).

**Extended Data Fig. 8.**
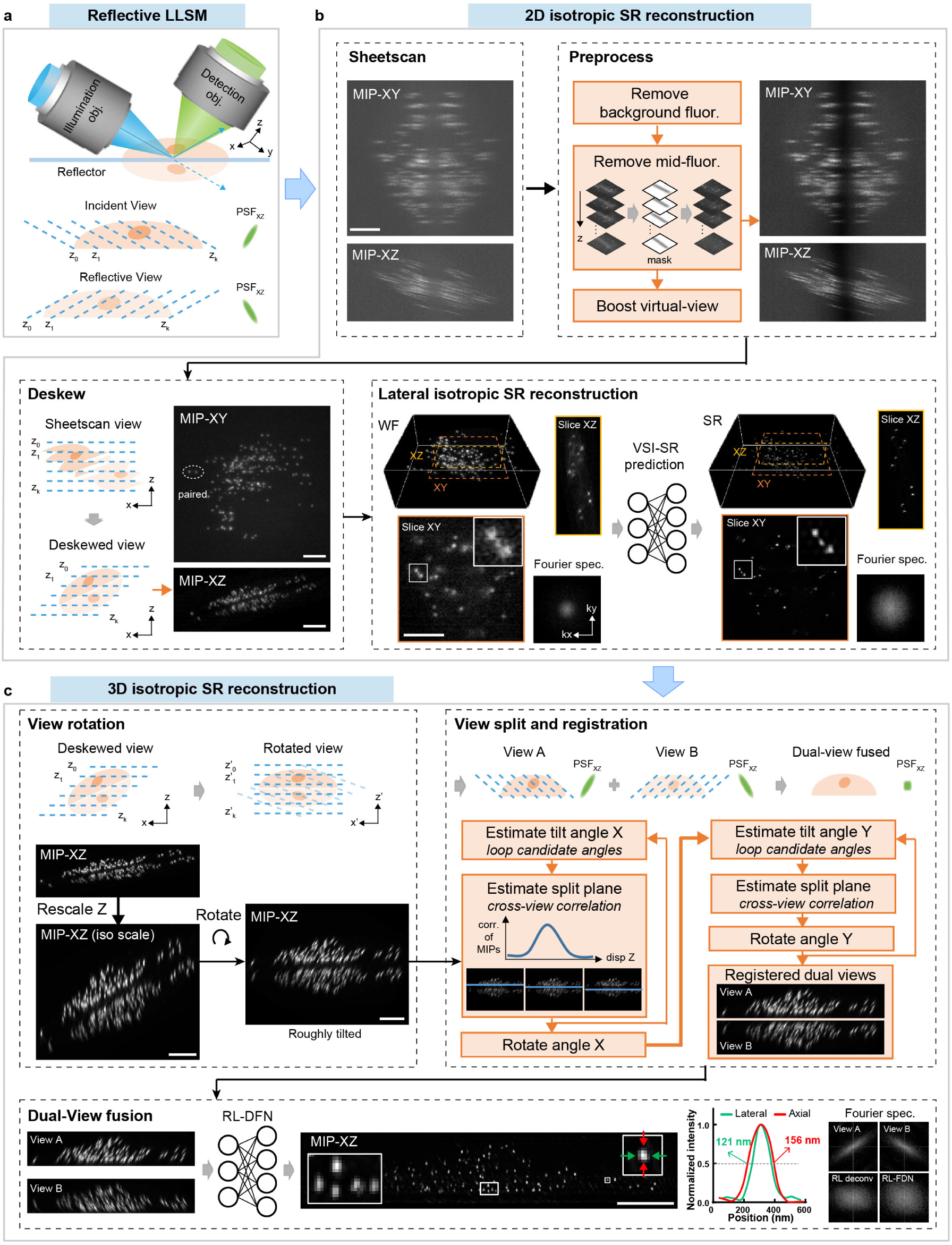
Pipeline of Meta-rLLS-VSIM reconstruction. **a**, Illustration of the imaging process of reflective lattice light-sheet microscopy (rLLSM). **b**, Step-by-step processing procedures for laterally isotropic SR reconstruction from wide-field input by VSI-SR models. **c**, Step-by-step processing procedures for isotropic 3D SR reconstruction by RL-DFN from the anisotropic SR views.

**Extended Data Fig. 9.**
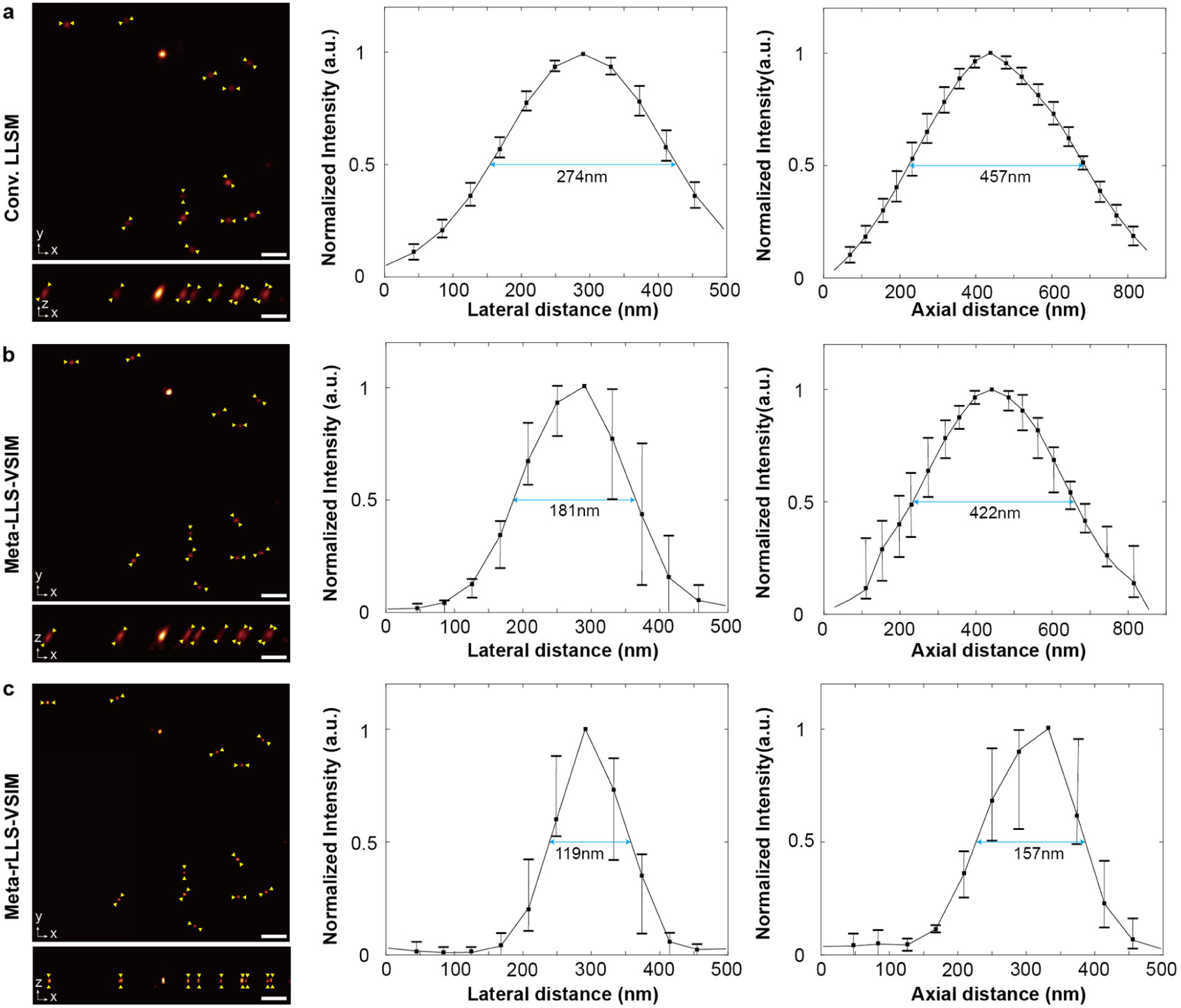
Resolution characterization for LLSM, Meta-LLS-VSIM, and Meta-rLLS-VSIM. **a-c**, LLSM (a), Meta-LLS-VSIM (b), and Meta-rLLS-VSIM (c) images (MIP) of beads shown in the x-y and x-z plane and corresponding line profiles indicated by yellow arrows in both lateral (n=12) and axial (n=9). The FWHM measured from the averaged profiles indicate that the resolution of Meta-rLLS-VSIM reaches 119 nm in lateral and 157 nm in axial. Central points, mean; whiskers, minimum and maximum. Scale bar, 1 μm.

**Extended Data Fig. 10.**
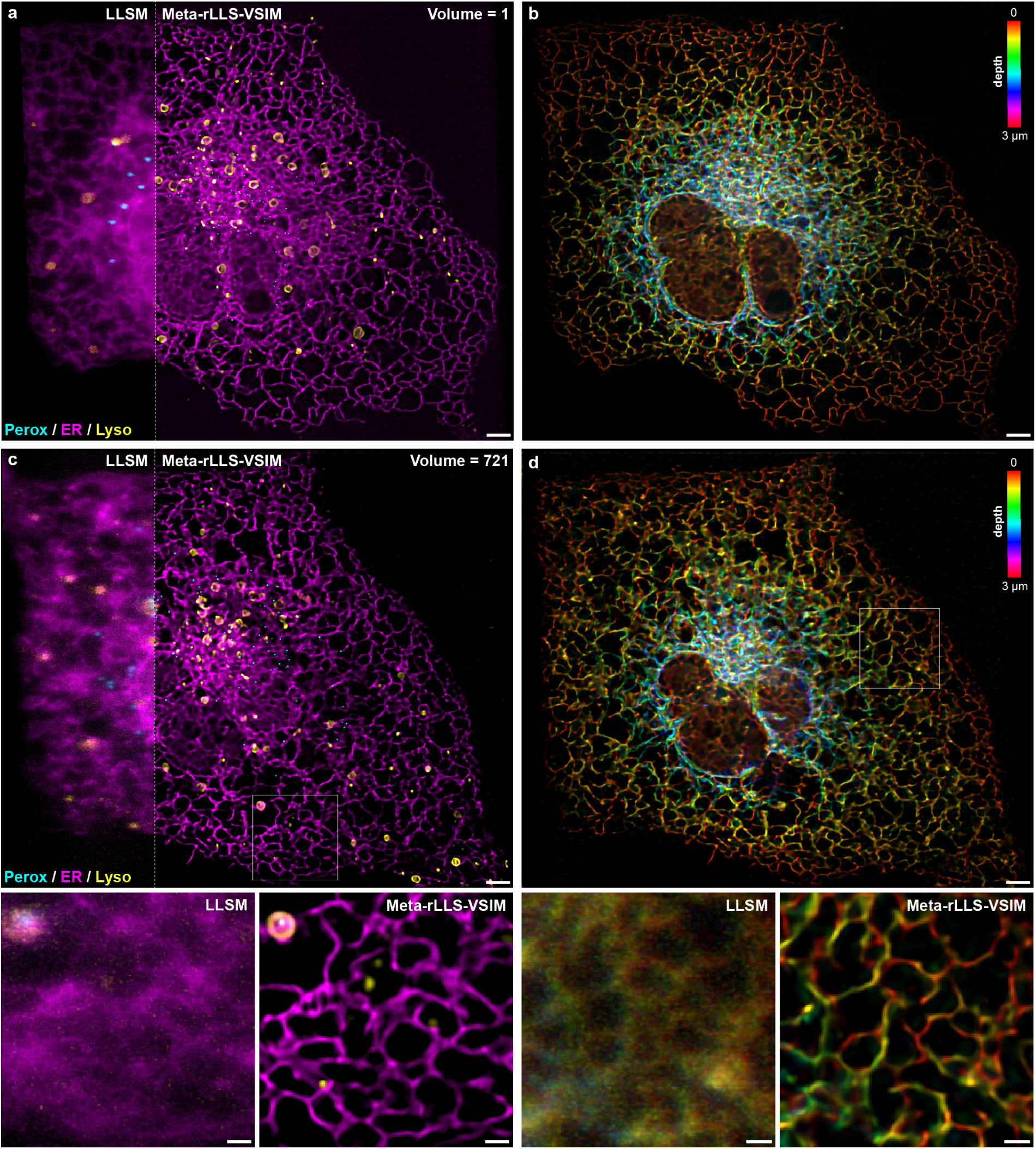
Three-color SR visualization of another Cos-7 cell by Meta-rLLS-VSIM. **a-d**, Three-color MIP (a) and z-coded visualization (b) of the first timepoint (a, b) and the last timepoint (c, d) from a time-lapse live recording of a Cos-7 cell expressing GFP-SKL, ER-mCherry, and Lyso-Halo by Meta-rLLS-VSIM (Supplementary Video 7). Scale bar, 3 μm (a-d), 1 μm (zoom-in regions of c and d).

## Notes

### Summary of Updates

Main manuscript updated and polished; Some main figures, supplementary figures, supplementary Videos revised; Author affiliations updated; Supplementary Notes updated.

