## Supplementary Materials for "Fast adaptive super-resolution lattice light-sheet microscopy for rapid, long-term, near-isotropic subcellular imaging"

**This PDF file includes:**

Supplementary Notes 1-6
Supplementary Figures 1-7
Supplementary Tables 1 and 2
Captions for Supplementary Videos 1-8
Supplementary References

#### Supplementary Notes

##### 1. Design of the VSI-SR model

The virtual structured illumination super-resolution (VSI-SR) model was constructed based on the residual channel attention network (RCAN)[1], which was initially developed for natural image super-resolution and then generalized to various low-level image processing tasks such as biological image restoration[2]. The objective function of the VSI-SR model was designed to minimize the structural differences between low-resolution (LR) and one-dimensional (1D) super-resolution (SR) images generated by lattice light-sheet structured illumination microscopy (LLS-SIM). Besides the two common types of penalty, mean square error (MSE) loss and structural similarity (SSIM) loss, which are applied to insure the output fidelity, we intended to introduce additional loss terms to alleviate the spectral bias[3] of deep learning-based SR (DLSR) models (i.e., DLSR models usually learn low-frequency modes faster and more robust than high-frequency modes[4]). Here, we examined three specialized loss terms and their combinations for enhancing the learning capability of high frequency components: the discriminative loss (or called GAN loss), which is typically used in the generative adversarial network (GAN)[5]; the fast Fourier transformation (FFT) loss that minimizes the difference between network output and GT in frequential space[6]; and the perceptual loss (or called VGG loss), which enhances the perceptual quality by calculating losses between deep output and target features extracted by a pretrained VGG-16 model[7]. Specifically, we trained VSI-SR models for two types of biological structures, clathrin-coated-pits (CCPs) and F-actin, with four different combined loss functions that were constituted by (i) MSE and SSIM loss, (ii) MSE, SSIM and GAN loss, (iii) MSE, SSIM, GAN and VGG loss, and (iv) MSE, SSIM, GAN, and FFT loss, respectively. We found that the loss functions consisting of MSE, SSIM, GAN and FFT loss outperformed other combinations in extending resolution along the SR orientation of the LLS-SIM targets, and the VSI-SR model trained with such a loss function was able to ideally restore the anisotropic power spectrums after meta-finetuning for both biological structures (Extended Data Fig. 3).

After determining the objective function of the VSI-SR model, the next question is how to fully utilize the spatial continuity in z-axis of volumetric lattice light-sheet microscopy (LLSM) and LLS-SIM images in the network training and inference procedures. Our intuitive thinking is that SR models with multi-slice inputs and outputs can take advantage of spatial continuity in z-axis of both LLSM inputs and

LLS-SIM targets, and produce SR images with higher fidelity than those with single-slice input and output. To verify this speculation, we trained two VSI-SR models with the identical network architecture constituted by 2D convolutional layers and channel attention mechanism (Supplementary Fig. 1), while differing in numbers of input and output channels: (i) a 2D VSI-SR model with multi-slice inputs and outputs, which considered volumetric LLSM as multi-channel data, and inferred middle three x-y SR slices from the corresponding seven consecutive x-y LR slices; (ii) a 2D VSI-SR model with single-slice input and output, which inferred a single x-y SR slice from the corresponding x-y LR slice. For each input/output configuration, three models were independently trained with datasets of peroxisomes (Perox), lysosomes (Lyso), and ER, respectively. We found that the SR images reconstructed by VSI-SR models with multiple input and output slices presented better axial continuity (xz-displays in Extended Data Fig. 1a-c), finer structures (xy-displays in Extended Data Fig. 1a-c), and higher fidelity quantified by PSNR and SSIM (Extended Data Fig. 1d, e) than those inferred by VSI-SR models with single-slice input and output.

On the other hand, neural network models constructed based on 3D convolutional layers could also effectively extract and aggregate volumetric features of its input data[2]. To explore whether 3D or 2D architecture is better for the VSI-SR task, we modified the previous 2D VSI-SR model into a 3D version. Compared with the 2D VSI-SR model, the major reconfigurations for the 3D scenario included: (i) all 2D convolutional layers were replaced by 3D convolutional layers for 3D feature extraction; (ii) adopting volumetric data as inputs and outputs, and the channel of the input and output was set to 1; (iii) we maintained the same network depth for 2D and 3D models but decreased the number of feature channels in 3D network, thereby the total parameters for both models were ~1.4 million, yielding a relatively fair comparison for these two network architectures. Similarly, for 2D and 3D models each, we trained three models separately using datasets of Perox, Lyso, and F-actin. The 3D models typically required 32 hours for 200,000 mini batch iterations in traning. In contrast, the 2D VSI-SR models only took 17 hours for the same anount of iterations, while required less GPU memory than the 3D version. After well-trained, we found that the 2D VSI-SR models could generate SR images of better spatial resolution and axial continuity with higher PSNR and SSIM than 3D models (Extended Data Fig. 2), implying that to view volumetric lattice light-sheet data as multiple 2D slices distributed in the channel dimension of 2D convolutional neural network is the optimal solution to fully utilize the axial continuity and obtain the best output spatial resolution and SR fidelity in our case.

#### 2. Evaluation of VSI-SR model

##### A. Validation of laterally isotropic SR reconstruction on TIRF-SIM data

The incapability to acquire isotropic 2D SR images by using the LLSM-SIM makes the validation of lateral isotropic SR reconstruction capability of the VSI-SR method missing ground-truth. To solidly validate the performance of the VSI-SR model and the proposed laterally isotropic reconstruction scheme, here we applied the VSI-SR method to process the diffraction-limited total internal reflective fluorescence (TIRF) images acquired by our home-built TIRF-SIM system[8], where the corresponding TIRF-SIM images with isotropic super resolution could serve as the reference.

In the TIRF-SIM system, sinusoidal structured illuminations along three orientations were sequentially generated and projected on the sample, resulting in uniform spatial frequency extension within the 2D lateral space[8,9]. Specifically, a 1.49 NA objective was adopted for fluorescence excitation and collection to achieve ~200 nm resolution. And the isotropic SR TIRF-SIM images could be reconstructed from three-orientation-illuminated raw acquisitions using the conventional SIM algorithm with resolution enhanced by up to 2-fold. The reconstructed TIRF-SIM images were taken as the ground-truth to evaluate the fidelity and resolution improvement of VSI-SR models in the following evaluation.

To emulate the process of VSI-SR reconstruction applied to the LLS-SIM system, we intentionally picked up the acquired raw TIRF-SIM data that were illuminated with sinusoidal patterns in a single orientation $\phi_0$. By applying the conventional SIM reconstruction algorithm on the picked data only, we generated anisotropic SR images with resolution enhancement only along the vertical dimension (analog to the SR data acquired by LLS-SIM). A fraction of the paired TIRF and anisotropic SR images were used for VSI-SR network training. The remaining TIRF data were used as the testing set, undergoing the three-orientation network inference and Wiener fusion process as described in Fig. 1c to generate laterally isotropic SR predictions, noted as TIRF+VSI-SR in the following section and Supplementary Fig. 3.

The comparison between TIRF+VSI-SR and TIRF-SIM images was conducted both visually and quantitatively. As shown in Supplementary Fig. 3, the reconstructed images of endoplasmic reticulum (ER) and microtubules by using TIRF imaging, VSI-SI prediction from TIRF images, and TIRF-SIM are presented for visualization. The Fourier spectrums are also provided by applying a Fourier transform on the corresponding spatial images. The results reveal an obvious structure refinement and OTF extension of TIRF+VSI-SR, which is comparable to the performance of TIRF-SIM. For the ER images, intensity profiles of frequency spectrums along two orientations $\phi_1$ and $\phi_2$ that are $60^\circ$ and $120^\circ$ clockwise rotated from $\phi_0$ were measured to demonstrate the spatial resolution improvement of the VSI-SR reconstruction. For the microtubule images, the line profiles of two closely spaced filaments were measured to validate the super-resolution capability of VSI-SR, with TIRF-SIM as the GT SR reference. In addition, the Fourier ring correlation (FRC) analysis[10] was conducted to numerically compare the spatial resolution of TIRF, TIRF+VSI-SR, and TIRF-SIM images.

**B. Comparison of the VSI-SR method and cross-modality training scheme**

Deep learning techniques aiming at predicting SR images with isotropic spatial resolution from their low-resolution (LR) counterparts have been frequently demonstrated in recent years, among which the supervised schemes commonly provide satisfactory fidelity and robustness[11,12]. A critical necessitate of these supervised approaches is the large sets of LR-SR image pairs for network training. However, collecting such isotropic SR images could be impracticable for some imaging setups, for example, the LLSM/LLS-SIM that is widely used for long-term live-cell volumetric imaging[13], because it is designed to generate structured illumination along one single orientation, yielding only one-dimensional super-resolution capability. This hinders us from directly training an end-to-end network to achieve single-image based 2D isotropic SR reconstruction. To address this issue, one feasible solution is to apply a cross-modality training scheme, which means that the LR-SR paired images for training are captured by other SR microscopes, e.g., TIRF-SIM, and then the trained model is applied to process images of the target imaging modality, e.g., LLSM, in a cross-modality manner.

As a demonstration of the cross-modality training scheme, we acquired 2D isotropic SR images of F-actin and ER by using TIRF-SIM to provide SR labels for single image super-resolution (SISR) model training. The corresponding LR inputs were generated by averaging the raw SIM images used for TIRF-SIM reconstruction. We showed that the SISR model trained with such data performed well when applied to TIRF images acquired by the same TIRF-SIM system (Supplementary Fig. 4a-c). Nevertheless, we found when these models were used to predict isotropic SR images from wide-field data acquired by the sheet-scan mode of the LLSM system, the inevitable differences between two imaging modalities, i.e., the domain shift problem, would cause artifacts during the cross-modality prediction process. The domain shift problem mainly arises from the different imaging process principles of the two modalities: i) LLSM illuminates a thin slice of sample using a light-sheet with a thickness of 300~500 nm determined by the excitation numerical aperture (NA), while the TIRF-SIM in its inverted design only works for 2D imaging scenarios that illuminated a thin slice of ~100 nm above the coverslip; ii) TIRF-SIM setup acquires single slice data in a parallel view with the coverslip, while the LLSM setup in our experiments acquires data in a tilted view by ~30°; iii) different imaging systems usually possess different amplification factors with a different pixel size for the acquired images. These factors will lead to cross-modality differences in terms of the sample size, spatial distribution of subcellular structure, and signal-to-background contrast of captured images. Supplementary Fig. 4g-j shows that if we directly applied the SISR networks that were trained on TIRF/TIRF-SIM data pairs to process LLSM data, background fuzziness and ring artifacts would occur, which contaminated the fine structure of the specimen. In the contrary, using the proposed VSI-SR scheme where a single-orientated anisotropic SR model was trained on the LLSM/LLS-SIM data pairs that can be feasibly acquired on the same LLSM setup with lattice structured illumination, we could avoid the domain shift problem, resulting in artifact-free isotropic SR reconstructions and better signal-to-background contrast compared with the cross-modality training scheme (Supplementary Fig. 4h, j).

#### 3. Design of Meta-VSI-SR

Meta-learning (or called learning to learn) has become one of the hottest topics in artificial intelligence, which tries to make artificial agents learn and adapt quickly to a new task using only a few examples. There are three types of mainstream meta-learning algorithms, including (i) metric-based meta-learning, which aims to learn a metric space where learning is quick with a few examples[14-16]; (ii) memory network-based meta-learning, which acts as a trained recurrent neural network that allows encoding and retrieval of episodic information[17-19]; (iii) optimization-based meta-learning, that directly optimizes the standard gradient descent rule to train a meta learner with fast adaptation capacity to new tasks[20-23]. Among these algorithms, model-agnostic meta-learning (MAML) has exhibited a huge impact on the research community since it can be directly utilized to any neural network models trained with a gradient descent procedure, and shows state-of-the-art performance[23].

The main idea of MAML is to train a set of neural network's initial parameters such that the network has optimal performance on a new task after being updated through one or more gradient steps with a small amount of data from the new task. There are several MAML variants, including (i) MAML with second order derivative terms, which considers second order derivatives in the meta training stage[23]; (ii) MAML with first order derivative approximation, which omits all second derivatives in the meta training stage[23]; (iii) MAML++, a hybrid version of first order MAML and second order MAML to cope with the instability of MAML training[24]; (iv) Reptile, a more simplified first order MAML[25]. Since we empirically found that the performance of first order MAML is nearly the same as that obtained by second order MAML in our task, we employed the first order MAML scheme to reduce computational expenses and speeding up our training process. It is noted that we introduced GAN scheme into MAML for better extraction of anisotropic high-frequency information, therefore, compared with the algorithm of original MAML, our MAML-based meta-learning algorithm trained two meta models (i.e. the meta VSI-SR model and the meta discriminator) simultaneously (Extended Data Fig. 5). The meta VSI-SR model and the meta discriminator were updated three times and once in the inner loop of MAML, respectively. During the training process, they competed with each other, and finally reached an equilibrium state.

Since MAML requires a set of diverse tasks for building an internal representation that is broadly suitable for many tasks, the following question is how to divide our biological datasets into a series of diverse task datasets, where each task corresponds to a specified SR problem. Our biological datasets were acquired via a home-built LLS-SIM system, which consisted of a large number of biological specimens with diverse structures and different light intensities, hence, a straightforward thinking was that task division could be conducted based on biological structures. In addition, we regarded the SR inference for LLSM images with different light intensities, i.e. signal-to-noise ratio (SNR), as different tasks because of the following two reasons: (i) similar to the structural complexity that is determined by the sample, the SNR of input images is another vital factor that substantially influences the final SR performance of VSI-SR models; (ii) the meta VSI-SR model trained with such datasets could be then finetuned to adapt to any specific specimen and SNR, yielding the optimal adaptation to the specific application condition. To this end, we established the datasets for meta-learning that contain 20 diverse tasks from 10 biological specimens with low (about 200 average photon counts for each raw frame) and high (more than 1000 average photon counts for each raw frame) fluorescence levels in this work. Each task-specific dataset represented low or high SNR LLSM images of a certain biological structure. Moreover, it needs to be emphasized that instead of generating more task datasets by introducing various fluorescence levels, we only considered low or high SNR tasks to enhance the discrimination among tasks, since we found that containing too many similar tasks in the meta training datasets would degrade the final meta model into a pretrained model, i.e., the model trained by standard stochastic gradient descent with the same sort of training data.

#### 4. Design of RL-DFN

The RL-DFN is developed to fuse the dual-view volumetric images acquired by the reflective lattice light-sheet microscopy (rLLSM) in a self-supervised manner. To devise the self-supervised objective function of RL-DFN, there are three key points to be considered: first, the loss function should impose a constraint that could maintain the structural consistency between the network output and its dual-view inputs; second, there should be a loss term that regulates the spatial resolution to be isotropic in 3D space; third, the prior knowledge of the sample and data should be fully exploited to rationalize the training and inference procedures. Bearing in mind these analyses, we designed a dual cycle-consistent adversarial loss function inspired by the conditional GAN (cGAN)[26] and cycle-consistent adversarial networks (cycleGAN)[27], which consists of three terms: the cycle loss $L_{cycle}$, discriminative loss $L_{Disc}$, and total variational (TV) regularization $R_{TV}$.

The cycle loss $L_{cycle}$ minimizes the difference between anisotropic input volumes and the network prediction degraded by the corresponding point spread functions (PSFs) of each view. The degradation operation simulates the physical imaging process of rLLSM; hence the cycle loss leads the model to learn the reverse process, which is similar to the Bayesian-based multi-view deconvolution[28]. It is noteworthy that different from the cycleGAN that degrades or transforms the generator's outputs by another deep neural network, here we adopted the deterministic dual-view PSF blurring strategy, which we empirically found much more computationally efficient and stable.

Next, the discriminative loss $L_{Disc}$ is designed to teach the RL-DFN model to enhance the resolution isotropy of its output volume after fusion. Assuming a data volume with isotropic resolution in 3D, its 2D sections from any arbitrary view should be isotropically resolved. Therefore, the 3D resolution isotropy could be evaluated and regulated by its 2D sections. Inspired by this, we assign two discriminators (for view-A and view-B respectively) to distinguish whether a 2D section is sampled from the original input volume along its lateral dimensions (i.e., with isotropic resolution) or from the output of RL-DFN. To fool the discriminators, the RL-DFN model endeavors to make any 2D sections of its output be isotropically resolved, finally resulting in 3D isotropic resolution of the whole output volume. In addition, since some subcellular structures have anisotropic distributions in lateral and axial dimensions, i.e., the cytoskeleton in adherent cells, there exists a difference in spatial morphology between sections from different angles, which may affect the discriminators' judgement of resolution during training. To tackle with this issue, we limited the sampling angle of the 2D sections to a certain range of [-π/6, π/6] around the orientations perpendicular to the optical axis of view-A and view-B (schematically shown in Extended Data Fig. 7a), to prevent the discriminators from making judgement on sections with obvious content differences in distribution. Besides the cycle loss and discriminative loss, we additionally introduced a TV regularization term into the overall objective function of RL-DFN, which could utilize the structural continuity in both lateral and axial of the biological specimen to regulate the training procedure of the neural network model.

After defining the objective function, we settled on a functional outline of RL-DFN: (i) dual-view volumes acquired by the rLLSM system are fused into one image volume with isotropic resolution in all dimensions; (ii) the cycle loss is calculated between the PSF-degraded output volume and dual-view input as a cycle-consistency constraint, which is a fundamental loss term to learn the inverse imaging model of rLLSM; (iii) the discriminative loss takes a step further to prompt the volumetric resolution isotropy of the output; (iv) and the TV term imposes a spatial continuity constraint and rationalize the final network output based on the property of the specimens. Within this framework, one remaining question is what is the most appropriate neural network architecture to conduct the forward fusion process. The simplest solution is to concatenate the dual-view volumes along the channel dimension and fuse them with a 3D neural network. To examine this idea, we devised a neural network architecture, dubbed as deep fusion network (DFN, see schematic illustration in Extended Data Fig. 7b), that first extracted deep features from the dual-view inputs by two 3D feature extraction blocks, then concatenated the extracted features along the channel dimension, and finally generated the fusion volume via a fusion block. It turned out that the simple DFN could recover the biology structures and enhance the lateral resolution, but the axial resolution of the fusion result was much worse than the lateral one, indicating that the simple DFN is insufficient to achieve isotropic SR reconstruction even with the elaborately designed objective function (Extended Data Fig. 7f).

We speculated that the invalidation of DFN was primarily due to the direct dual-view fusion by simple concatenation and implicit convolutional layers could not capture the image formation process, which has been proved to be of vital importance for accurate image restoration of fluorescence images[4,29]. To effectively take advantage of the prior knowledge about the imaging forward model in the fusion process, we introduced the multi-view Richardson–Lucy (RL) deconvolution into the neural network architecture design to provide a physics-consistent feature fusion scheme in the DFN, thus we named this model as RL-DFN. Specifically, in RL-DFN, the dual-view input volumes were first processed by a single iteration of RL deconvolution using unmatched back projectors[30] to obtain two initial estimations of the fusion volume, then two feature extraction blocks were used to extract deep feature maps from them. Instead of concatenating the extracted feature channels, we conducted the Hadmard product following the Bayesian-based derivation[28] to perform the feature fusion, and then the merged features went through a 3D fusion block to generate the final isotropic SR volume. The overall architecture of the proposed RL-DFN is depicted in Fig. 4b, Supplementary Fig. 6 and Extended Data Fig. 7a. We demonstrated that the incorporation of RL deconvolution pre-processing and Bayesian-based feature fusion into the neural network permitted us to feed the prior-knowledge about the physical process and parameters of microscopy into both training and inference processes, and substantially rationalized data forward propagation within RL-DFN. Moreover, we tried to upgrade the architecture of RL-DFN by replacing the 3D convolutional blocks with RL network (RLN)[29] (corresponding model was dubbed as RLN-DFN) to further improve the SR performance. However, we empirically found that the introduction of RLN destabilized the training process and reduced the output resolution. As a consequence, we reached out the optimal RL-DFN architecture that achieved the best dual-view fusion performance and isotropic SR resolution restoration compared with conventional multi-view RL deconvolution, DFN, RLN-DFN, and other state-of-the-art self-learning-based isotropic reconstruction methods (Extended Data Figs. 6 and 7d-g).

#### 5. Simulation of 3D LLSM data

Two sets of simulated 3D LLSM datasets were computationally generated resembling different categories of biological specimens to evaluate and compare the 3D isotropic SR reconstruction performance of various dual-view fusion and isotropic reconstruction methods in Fig. 4c and Extended Data Fig. 6.

(1) Generation of 3D distribution of sphere structures. We simulated a 3D ellipsoid/semi-ellipsoid consisting of multiple randomly located small hollow spheres with random radius and intensity within a predefined range in Matlab to emulate the granular structures in the cell. The ground-truth volume was generated at a much higher resolution than the diffraction limit. In Extended Data Fig. 6, an ellipsoid of 128×128×128-voxels consisting of 300 hollow spheres with radius of 5 to 8-pixels and normalized intensity of 0.4 to 1.0 was generated.

(2) Generation of 3D distribution of tubular structures. A 3D volume consisting of twisted filaments was generated in Python to emulate the tubular structures in live cells such as microtubules. The simulation data was generated by first drawing randomly located straight lines in 3D space determined by two points, then applying elastic grid-based deformation in 3D using the open-source module (https://github.com/gvtulder/elasticdeform) to the volume.

For both simulation datasets, low-resolution slices from LLSM detection view and its virtual view were generated from simulated GT volume by applying convolution with corresponding tilted 3D PSFs, adding background, Poisson and Gaussian noise, and rescaling in z-axis to the sampling axial resolution in real experiments. 2D and 3D super-resolution references were generated by similar procedure but with the theoretical 2D-SR and 3D-SR PSFs.

#### 6. Detailed reconstruction pipeline of Meta-rLLS-VSIM

The proposed Meta-rLLS-VSIM method achieves 3D isotropic SR reconstruction from a single frame of wide-field capture. The process of 3D isotropic SR reconstruction is divided into three subsequent steps depicted in Extended Data Fig. 8 and explained below.

##### A. Data collection by reflective lattice light-sheet microscopy

The first step is to acquire a dual-view image volume of the sample by using the reflective LLSM (rLLSM) system. The sample is illuminated by the incident and the reflective light, resulting in a normal and a virtual view simultaneously collected by the detection objective. The two views both have anisotropic spatial resolution, with the PSF elongated along the detection axis, nearly orthogonal to each other.

##### B. 2D isotropic super-resolution reconstruction

The next step is to achieve 2D isotropic SR reconstruction for each view of the rLLSM imaging. This process consists of several steps:

**B.1 Background removal:** The maximum intensity projection (MIP) of the raw sheet-scan stack by rLLSM is presented as an example, indicating the strong background fluorescence. Specifically, there exist two sources of background fluorescence: (i) a global background signal caused by the out-of-focus emission and camera offsets, (ii) and the signal in the middle of two views caused by the reflective mirror. Therefore, a two-step background removal is applied. We first apply an offset removal to suppress the global background (we empirically set an intensity value of 100). Then, we estimate the reflector-induced fluorescence and mask out the mid-fluorescence. To do so, the image stack is divided into several groups of slices, and a stripe pattern with a 1D Gaussian profile is iteratively rotated and shifted to find the optimal match with the average XY projection of the dual-view sheet-scan image for each group. The estimated mask patterns for all groups are further regressed along the Z-axis to produce the 3D mask pattern, which is used to attenuate the mid-fluorescence in the sheet-scan stack. Moreover, to compensate for the intensity attenuation in the virtual view, we calculate the total intensity for each view and boost the intensity of the virtual view by the intensity ratio.

**B.2 Deskew the sheet-scan stack:** During the sheet-scan imaging process, the sample is translated horizontally along the coverslip plane, making the captured slices misaligned in the X-axis. To preserve the spatial continuity of the sample structure, we deskew the stack into the deskewed view. The MIPs in XY and XZ planes are presented, showing a tilted but visual-friendly view of the sample.

**B.3 2D SR prediction by VSI-SR and multi-orientation combination:** After the pre-processing, the deskewed stack is sent to the VSI-SR model for super-resolution reconstruction. As described in Fig. 1c and Extended Data Fig. 4a, the low-resolution input stack is rotated by three orientations, anisotropically super-resolved by the trained VSI-SR model, and combined via the generalized Wiener filter to produce 2D isotropic SR reconstruction.

**C. 3D isotropic super-resolution reconstruction**

The final step is to fuse the two views of 2D SR image stacks to achieve 3D isotropic SR reconstruction, which contains the following three steps:

**C.1 View rotation:** To facilitate dual-view splitting, we first rescale the stack in Z-axis to ensure an isotropic sampling rate of the volume, and then rotate the deskewed stack to transform the sample from the detection objective coordinate XYZ to the sample coordinate X'Y'Z', where the X'Y' plane is parallel to the coverslip plane. The rotation angle is set as the designed angle between the detection objective and the coverslip, which is 30.8° in our setup. Ideally, after the rotation, the two views of the sample should be in perfect symmetry vertically. However, the practical system errors such as the slight illumination off-axis and reflective coverslip tilting will cause the misalignment of the two views.

**C.2 Dual-view splitting and registration:** After the initial rotation in C.1, the two views are roughly aligned. To precisely split and align the two imaging views, we adopt an iterative refinement procedure. In our system, we assume a rigid alignment, taking account of the tilting in both X- and Y-axis as well as the splitting position in Z-axis. The tilting angles along X- and Y-axis are estimated alternatively. For tilting angle X, we calculate the MIP in YZ plane and estimate both tilting angles and splitting planes to optimize the alignment of two views. Specifically, we first loop over angles within a small search range (e.g., -1.5°~1.5°) by small steps (e.g., 0.1°); and for each candidate tilting angle, we loop over the splitting Z-position and calculate the cross-view correlation. The paired choice of tilting angle X and splitting position Z that maximizes the cross-view correlation gives the optimal estimation of view splitting and registration. The same procedure is applied to the MIP in XZ plane to estimate the tilting angle Y. The whole process can be optionally repeated for 1-2 rounds to better refine the angle and splitting position estimation.

**C.3 Dual-view fusion with RL-DFN:** Finally, the registered dual-view stacks are fused via the proposed RL-DFN, as described in Fig. 4b, Extended Data Fig. 7a and Supplementary Fig. 6.

### Supplementary Figures

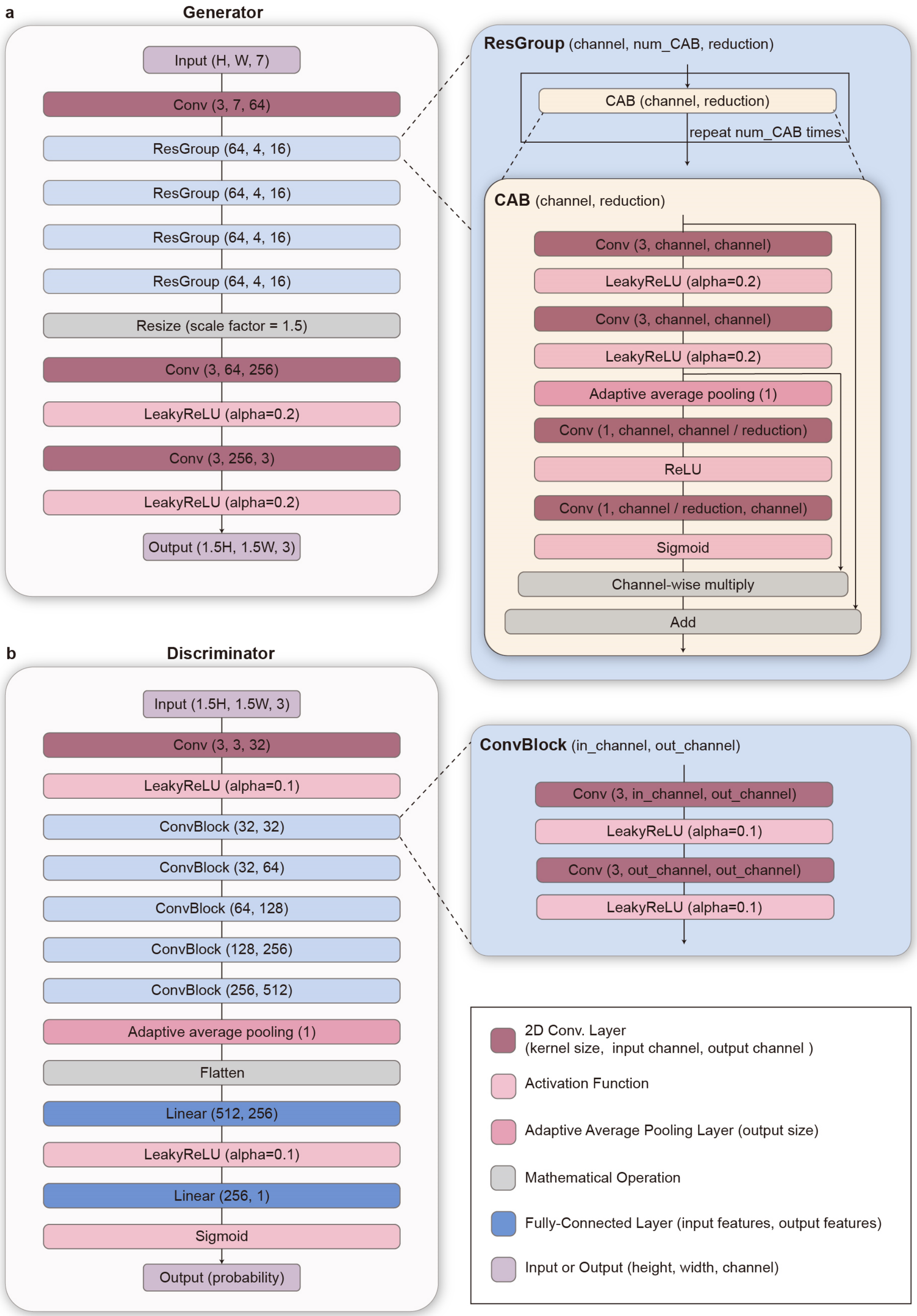

**Supplementary Fig. 1 | Network architecture of the VSI-SR model. a,** Generator of the VSI-SR model. **b**, Discriminator of the VSI-SR model.

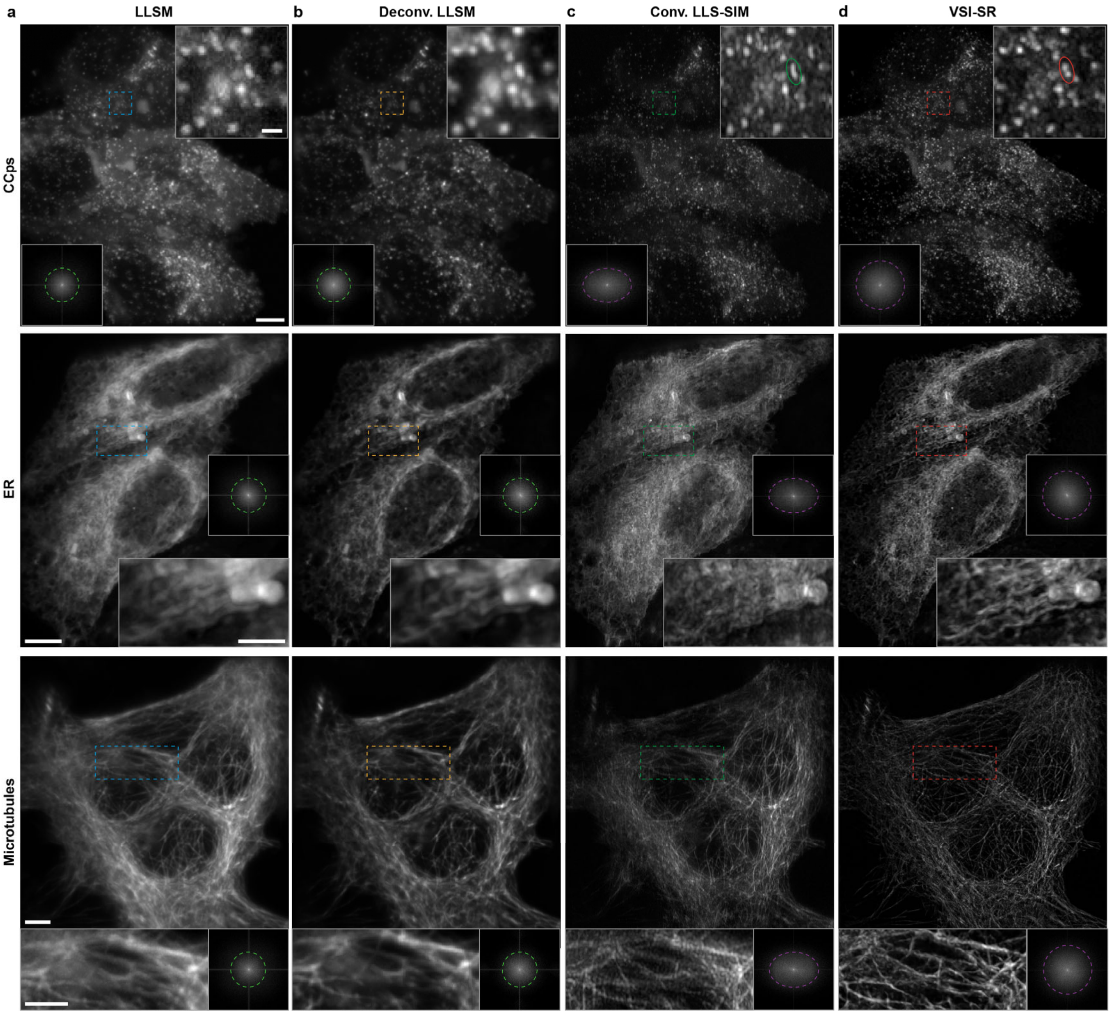

**Supplementary Fig. 2 | Laterally isotropic SR reconstruction for LLSM images of different organelles by VSI-SR methods.** Maximum intensity projections (MIPs) of CCPs, ER, and microtubules imaged by LLSM (a), and post-processed by RL deconvolution (b), conventional LLS-SIM (c), VSI-SR model-based laterally isotropic SR reconstruction (d). Logarithmic Fourier spectrums are shown in the bottom right corner of each full-FOV image to indicate the frequency component extension by the methods compared. Scale bar, 5 μm (full FOV), and 2 μm (magnified insets).

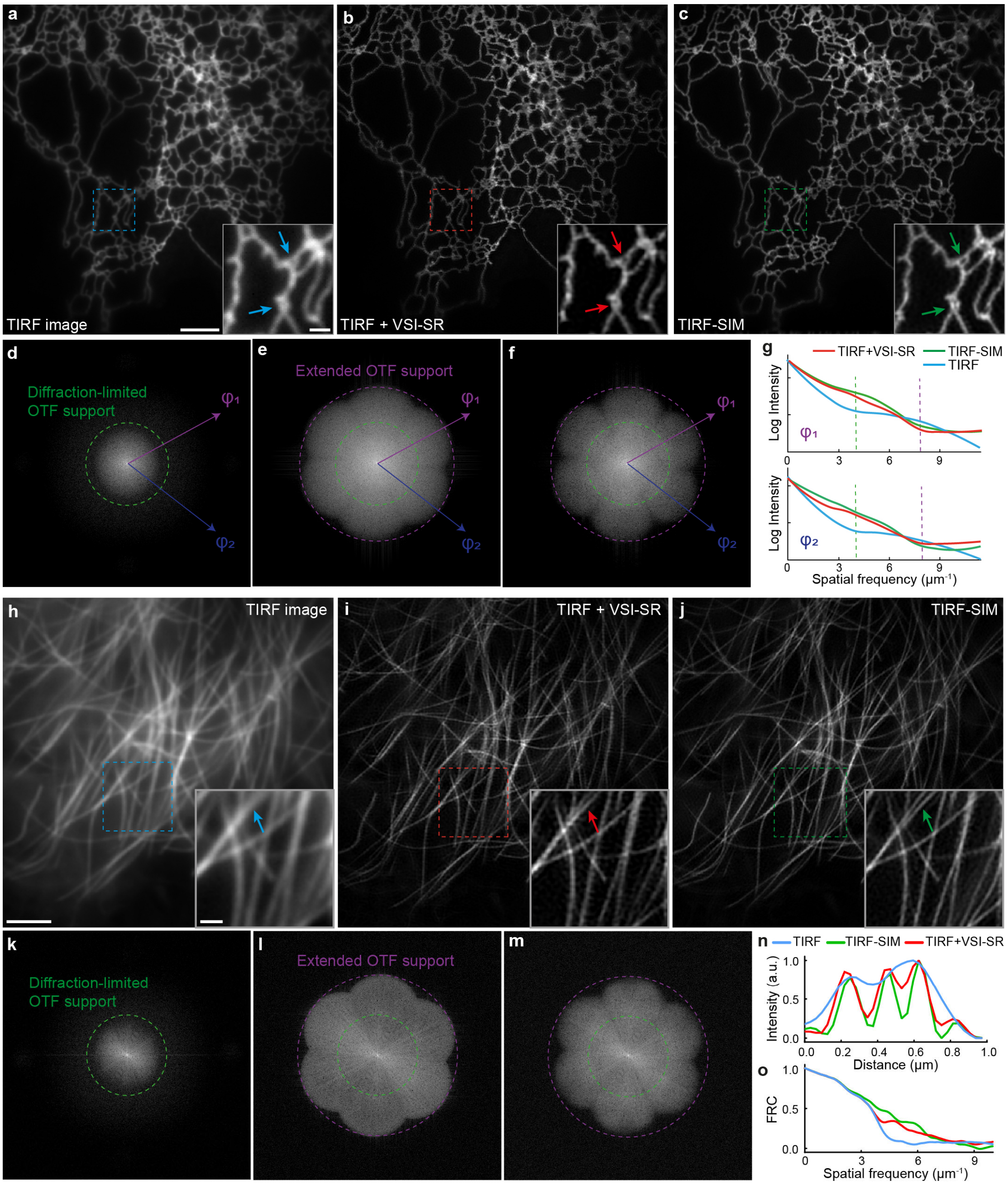

**Supplementary Fig. 3 | Characterization of 2D isotropic SR recovery for TIRF images via VSI-SR methods.** **a**, TIRF image of ER using uniform wide-field illumination; **b**, Laterally isotropic SR image reconstructed by VSI-SR from the TIRF image shown in a. **c**, TIRF-SIM image of the same region with a. **d-f**, Corresponding Fourier spectrums of a-c. **g**, Line profiles in log-scale along orientations $\phi_1$ and $\phi_2$ as drawn with the purple and blue lines in d-f. **h-m**, Images (h-j) and Fourier spectrums (k-m) of microtubules acquired using TIRF, VSI-SR model, and TIRF-SIM with three-orientation structured illumination. **n**, Line profiles of two closely located microtubules indicated by the arrow in h-j. **o**, Fourier ring correlation (FRC) analysis of the images shown in h-j. Scale-bar, 5 μm (a-c, h-j), 1 μm (zoom-in regions of a-c), 2 μm (zoom-in regions of h-j).

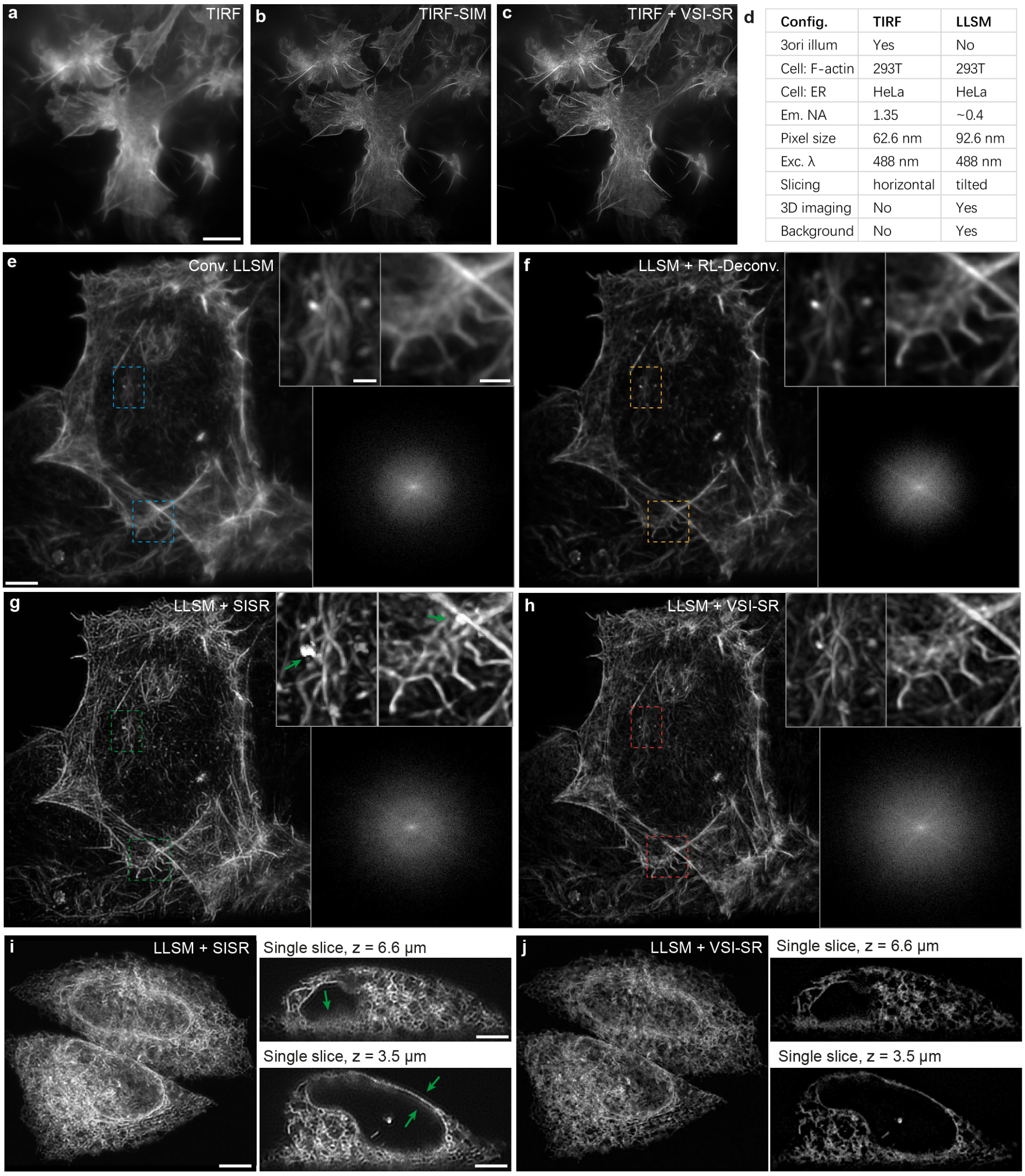

| Config. | TIRF | LLSM |
|---|---|---|
| 3ori illum | Yes | No |
| Cell: F-actin | 293T | 293T |
| Cell: ER | HeLa | HeLa |
| Em. NA | 1.35 | ~0.4 |
| Pixel size | 62.6 nm | 92.6 nm |
| Exc. λ | 488 nm | 488 nm |
| Slicing | horizontal | tilted |
| 3D imaging | No | Yes |
| Background | No | Yes |

**Supplementary Fig. 4 | Comparison of 2D isotropic SR reconstruction between VSI-SR and cross-modality learning. a-c**, Representative F-actin images obtained by TIRF microscopy (a), TIRF-SIM (b), and VSI-SR reconstructed from a (c). **d**, Comparisons between TIRF-SIM and LLSM imaging modalities in terms of 3D imaging capability, imaging geometry, etc. **e-h**, F-actin images (MIP) acquired by conventional LLSM (e) and processed by RL deconvolution (f), single image super-resolution (SISR) model (g), i.e., DFCAN[11] trained with TIRF/3-orientation-TIRF-SIM image pairs and applied to LLSM inputs in a cross-modality manner, and VSI-SR model (h) trained with LLSM/LLS-SIM image pairs. The magnified insets reveal reconstruction artifacts of the cross-modality approach (see the bright-spots highlighted by green arrows in g). **i,j**, SR images of ER reconstructed using the cross-modality SISR model and the proposed VSI-SR scheme. Obvious background fuzziness and ring artifacts occur for the cross-modality results (highlighted by green arrows) because of the domain shift problem. Scale bar, 5 μm (full-FOV images of a-j), 3 μm (magnified insets of a-j).

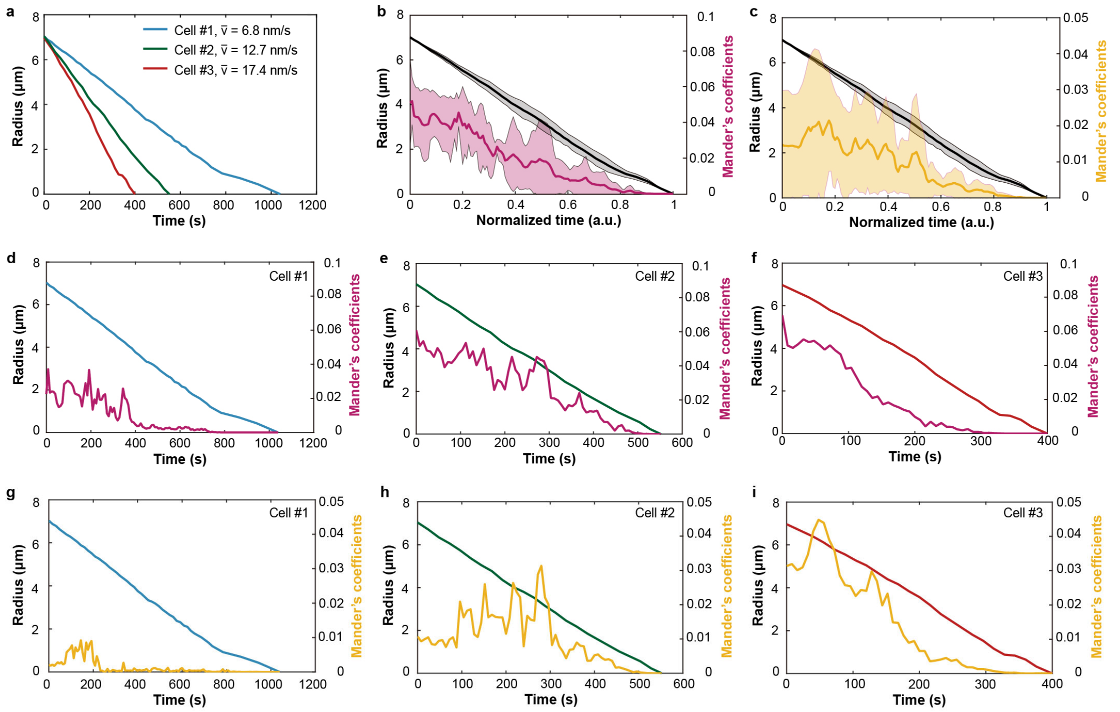

**Supplementary Fig. 5 | Interaction quantification between the contractile ring and other organelles during mitosis. a**, Plots of the contractile ring (CR) radius along with time for three mitotic HeLa cells. **b**,**c**, Plots of the CR radius (gray) and Mander's overlap coefficients (MOCs) between CR and ER (b, magenta) or lysosomes (c, yellow) calculated in the cross-sectional area of the CR over the normalized time course of mitosis. **d**-**f**, Quantification of CR radius and MOCs between CR and ER over the time course of mitosis for cell #1 (d), cell #2 (e), and cell #3 (f), respectively. **g**-**i**, Quantification of CR radius and MOCs between CR and lysosomes over the time course of mitosis for cell #1 (g), cell #2 (h), and cell #3 (i), respectively.

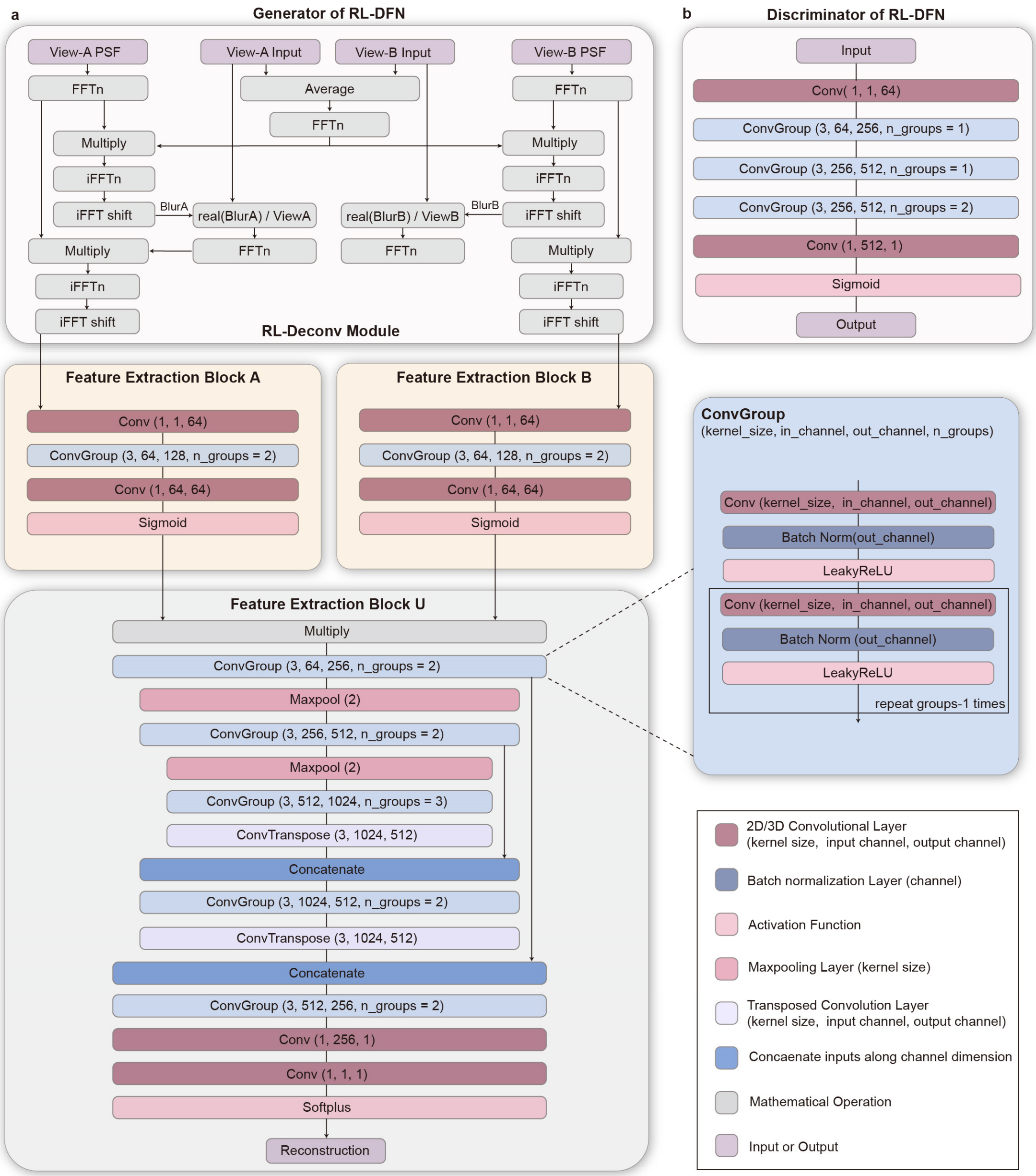

**Supplementary Fig. 6 | Network architecture of RL-DFN. a**, Generator of the RL-DFN model. **b**, Discriminator of the RL-DFN model.

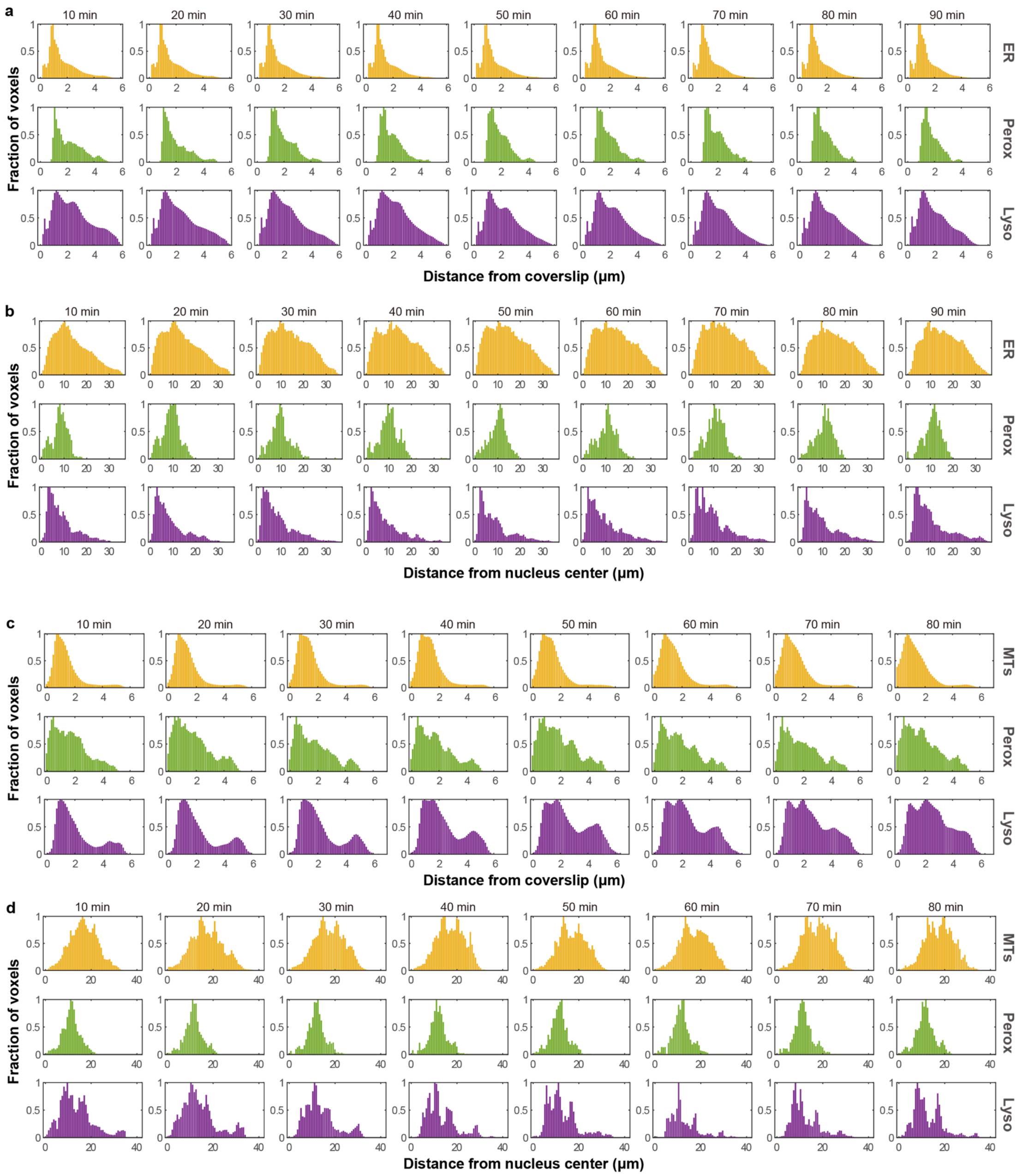

**Supplementary Fig. 7 | Voxel distributions of organelles in axial and lateral dimensions. a**,**b**, Distributions of organelles in the axial (a) and lateral (b) dimensions of a Cos-7 cell labelling endoplasmic reticulum (ER), peroxisomes (Perox), and lysosomes (Lyso). **c**,**d**, Distributions of organelles in the axial (c) and lateral (b) dimensions of another Cos-7 cell labelling microtubules (MTs), Perox, and Lyso. To display the distribution progression along with time, averaged results for each 10-minute duration are shown.

### Supplementary Tables

#### Supplementary Table 1 | Imaging conditions of live-cell experiments

| Data | Imaging method (Acquisition mode) | Sample | Label | Excitation NA | Excitation λ (nm) | Exposure time per raw image (ms) | Total acquisition time (sec) | Volume size of raw data (Width×Height ×Z-slice×Channel) | Cycle time (Acquisition + resting time) (sec) | Time points |
|---|---|---|---|---|---|---|---|---|---|---|
| Fig. 3a-d<br>Supplementary Video 2 | Meta-LLS-VSIM (slit-scan mode) | HeLa | Lifeact-mEmerald<br>KDEL-mCherry<br>Lamp1-Halo | 0.11 | 488<br>560<br>642 | 6 | ~5.1 | 600×600×181×3 | 8 | 357 |
| Fig. 3e<br>Supplementary Video 3 | Meta-LLS-VSIM (slit-scan mode) | Mouse embryo | TOMM20-mEmerald | 0.069 | 488 | 30 | ~9.1 | 1024×1024×301×1 | 30 | 100 |
| Fig. 3f, g<br>Supplementary Video 4 | Meta-LLS-VSIM (slit-scan mode) | Pollen tube | Lifeact-GFP | 0.11 | 488 | 1 | ~0.24 | 256×512×101×1 | 0.24 | 1000 |
| Fig. 3h-k<br>Supplementary Video 5 | Meta-LLS-VSIM (slit-scan mode) | *C. elegans* embryo | wyEx51119, jcIs1<br>qxIs257 | 0.11 | 488<br>560 | 6 | ~5 | 608×608×201×2 | 60 | 67 |
| Fig. 5a-f<br>Supplementary Video 6 | Meta-rLLS-VSIM (sheet-scan mode) | Cos-7 | GFP-SKL<br>ER-mCherry<br>Lyso-Halo | 0.35, 0.14 | 488<br>560<br>642 | 6 | ~6.2 | 320×1024×301×3 | 12 | 400 |
| Fig. 5k, l<br>Supplementary Video 8 | Meta-rLLS-VSIM (sheet-scan mode) | Cos-7 | 3×mEmerald-Ensconsin<br>mCherry-SKL<br>Lamp1-Halo | 0.35, 0.14 | 488<br>560<br>642 | 6 | ~7.8 | 800×600×181×3 | 8 | 721 |
| Extended Data Fig. 10<br>Supplementary Video 7 | Meta-rLLS-VSIM (sheet-scan mode) | Cos-7 | GFP-SKL<br>ER-mCherry<br>Lyso-Halo | 0.35, 0.14 | 488<br>560<br>642 | 6 | ~5.2 | 384×600×221×3 | 8 | 812 |

**Supplementary Table 2 | Information of dataset used for training the Meta-VSI-SR model.**

| Cell type | Structure | Label | Number of stack pairs | Raw data size before deskew (Width×Height×Z-slice×Phase) | LLS-SIM data size after deskew (Width×Height×Z-slice) |
|---|---|---|---|---|---|
| HeLa | Outermost granular component | mEmerald-NPM1 | 104 | 256×512×121×3 | 1042×768×121 |
| HeLa | Chromosome | mCherry-H2B | 96 | 256×512×151×3 | 1205×768×151 |
| HeLa | Innermost fibrillar center | Halo-RPA49 | 46 | 256×512×151×3 | 1205×768×151 |
| Cos-7 | ER (in adherent cells) | Calnexin-mEmerald | 50 | 512×512×151×3 | 1589×768×151 |
| HeLa | Fibrillarin | FBL-mEmerald | 32 | 256×512×121×3 | 1042×768×121 |
| HeLa | Lysosome | Lamp1-Halo | 68 | 512×512×151×3 | 1589×768×151 |
| HeLa | Microtubule | 3×mEmerald-Ensconsin | 34 | 256×512×121×3 | 1042×768×121 |
| Pollen tube | F-actin (in pollen tube) | Lifeact-GFP | 46 | 512×512×201×3 | 1027×768×201 |
| Cos-7 | Inner mitochondrial membrane | PKMR | 38 | 256×512×121×3 | 1042×768×121 |
| HeLa | ER (metaphase of mitosis) | KDEL-mEmerald | 22 | 256×512×151×3 | 1205×768×151 |

### Captions for Supplementary Videos

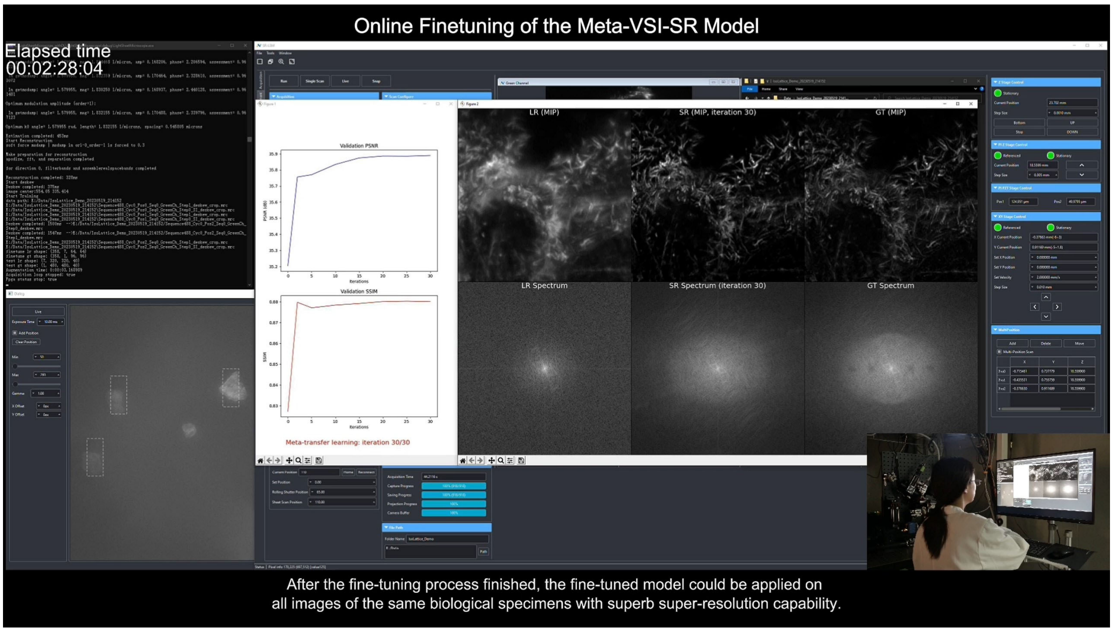

**Supplementary Video 1 | Demonstration of online finetuning of the Meta-VSI-SR model to achieve quick adaptation to a new biological structure.** After operating on the developed GUI to select three ROIs of the target biological sample, an automatic streamline including data acquisition, LLS-SIM SR image reconstruction and finetuning of the Meta-VSI-SR was produced. As the finetuning goes on, the reconstruction performance of the meta-model was substantially improved within few iterations. All of above operations typically take less than 2.5 minutes as shown in this video.

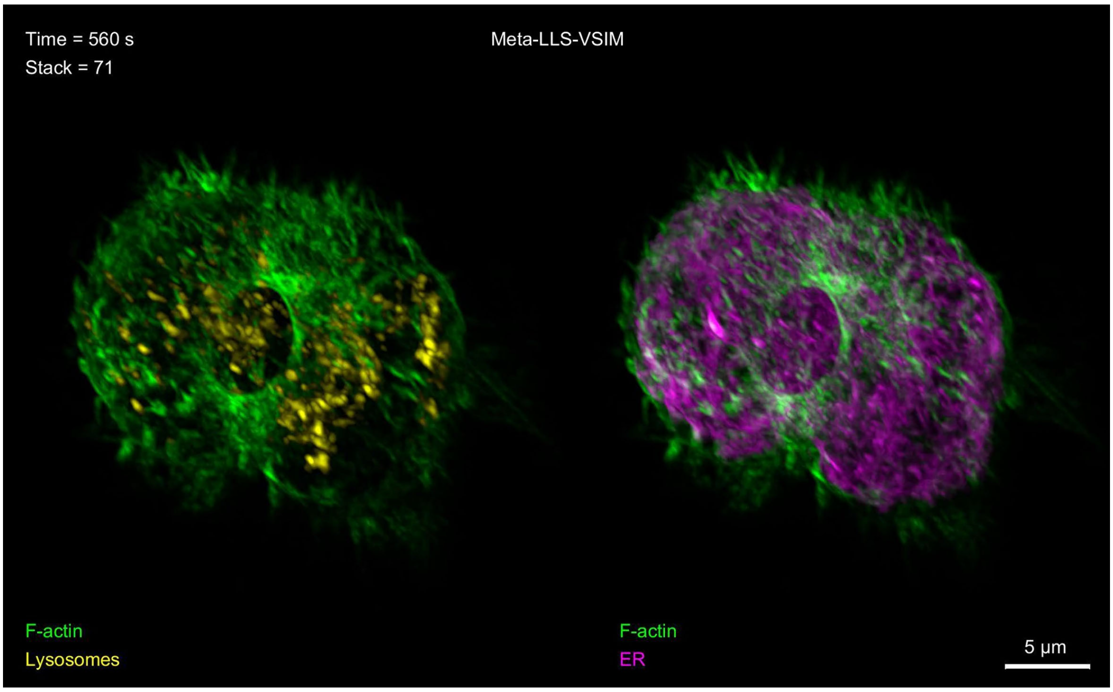

**Supplementary Video 2 | Entire process of contractile ring formation, contraction and disassembly during mitosis imaged by Meta-LLS-VSIM.** Three-color Meta-LLS-VSIM imaging for 357 timepoints at 8-second volume intervals of a Hela cell stably expressing Lifeact-mEmerald (F-actin in green), KDEL-mCherry (ER in magenta), and Lamp1-Halo (lysosomes in yellow).

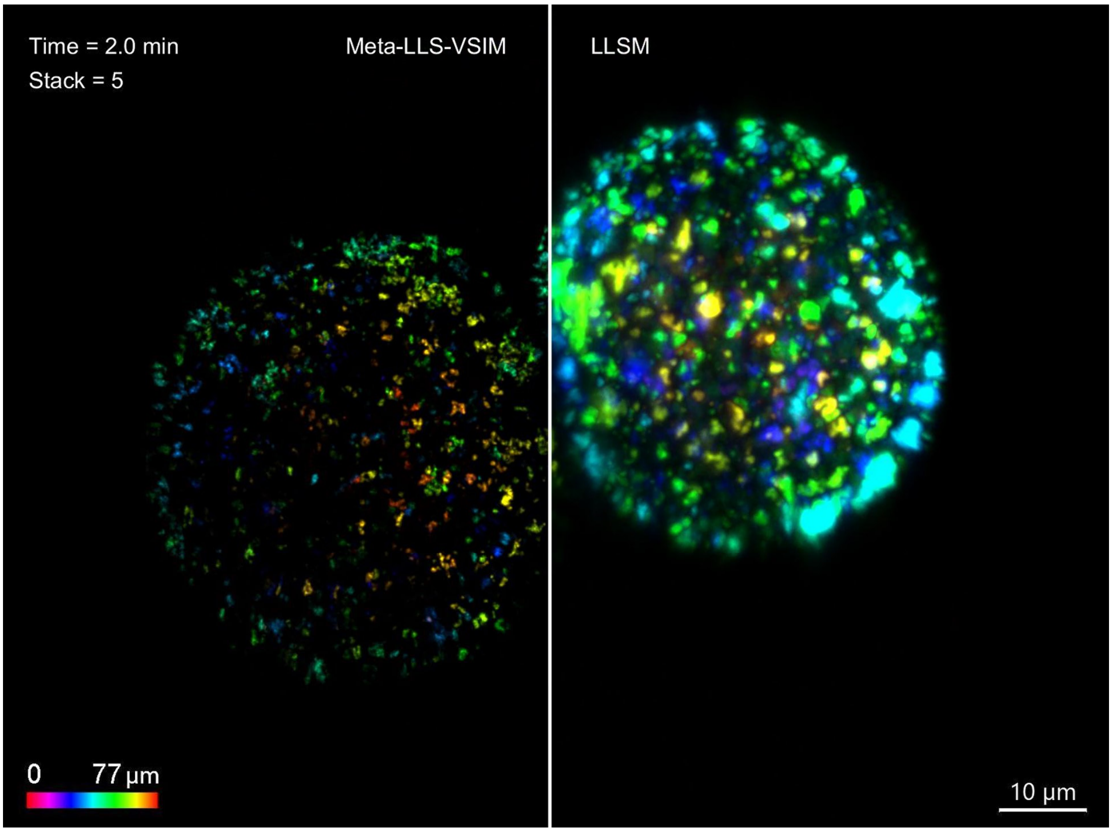

**Supplementary Video 3 | The dynamics of mitochondria in a mouse embryo imaged by Meta-LLS-VSIM.** Meta-LLS-VSIM imaging in rolling-shutter confocal slit-scan mode of a developing mouse embryo with mitochondria labelled by TOMM20-mEmerald across a thick area of 95×95×96 $\mu m^3$ for 100 timepoints at 30-second intervals. The trajectory of mitochondria is delineated.

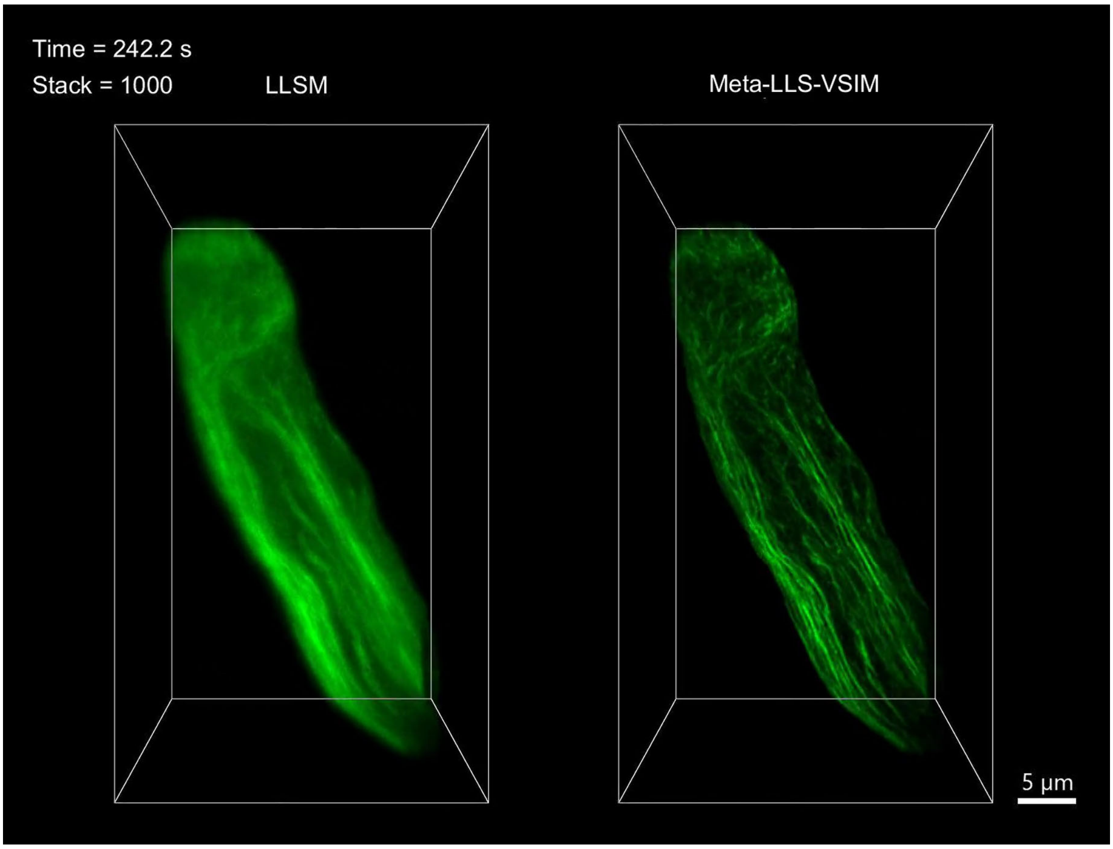

**Supplementary Video 4 | The spatiotemporal dynamics of cytoskeleton during growing of pollen tubes.** LLSM and Meta-LLS-VSIM recording a growing pollen tube labelled with Lifeact-GFP at a high speed of 4.125 Hz for 1,000 timepoints.

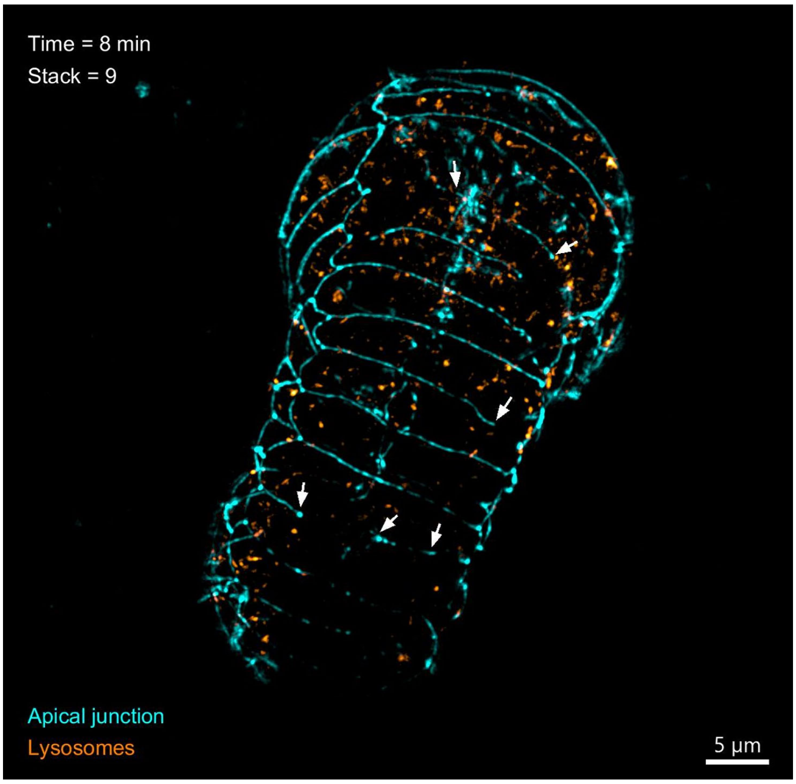

**Supplementary Video 5 | Plasma membrane fusion process in *C. elegans* embryo development by two-color Meta-LLS-VSIM.** Two-color volumetric recording of the seam cell fusion process in a *C. elegans* embryo with apical junction (cyan) and lysosomes (orange) labelled by Meta-LLS-VSIM with a time duration of 66 minutes at 1-minute intervals.

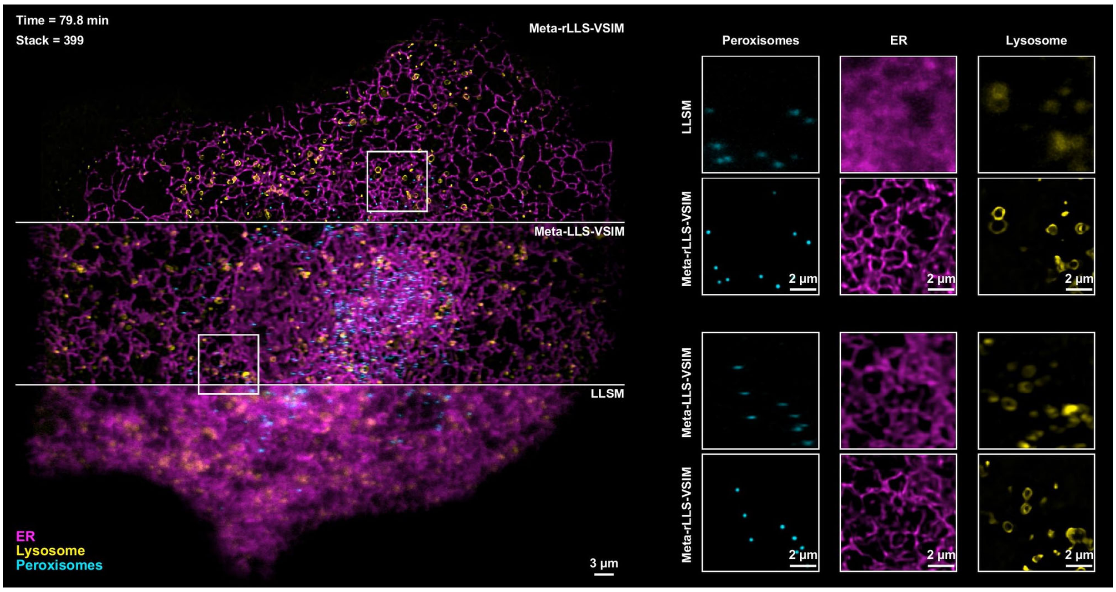

**Supplementary Video 6 | Long-term 4D SR imaging of the whole cell by Meta-rLLS-VSIM.** Three-color, long-term, near-isotropic SR 4D imaging of a Cos-7 cell transferred with GFP-SKL (peroxisomes in cyan), ER-mCherry (ER in magenta), and Lyso-Halo (lysosomes in yellow) for 400 timepoints at 12-second intervals by Meta-rLLS-VSIM.

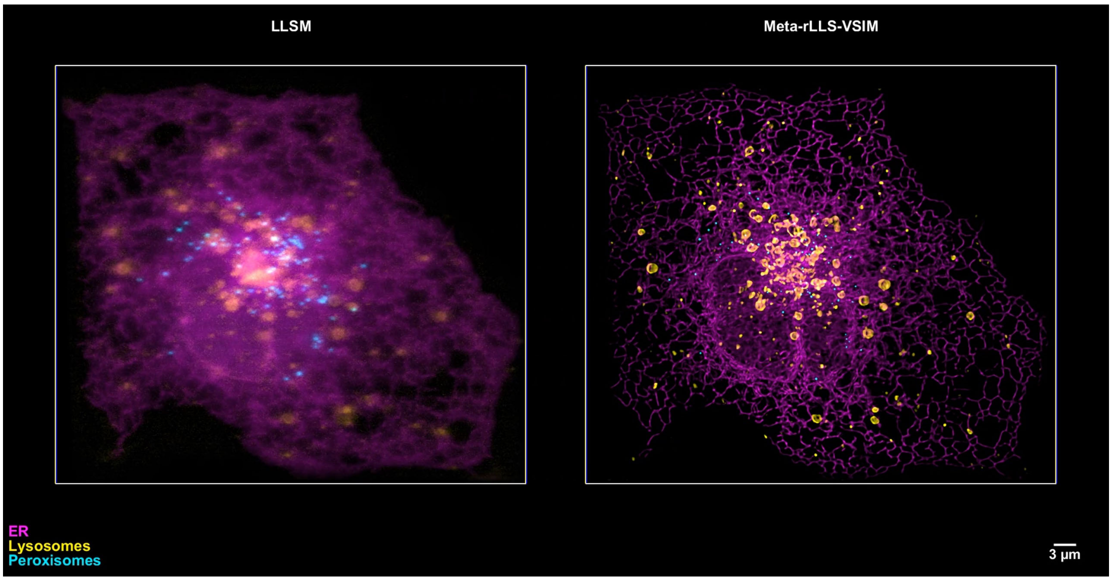

**Supplementary Video 7 | Illustration of near-isotropic SR imaging by dual-view fusion in Meta-rLLS-VSIM.** Near-isotropic 3D SR reconstruction from dual-view volumes by Meta-rLLS-VSIM of a Cos-7 cell transferred with GFP-SKL (peroxisomes in cyan), ER-mCherry (ER in magenta), and Lyso-Halo (lysosomes in yellow) for 721 timepoints at a speed of 8 seconds per three-color whole-cell volume.

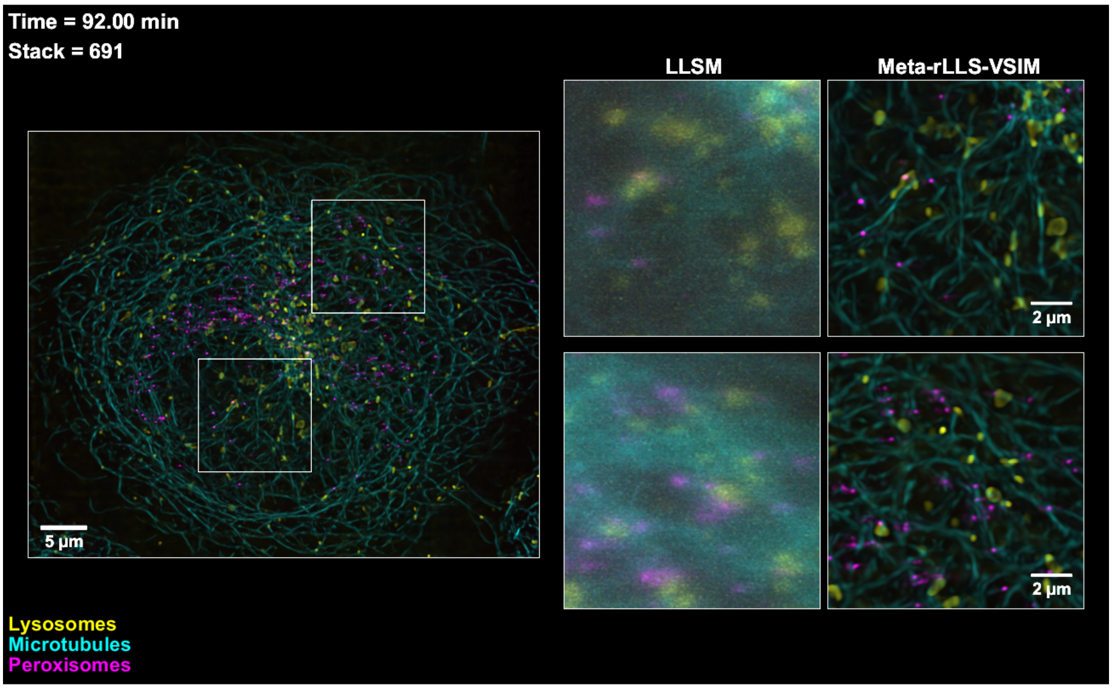

**Supplementary Video 8 | Long-term 4D subcellular imaging of multi-organelles and cytoskeleton.** Meta-rLLS-VSIM records the spatiotemporal coordination and interactions between multiple organelles and cytoskeleton in a Cos-7 cell labelled by 3×mEmerald-Ensconsin (microtubules in cyan), mCherry-SKL (peroxisomes in magenta), and Lamp1-Halo (lysosomes in yellow) for 814 timepoints at a speed of 8 seconds per three-color cell volume.

5 Goodfellow, I. *et al.* in *Advances in neural information processing systems.* 2672-2680.

6 Fuoli, D., Van Gool, L. & Timofte, R. in *Proceedings of the IEEE/CVF International Conference on Computer Vision.* 2360-2369.

7 Ledig, C. *et al.* in *Proceedings of the IEEE conference on computer vision and pattern recognition.* 4681-4690.

8 Guo, Y. *et al.* Visualizing Intracellular Organelle and Cytoskeletal Interactions at Nanoscale Resolution on Millisecond Timescales. *Cell* **175**, 1430-1442 e1417 (2018). https://doi.org/10.1016/j.cell.2018.09.057

9 Li, D. *et al.* Extended-resolution structured illumination imaging of endocytic and cytoskeletal dynamics. *Science* **349**, aab3500 (2015). https://doi.org/10.1126/science.aab3500

10 Nieuwenhuizen, R. P. *et al.* Measuring image resolution in optical nanoscopy. *Nature methods* **10**, 557-562 (2013).

11 Qiao, C. *et al.* Evaluation and development of deep neural networks for image super-resolution in optical microscopy. *Nature Methods* **18**, 194-202 (2021). https://doi.org/10.1038/s41592-020-01048-5

12 Wang, H. *et al.* Deep learning enables cross-modality super-resolution in fluorescence microscopy. *Nature Methods* **16**, 103-110 (2019). https://doi.org/10.1038/s41592-018-0239-0

13 Chen, B. C. *et al.* Lattice light-sheet microscopy: imaging molecules to embryos at high spatiotemporal resolution. *Science* **346**, 1257998 (2014). https://doi.org/10.1126/science.1257998

14 Sung, F. *et al.* in *Proceedings of the IEEE conference on computer vision and pattern recognition.* 1199-1208.

15 Snell, J., Swersky, K. & Zemel, R. Prototypical networks for few-shot learning. *Advances in neural information processing systems* **30** (2017).

16 Vinyals, O., Blundell, C., Lillicrap, T. & Wierstra, D. Matching networks for one shot learning. *Advances in neural information processing systems* **29** (2016).

17 Oreshkin, B., Rodríguez López, P. & Lacoste, A. Tadam: Task dependent adaptive metric for improved few-shot learning. *Advances in neural information processing systems* **31** (2018).

18 Mishra, N., Rohaninejad, M., Chen, X. & Abbeel, P. in *International Conference on Learning Representations.*

19 Santoro, A., Bartunov, S., Botvinick, M., Wierstra, D. & Lillicrap, T. in *International conference on machine learning.* 1842-1850 (PMLR).

20 Lee, Y. & Choi, S. in *Proceedings of the 35th International Conference on Machine Learning* Vol.

80 (eds Dy Jennifer & Krause Andreas) 2927--2936 (PMLR, Proceedings of Machine Learning Research, 2018).

21 Grant, E., Finn, C., Levine, S., Darrell, T. & Griffiths, T. in *International Conference on Learning Representations.*

22 Finn, C. & Levine, S. in *International Conference on Learning Representations.*

23 Finn, C., Abbeel, P. & Levine, S. in *International Conference on Machine Learning.* 1126-1135 (PMLR).

24 Antoniou, A., Edwards, H. & Storkey, A. How to train your MAML. *arXiv preprint arXiv:1810.09502* (2018).

25 Nichol, A., Achiam, J. & Schulman, J. On first-order meta-learning algorithms. *arXiv preprint arXiv:1803.02999* (2018).

26 Mirza, M. & Osindero, S. Conditional Generative Adversarial Nets. arXiv:1411.1784 (2014). <https://ui.adsabs.harvard.edu/abs/2014arXiv1411.1784M>.

27 Zhu, J.-Y., Park, T., Isola, P. & Efros, A. A. in *Proceedings of the IEEE international conference on computer vision.* 2223-2232.

28 Preibisch, S. *et al.* Efficient Bayesian-based multiview deconvolution. *Nat Methods* **11**, 645-648 (2014). https://doi.org/10.1038/nmeth.2929

29 Li, Y. *et al.* Incorporating the image formation process into deep learning improves network performance. *Nature Methods* **19**, 1427-1437 (2022). https://doi.org/10.1038/s41592-022-01652-7

30 Guo, M. *et al.* Rapid image deconvolution and multiview fusion for optical microscopy. *Nature Biotechnology* **38**, 1337-1346 (2020). https://doi.org/10.1038/s41587-020-0560-x

31 Wu, Y. *et al.* Multiview confocal super-resolution microscopy. *Nature* **600**, 279-284 (2021). https://doi.org/10.1038/s41586-021-04110-0

32 Li, X. *et al.* Three-dimensional structured illumination microscopy with enhanced axial resolution. *Nature Biotechnology* (2023). https://doi.org/10.1038/s41587-022-01651-1

33 Park, H. *et al.* Deep learning enables reference-free isotropic super-resolution for volumetric fluorescence microscopy. *Nature Communications* **13**, 3297 (2022). https://doi.org/10.1038/s41467-022-30949-6

34 Ning, K. *et al.* Deep self-learning enables fast, high-fidelity isotropic resolution restoration for volumetric fluorescence microscopy. *Light: Science & Applications* **12**, 204 (2023). https://doi.org/10.1038/s41377-023-01230-2
